# Personality and local brain structure: their shared genetic basis and reproducibility

**DOI:** 10.1101/645945

**Authors:** Sofie L. Valk, Felix Hoffstaedter, Julia A. Camilleri, Peter Kochunov, B.T. Thomas Yeo, Simon B. Eickhoff

## Abstract

Local variation in cortical architecture is highly heritable and distinct genes are associated with specific cortical regions. Total surface area has been shown to be genetically correlated with complex cognitive capacities, suggesting cortical brain structure is a viable endophenotype linking genes to behavior. However, to what extend local brain structure has a genetic association with cognitive and emotional functioning is incompletely understood. Here, we study the genetic correlation between personality traits and local cortical structure in a large-scale twin sample (Human Connectome Project, n=1106, 22-37y). We found a genetic overlap between personality traits and local cortical structure in 10 of 17 observed phenotypic associations in predominantly frontal cortices. To evaluate the robustness of observed personality-brain associations we studied two independent age-matched samples (GSP: n=926, age=19-35y, eNKI: n=210, age: 19-39y). We observed anecdotal to moderate evidence for a successful replication of the negative association between surface area in medial prefrontal cortex and Neuroticism in both samples. Quantitative functional decoding indicated this region is implicated in emotional and socio-cognitive functional processes. In sum, our observations suggest that associations between local brain structure and personality are, in part, under genetic control. However, associations are weak and only the relation between frontal surface area and Neuroticism was consistently observed across three independent samples of young adults.

## Introduction

The local macro-anatomical structure of the cerebral cortex is largely heritable, and has a highly polygenetic architecture (Grasby et al., 2020; Panizzon et al., 2009; Strike et al., 2019; Winkler et al., 2010). Recently, it has been shown that common genetic variants that influence surface area also affect various behavioral traits, suggesting that brain structure is an essential endophenotype linking genes and behavior (Grasby et al., 2020). However, to what extend the correlation between local cortical structure on the one hand and cognitive and emotional functioning on the other is driven by shared genetic factors is incompletely understood.

One of the most broadly used summaries of an individual’s characteristic patterns of behavior, thought, and emotions is personality (Funder, 2001). Behavioral science establishes personality structure by parcellating the individual variability in goals, cognition, and emotion into independent components (Mischel, 2004). A widely used personality taxonomy is the Big Five Personality inventory (John and Srivastava, 1999; McCrae and Costa, 1997; Saucier and Srivastava, 2015). The Five-factor personality structure derives five orthogonal dimensions or traits of Agreeableness, Conscientiousness, Extraversion, Neuroticism, and Openness (John et al., 2008; Saucier and Srivastava, 2015). Personality traits have been related to the quality of social relationships (Asendorpf and Wilpers, 1998), job performance (Rothmann and Coetzer, 2003), risk for mental disorders (Miller et al., 2001; Trull, 2013), general health and wellbeing, and reproductive success (Alvergne et al., 2010; Strickhouser et al., 2017).

Personality has both stable and malleable features (Damian et al., 2019; Harris et al., 2016; Penke and Jokela, 2016) and has been found heritable with approximately 40% of the variance attributable to additive genetic factors (Bouchard, 1994; Bouchard and Loehlin, 2001; Bouchard and McGue, 2003; Jang et al., 1996; Loehlin et al., 1998; Vukasovic and Bratko, 2015). Evolutionary causes for variability in personality traits have been suggested to be due to balancing selection, where selection pressures in different directions affect the same traits enabling adaptation to changing environmental demands (Penke and Jokela, 2016). Indeed, genome-wide association studies (GWAS) have reported a large number of genetic variants associated with personality traits with each contributing to the heritability of personality (Consortium, 2015; de Moor et al., 2012; Lo et al., 2017; van den Berg et al., 2016; Verweij et al., 2012).

The biological basis of personality in humans has also been studied in relation to macroscale brain structure and function using magnetic resonance imaging (MRI) (Bjornebekk et al., 2013; DeYoung et al., 2010; Dubois et al., 2018; Ferschmann et al., 2018; Kong et al., 2019; Nostro et al., 2017; Riccelli et al., 2017; Wu et al., 2019). Various studies have reported a phenotypic relationship between local brain structure and personality traits (Bjornebekk et al., 2013; DeYoung et al., 2010; Gray et al., 2019; Nostro et al., 2017; Riccelli et al., 2017). Using the Human Connectome Project, young adult (HCP) sample, including monozygotic and dizygotic twins, Owens and colleagues (Owens et al., 2019) report significant phenotypic relationships between personality traits and various markers of cortical structure. For example, Owens and colleagues observed associations between Agreeableness, Conscientiousness, Neuroticism, and Openness with morphometry in prefrontal areas. However findings on the relationship between personality traits and local brain structure have been inconsistent, for instance, Avinun and colleagues failed to observe significant relations between personality and various markers of brain structure using the largest sample to date (Avinun et al., bioarXiv). In line with this report, Kharabian et al. have recently shown that, in general, relationships between local brain structure and psychometric variables are not robust and highly dependent on sample and effect size (Kharabian Masouleh et al., 2020; Kharabian Masouleh et al., 2019). At the same time, it has recently been shown that traits such as neuroticism, general cognitive function, educational attainment, and depressive symptoms show a genetic correlation with total surface area, suggesting brain structure is a key phenotype reflecting individual differences in behavior (Grasby et al., 2020).

Taken together contemporary theory suggests that (a) individual variation in both local brain structure and personality can be, in part, attributed to genetic effects (b) brain structure is a viable endophenotype linking genes and behavior (c) personality relates to macro-scale brain structure and function, but local relationships are weak and vary as a function of sample and effect size. However, whether regional brain structure and personality have a shared genetic basis remains unclear. To answer our research question, we studied the relationship between the Big Five personality traits and local cortical thickness and surface area. We captured variations in brain morphometry using an atlas-based approach, dividing the cortex in 200 functionally-defined parcels (Eickhoff et al., 2018; Schaefer et al., 2018). We studied three independent samples of young adults, the Human Connectome Project Young Adult (HCP, n=1106), Brain Genomics superstructure project (GSP, n=926) and enhanced Nathan Kline Institute dataset (eNKI, n=210). The HCP sample is unique in that it provided us with high quality neuroimaging and personality trait (NEO-FFI) data in a large number of twins, siblings, and unrelated individuals, enabling us to compute genetic correlation between personality and local brain structure. Analysis of heritability and genetic correlation was performed using maximum likelihood variance-decomposition methods using Sequential Oligogenic Linkage Analysis Routines (www.solar-eclipse-genetics.org; Solar Eclipse 8.4.0.). Second, to assess whether observed associations between personality and local brain structure in the HCP sample reflect generalizable relationships between personality and local brain structure, we selected two samples (Brain Genomics superstructure project – GSP (n=926) and enhanced Nathan Kline Institute dataset – eNKI (n=210)) of unrelated individuals between 20 and 40 years of age in which we studied whether personality-brain relationships observed in the HCP sample would replicate in two independent samples. Last, we performed functional decoding to further evaluate the functional mechanisms underlying brain regions robustly associated with personality.

## Materials and methods

### HCP sample

#### Participants and study design

For our analysis we used the publicly available data from the Human Connectome Project S1200 release (HCP; http://www.humanconnectome.org/), which comprised data from 1206 individuals (656 females), 298 MZ twins, 188 DZ twins, and 720 singletons, with mean age 28.8 years (SD = 3.7, min-max = 22–37). We included individuals for whom the scans and data had been released (humanconnectome.org) after passing the HCP quality control and assurance standards (Marcus et al., 2013). The full set of inclusion and exclusion criteria are described elsewhere (Glasser et al., 2013; Van Essen et al., 2013). In short, the primary participant pool comes from healthy individuals born in Missouri to families that include twins, based on data from the Missouri Department of Health and Senior Services Bureau of Vital Records. Additional recruiting efforts were used to ensure participants broadly reflect ethnic and racial composition of the U.S. population. Healthy is broadly defined, in order to gain a sample generally representative of the population at large. Sibships with individuals having severe neurodevelopmental disorders (e.g., autism), documented neuropsychiatric disorders (e.g. schizophrenia or depression) or neurologic disorders (e.g. Parkinson’s disease) are excluded, as well as individuals with diabetes or high blood pressure. Twins born prior 34 weeks of gestation and non-twins born prior 37 weeks of gestation are excluded as well. After removing individuals with missing structural imaging or behavioral data our sample consisted of 1106 individuals (including 286 MZ-twins and 170 DZ-twins) with mean age of 28.8 years (SD =3.7, min-max =22-37). See further Table 1.

**Table 1.** Behavioral characteristics of the HCP sample. Behavioral characteristics for gender, age, intelligence as well as the NEO-FFI scores in the HCP sample.

| Measure | n | mean±SD (min-max) |
| --- | --- | --- |
| *Males/Females* | 506/600 | - |
| *Age* | 1106 | 28.8±3.7 (22-36) |
| *Intelligence (Composite score)* | 1089 | 121.9±14.6 (84.6-153.4) |
| *Agreeableness* | 1106 | 33.5±5.8 (10-48) |
| *Conscientiousness* | 1106 | 34.5±5.9 (11-48) |
| *Extraversion* | 1106 | 30.7±6 (10-47) |
| *Neuroticism* | 1106 | 16.6±7.4 (0-43) |
| *Openness* | 1106 | 28.3±6.2 (10-47) |

#### Structural imaging processing

MRI protocols of the HCP are previously described (Glasser et al., 2013; Van Essen et al., 2013). The pipeline used to obtain the Freesurfer-segmentation is described in detail in a previous article (Glasser et al., 2013) and is recommended for the HCP-data. In short, the pre-processing steps included co-registration of T1 and T2 scans, B1 (bias field) correction, and segmentation and surface reconstruction to estimate cortical thickness. The HCP structural pipelines use Freesurfer 5.1 software (http://surfer.nmr.mgh.harvard.edu/) (Dale et al., 1999; Fischl, 2013; Fischl and Dale, 2000; Fischl et al., 1999) plus a series of customized steps that combine information from T_1_w as well as T_2_w scans for more accurate white and pial surfaces (Glasser et al., 2013). Next, the individual cortical thickness and surface area maps were standardized to fsaverage5 for further analysis. Segmentation maps were visually inspected (S.L.V.) for inaccuracies.

#### Five Factor Model of Personality

The Big Five personality traits were assessed using the NEO-Five-Factors-Inventory (NEO-FFI)(McCrae and Costa, 2004). The NEO-FFI is composed of a subset of 60-items extracted from the full-length 240-item NEO-PI-R. For each item, participants reported their level of agreement on a 5-point Likert scale, from strongly disagree to strongly agree. The NEO instruments have been previously validated in USA and several other countries (McCrae and Terracciano, 2005). See further **Table 1**.

As a proxy for IQ we used the NIH Toolbox Cognition (Weintraub et al., 2013), ‘total composite score’. The Cognitive Function Composite score is derived by averaging the normalized scores of each of the Fluid and Crystallized cognition measures, then deriving scale scores based on this new distribution. Higher scores indicate higher levels of cognitive functioning. Participant score is normed to those in the entire NIH Toolbox Normative Sample (18 and older), regardless of age or any other variable, where a score of 100 indicates performance that was at the national average and a score of 115 or 85, indicates performance 1 SD above or below the national average. See further **Table 1**.

### GSP sample

#### Participants and study design

To evaluate the cross-sample reproducibility of observations we additionally investigated the association between personality and local cortical brain structure in the Brain Genomics Superstruct Project (GSP) (Holmes et al., 2015). In short, between 2008 and 2012 young adults (ages 18 to 35) with normal or corrected-to-normal vision were recruited from the Boston community to participate in the GSP. The 1,570 individuals included in the data release (Holmes et al., 2015) were selected from a larger databased of individuals who participated in the ongoing GSP data collection initiative. Participants included well-educated individuals with relatively high IQs (many of the college age students are from local colleges). Participants provided written informed consent in accordance with guidelines established by the Partners Health Care Institutional Review Board and the Harvard University Committee on the Use of Human Subjects in Research (See Supplementary Appendix A in (Holmes et al., 2015)).

#### Structural imaging processing

All imaging data were collected on matched 3T Tim Trio scanners (Siemens Healthcare, Erlangen, Germany) at Harvard University and Massachusetts General Hospital using the vendor-supplied 12-channel phased-array head coil. Structural data included a high-resolution (1.2mm isotropic) multi-echo T1-weighted magnetization-prepared gradient-echo image (multi-echo MPRAGE, see further: (Holmes et al., 2015)). The low participant burden resulting from the use of multi-echo MPRAGE anatomical scans makes this sequence well suited for high-throughput studies. The morphometric features derived through conventional 6-min 1mm MPRAGE and the 2-min 1.2mm multi-echo MPRAGE are highly consistent (r2>0.9 for most structures) suggesting that rapid acquisition multi-echo MPRAGE can be used for many purposes in place of longer anatomical scans without degradation of the quantitative morphometric estimates. All T1 scans pre-processed using the Freesurfer software library (http://surfer.nmr.mgh.harvard.edu/) version 6.0.0 (Dale et al., 1999; Fischl, 2013; Fischl and Dale, 2000; Fischl et al., 1999). Next, the individual cortical thickness and surface area maps were standardized to fsaverage5 for further analysis. Segmentation maps were visually inspected (S.L.V.) for inaccuracies and 1 individual was excluded due to regional abnormalities.

#### Five Factor Model of Personality

The Big Five personality traits were assessed using the full-length 240-item Revised NEO Personality Inventory NEO-Five-Factors-Inventory (NEO-PI-R)(Costa and McCrae, 1992), the full-length 240-item NEO-PI-R. For each item, participants reported their level of agreement on a 5-point Likert scale, from strongly disagree to strongly agree. The NEO instruments have been previously validated in USA and several other countries (McCrae and Terracciano, 2005). As a proxy for IQ we used the estimated IQ derived through the Oklahoma Premorbid Intelligence Estimate--3 (OPIE3) formula (Schoenberg et al., 2002). Reported values are in integers and binned. It is of note that distribution of IQ values is positively skewed relative to the general population and that many personality traits, including negative affect and Neuroticism were observed to have distribution that would be expected of a clinically-screened population-based sample (Holmes et al., 2015). See further **Table 2**.

**Table 2.** Behavioral characteristics of the GSP sample. Behavioral characteristics for gender, age, intelligence as well as the NEO-FFI scores in the GSP sample.

| Measure | n | mean±SD (min-max) |
| --- | --- | --- |
| *Males/Females* | 536/390 | - |
| *Age* | 926 | 21.6±3.9 (19-35) |
| *Estimated IQ* | 891 | 108.7±8.1 (77-129) |
| *Agreeableness* | 926 | 32.0±6.6 (9-47) |
| *Conscientiousness* | 926 | 31.7±7.2 (8-48) |
| *Extraversion* | 926 | 30.7±6.5 (9-48) |
| *Neuroticism* | 926 | 20.3±8.8 (0-48) |
| *Openness* | 926 | 31.6±6.1 (14-47) |

### eNKI sample

#### Participants and study design

To evaluate the cross-sample reproducibility of observations we additionally investigated correspondence between personality and cortical brain structure in the eNKI where we selected adults between 20 and 40 years of age to match the age-range of the HCP and GSP samples. The sample was made available by the Nathan-Kline Institute (NKY, NY, USA), as part of the ‘*enhanced NKI-Rockland sample*’ (https://www.ncbi.nlm.nih.gov/pmc/articles/PMC3472598/). In short, eNKI was designed to yield a community-ascertained, lifespan sample in which age, ethnicity, and socioeconomic status are representative of Rockland County, New York, U.S.A. ZIP-code based recruitment and enrollments efforts were being used to avoid over-representation of any portion of the community. Participants below 6 years were excluded to balance data losses with scientific yield, as well as participants above the age of 85, as chronic illness was observed to dramatically increase after this age. All approvals regarding human subjects’ studies were sought following NKI procedures. Scans were acquired from the International Neuroimaging Data Sharing Initiative (INDI) online database http://fcon_1000.projects.nitrc.org/indi/enhanced/studies.html For our phenotypic analyses, we selected individuals with complete personality and imaging data within the age-range of 18-40 years to match the age-range of the HCP and GSP samples. Our sample for phenotypic correlations consisted of 210 (121 females) individuals with mean age of 26.0 years (SD =6.1, min-max =18-39). Please see **Table 3** for demographic characteristics.

**Table 3.** Behavioral characteristics of the eNKI sample. Behavioral characteristics for gender, age, intelligence as well as the NEO-FFI scores in the eNKI sample.

| Measure | n | mean±SD (min-max) |
| --- | --- | --- |
| *Males/Females* | 121/210 | - |
| *Age* | 210 | 26.0±6.1 (18-39) |
| *Intelligence (WASI)* | 210 | 100.3±12.3 (69-135) |
| *Agreeableness* | 210 | 33.6±6.1 (18-48) |
| *Conscientiousness* | 210 | 33.9±7.2 (13-48) |
| *Extraversion* | 210 | 30.5±6.3 (7-44) |
| *Neuroticism* | 210 | 19.7±8.1(2-42) |
| *Openness* | 210 | 33.0±6.3 (12-48) |

#### Structural imaging processing

3D magnetization-prepared rapid gradient-echo imaging (3D MP-RAGE) structural scans(Mugler and Brookeman, 1990) were acquired using a 3.0□T Siemens Trio scanner with TR=2500□ms, TE=3.5□ms, Bandwidth=190□Hz/Px, field of view=256 × 256□mm, flip angle=8°, voxel size=1.0 × 1.0 × 1.0□mm. More details on image acquisition are available at http://fcon_1000.projects.nitrc.org/indi/enhanced/studies.html. All T1 scans were pre-processed using the Freesurfer software library (http://surfer.nmr.mgh.harvard.edu/) version 6.0.0 (Dale et al., 1999; Fischl, 2013; Fischl and Dale, 2000; Fischl et al., 1999) to compute cortical thickness and surface area. Next, the individual cortical thickness and surface area maps were standardized to fsaverage5 for further analysis. Segmentations were visually inspected for anatomical errors (S.L.V.).

#### Five Factor Model of Personality

The Big Five personality traits were assessed using the NEO-FFI3(McCrae and Costa, 2004; McCrae and Terracciano, 2005).

For an assessment of intelligence we used the Wechsler Abbreviated Scale of Intelligence (WASI-II)(Wechsler, 1999), full scale IQ. The WASI is a general intelligence, or IQ test designed to assess specific and overall cognitive capabilities and is individually administered to children, adolescents and adults (ages 6-89). It is a battery of four subtests: Vocabulary (31-item), Block Design (13-item), Similarities (24-item) and Matrix Reasoning (30-item). In addition to assessing general, or Full Scale, intelligence, the WASI is also designed to provide estimates of Verbal and Performance intelligence consistent with other Wechsler tests. Specifically, the four subtests comprise the full scale and yield the Full Scale IQ (FSIQ-4), see further **Table 3**.

#### Parcellation approach

In all three samples, we used a parcellation scheme (Schaefer et al., 2018) based on the combination of a local gradient approach and a global similarity approach using gradient-weighted Markov Random models. The parcellation has been extensively evaluated with regards to stability and convergence with histological mapping and alternative parcellations. In the context of the current study, we focus on the granularity of 200 parcels, as averaging will improve signal-to-noise ratio. In order to improve signal-to-noise ratio and to accelerate analysis speed, we opted to average unsmoothed structural data within each parcel. Thus, cortical thickness of each ROI was estimated as the trimmed mean (10 percent trim) and surface area as the sum of area within an ROI.

#### Phenotypic correlation analysis

As in previous structural MRI analyses (Bernhardt et al., 2014; Valk et al., 2016a; Valk et al., 2016b), we used SurfStat for Matlab [R2017a, The Mathworks, Natick, MA](Worsley et al., 2009). Phenotypic correlation analyses between personality traits and local brain structure were carried out per parcel, using a 200 parcel-parcellation scheme (Schaefer et al., 2018) on surface area and cortical thickness. We controlled for the same variables as in the genetic analysis, namely age, sex, age × sex interaction, age^2^, age^2^ × sex interaction, as well as global thickness effects when investigating cortical thickness and intracranial volume when assessing surface area. Results were corrected for multiple comparisons using Benjamini-Hochberg FDR (Benjamini and Hochberg, 1995) at whole-brain analysis, where we corrected for number of analysis within the current step and report FDR thresholds. When investigating personality or in post-hoc brain analysis, we corrected for number of analysis x ROIs. Post-hoc we also controlled for a proxy for intelligence, total cognitive score (Weintraub et al., 2013). We displayed significant (FDRq<0.05) findings on the brain surface.

#### Heritability and genetic correlation analysis

To investigate the heritability and genetic correlation of brain structure and personality traits, we analyzed 200 parcels of cortical thickness and surface area, as well as personality trait score of each subject in a twin-based heritability analysis. As in previous studies (Glahn et al., 2010), the quantitative genetic analyses were conducted using Sequential Oligogenic Linkage Analysis Routines (SOLAR) (Almasy and Blangero, 1998). SOLAR uses maximum likelihood variance-decomposition methods to determine the relative importance of familial and environmental influences on a phenotype by modeling the covariance among family members as a function of genetic proximity. This approach can handle pedigrees of arbitrary size and complexity and thus, is optimally efficient with regard to extracting maximal genetic information. To ensure that our traits, behavioral as well as of brain structure, were conform to the assumptions of normality, an inverse normal transformation was applied for all behavioral and neuroimaging traits (Glahn et al., 2010).

Heritability (*h*^*2*^) represents the portion of the phenotypic variance (σ^2^_p_) accounted for by the total additive genetic variance (σ^2^_g_), i.e., *h*^*2*^ = σ^2^_g_ /σ^2^_p_. Phenotypes exhibiting stronger covariances between genetically more similar individuals than between genetically less similar individuals have higher heritability. Heritability analyses were conducted with simultaneous estimation for the effects of potential covariates. For this study, we included covariates including age, sex, age × sex interaction, age^2^, age^2^ × sex interaction. Post-hoc we also controlled for a proxy for intelligence, total cognitive score(Weintraub et al., 2013). When investigating cortical thickness, we additionally controlled for global thickness effects (mean cortical thickness) and in case of surface area we controlled for intracranial volume.

To determine if variations in personality and brain structure were influenced by the same genetic factors, genetic correlation analyses were conducted. More formally, bivariate polygenic analyses were performed to estimate genetic (ρ_g_) and environmental (ρ_e_) correlations, based on the phenotypic correlation (ρ_p_), between brain structure and personality with the following formula: ρ_p_ = ρ_g_√(*h*^*2*^_1_*h*^*2*^_2_) + ρ_e_√[(1 − *h*^*2*^_1_)(1 − *h*^*2*^_2_)], where *h*^*2*^_1_and *h*^*2*^_2_ are the heritability’s of the parcel-based cortical thickness and the various behavioral traits. The significance of these correlations was tested by comparing the log likelihood for two restricted models (with either ρ_g_ or ρ_e_ constrained to be equal to 0) against the log likelihood for the model in which these parameters were estimated. A significant genetic correlation (corrected for multiple comparisons using Benjamin-Hochberg FDR (Benjamini and Hochberg, 1995)) is evidence suggesting that (a proportion of) both phenotypes are influenced by a gene or set of genes (Almasy et al., 1997).

To compute the contribution of genetic effects relative to the phenotypic correlation, we computed the contribution of the genetic path to the phenotypic correlation (√ *h*^*2*^_1_× ρ_g_ × √ *h*^*2*^_2_) (ρ_ph_g) divided by the phenotypic correlation. For the relative contribution of environmental correlation to the phenotypic correlation we computed (√ 1-*h*^*2*^_1_× ρ_e_ × √ *1-h*^*2*^_2_) (ρ_ph_e) divided by the phenotypic correlation (Zheng et al., 2019).

#### Bayes Factors of replication

To compare the evidence that the personality-local brain structure could be replicated in two independent samples (H1, replication, and H0, no replication), we additionally quantified personality-brain associations within each ROI, using Bayes factors (Verhagen and Wagenmakers, 2014). In line with previous work of our group (Kharabian Masouleh et al., 2020; Kharabian Masouleh et al., 2019) Bayes factors (BF) were summarized into four categories. These categories are used to simplify the interpretation and comparison of replication rates. For example, a BF_01_ lower than 1/3 shows that the data is three times or more likely to have happened under H1 than H0. “Successful” replication is defined as a replication lower than 1 in both replication samples.

#### Functional decoding

Parcel that were significantly replicated in at least one sample were functionally characterized using the Behavioral Domain meta-data from the BrainMap database (http://www.brainmap.org (Laird et al., 2011; Laird et al., 2009)). To do so, volumetric counterparts, delineating the surface-based parcels in volume space, as provided by Schaefer, (Schaefer et al., 2018) (https://github.com/ThomasYeoLab/CBIG/tree/master/stable_projects/brain_parcellation/Schaefer2018_LocalGlobal/Parcellations) were used. In particular, we identified those meta-data labels (describing the computed contrast [behavioral domain as well as paradigm]) that were significantly more likely than chance to result in activation of a given parcel (Fox et al., 2014; Genon et al., 2018; Nostro et al., 2017). That is, functions were attributed to the parcels by quantitatively determining which types of experiments are associated with activation in the respective parcellation region. Significance was established using a binomial test (q < 0.05, corrected for multiple comparisons using false discovery rate, FDR).

## Results

### Association between personality traits and cortical brain structure

To assess the association between personality and macroscale cortical brain structure we first evaluated distribution of behavioral measures. Using the Kolmogorov-Smirnov test we found that all personality traits in the HCP sample (n=1106 including 286 MZ-twins and 170 DZ-twins) were conform to normal distributions (KS-score between 0.97 and 1) (**Figure 1**). We observed significant phenotypic correlation between all personality traits, with the exception of Openness and Neuroticism (r =0.01) (**Figure 1**, Supplementary Table 1).

**Figure 1.**
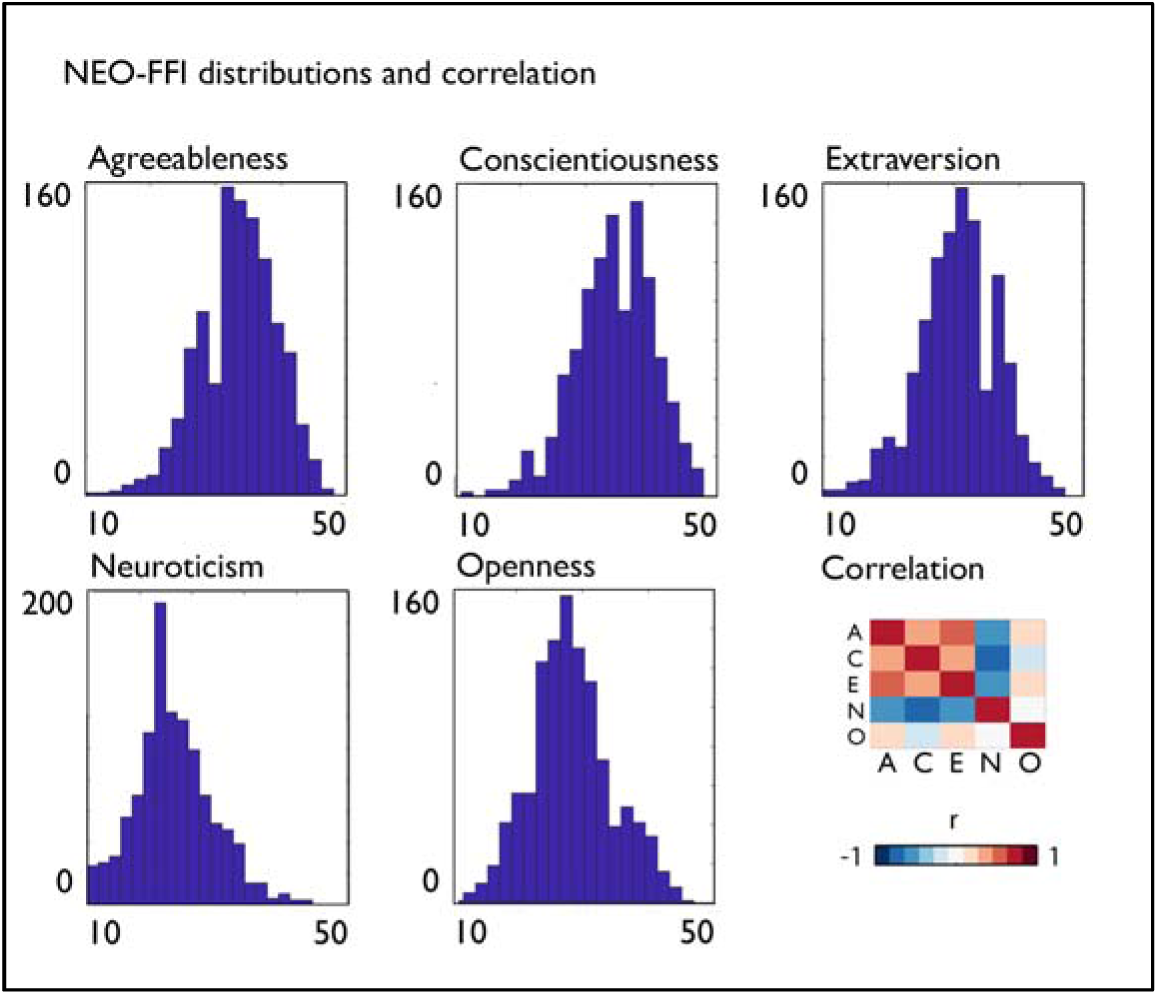
Distribution of personality traits in the full HCP sample. Distribution of NEO-FFI personality traits in the HCP dataset, score on x-axis, number of occurrences on the y-axis, as well as the correlation between NEO-FFI traits in the HCP sample.

Next, we assessed phenotypic correlation between personality traits and cortical structure, specifically cortical thickness and surface area. Distribution of cortical thickness values summarized in 200 functionally informed parcels (Schaefer et al., 2018) showed highest thickness in anterior insula, and relatively low values in occipital regions (**Figure 2A**). At the regional level, we observed correlations between Agreeableness, Neuroticism, and Openness and local cortical thickness (**Figure 2B**). Specifically, Agreeableness related negatively to variations in thickness in left lateral and bilateral medial prefrontal cortex (FDRq<0.01). Neuroticism related positively to thickness in dorsolateral frontal areas and left posterior operculum, and negatively to thickness in left posterior occipital regions (FDRq<0.015). Openness related negatively to thickness in left ventrolateral cortex, and positively to right temporal pole (FDRq<0.01). We did not observe significant associations between mean cortical thickness and personality scores (**Table 4**).

**Table 4.** Association between personality traits and whole brain summaries of surface area and cortical thickness in the full HCP sample. T-values of the association between average cortical thickness and total surface area and personality traits. ** indicates FDRq<0.05, * indicates p<0.05.

|  | Average cortical thickness | Total surface area |
| --- | --- | --- |
| Agreeableness | -0.24 | 0.66 |
| Conscientiousness | 0.55 | -2.59** |
| Extraversion | 0.66 | 1.14 |
| Neuroticism | 1.90* | -2.03* |
| Openness | -1.28 | 2.94** |

**Figure 2.**
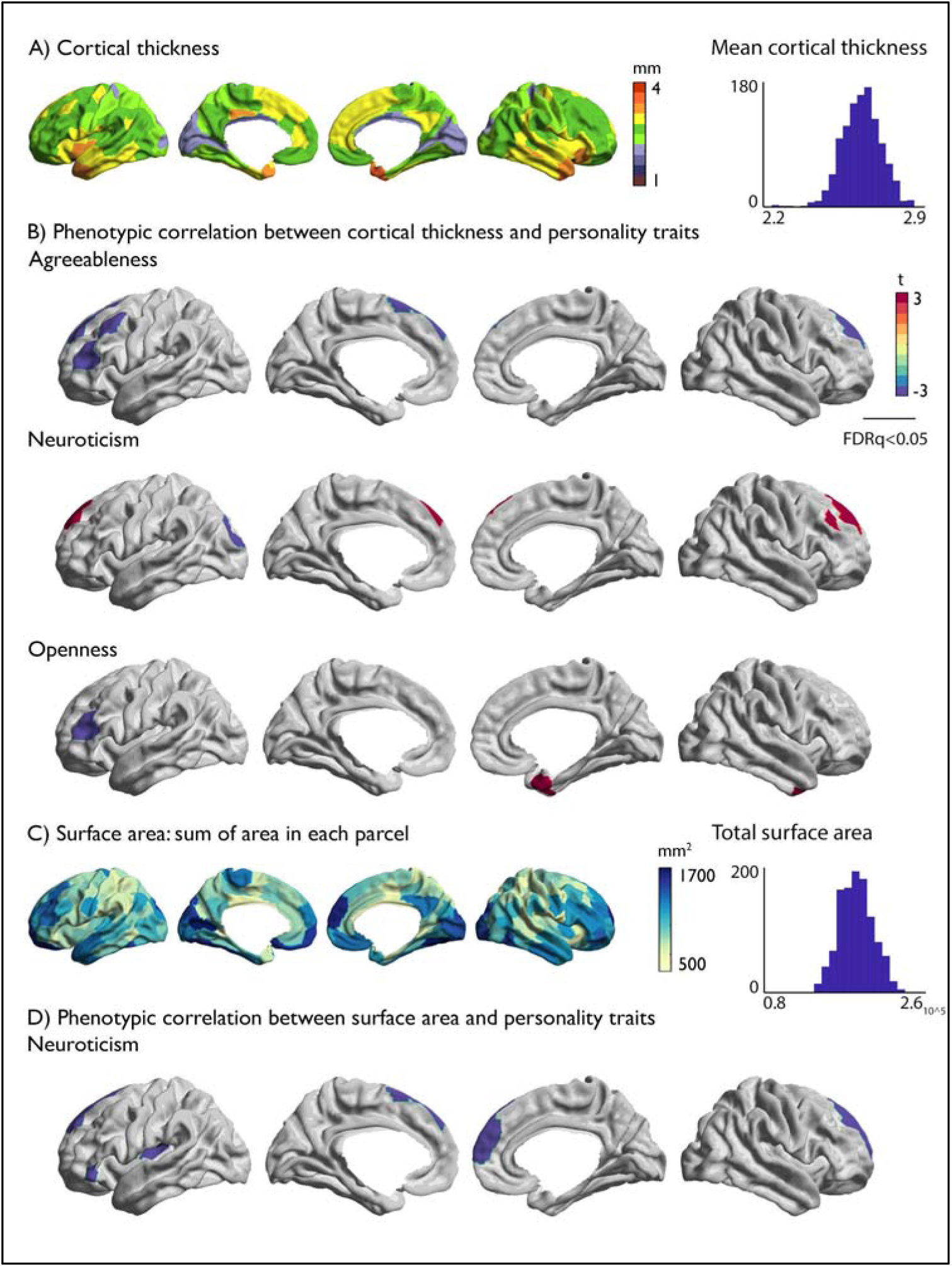
Relation between personality traits and local brain structure in the full HCP sample. A) Mean cortical thickness of each parcel and the distribution of average cortical thickness across participants; B) Regional associations between personality traits and cortical thickness; C) Average surface area sum per parcel and the distribution of total surface area across participants; D) Regional associations between surface area and personality traits. Positive associations between local brain structure and each personality trait are displayed in red and negative associations displayed in blue. Multiple comparisons were accounted for by using FDR corrections at q<0.05 correcting for the number of parcels (200) and only significant associations are displayed.

Total surface area had a negative relation with conscientiousness (t=-2.59, p<0.005) and a positive association with openness (t=2.94, p<0.002) (**Table 4**). Regionally, we found a negative relation between Neuroticism and local surface area in bilateral medial frontal cortex, left inferior frontal gyrus, and left posterior insula (FDRq<0.02).

To test stability of our findings we additionally evaluated the robustness of phenotypic associations between personality and global and local brain structure while controlling for total cognitive score and the other personality traits (Supplementary Materials, Supplementary Table 2 and 3). While all local associations remained significant at p<0.01, strength of associations was generally reduced and few regions reached FDRq<0.05 significance levels.

### Genetic relationship between personality traits and cortical brain structure

Subsequently, we sought to evaluate whether the phenotypic correlations observed in the twin-sample were due to shared genetic or environmental effects on grey matter brain structure and personality traits. All personality traits were significantly heritable in our current sample (**Figure 3C**, Supplementary Table 4), as were mean cortical thickness (h^2^=0.85) and total surface area (h^2^=0.93), and we confirmed also local cortical thickness (h^2^: mean±std: 0.33±0.11) and surface area (h^2^: mean±std: 0.41±0.13) to be heritable in our parcel-based approach (**Figure 3A-B**, Supplementary Table 5 and 11).

**Figure 3.**
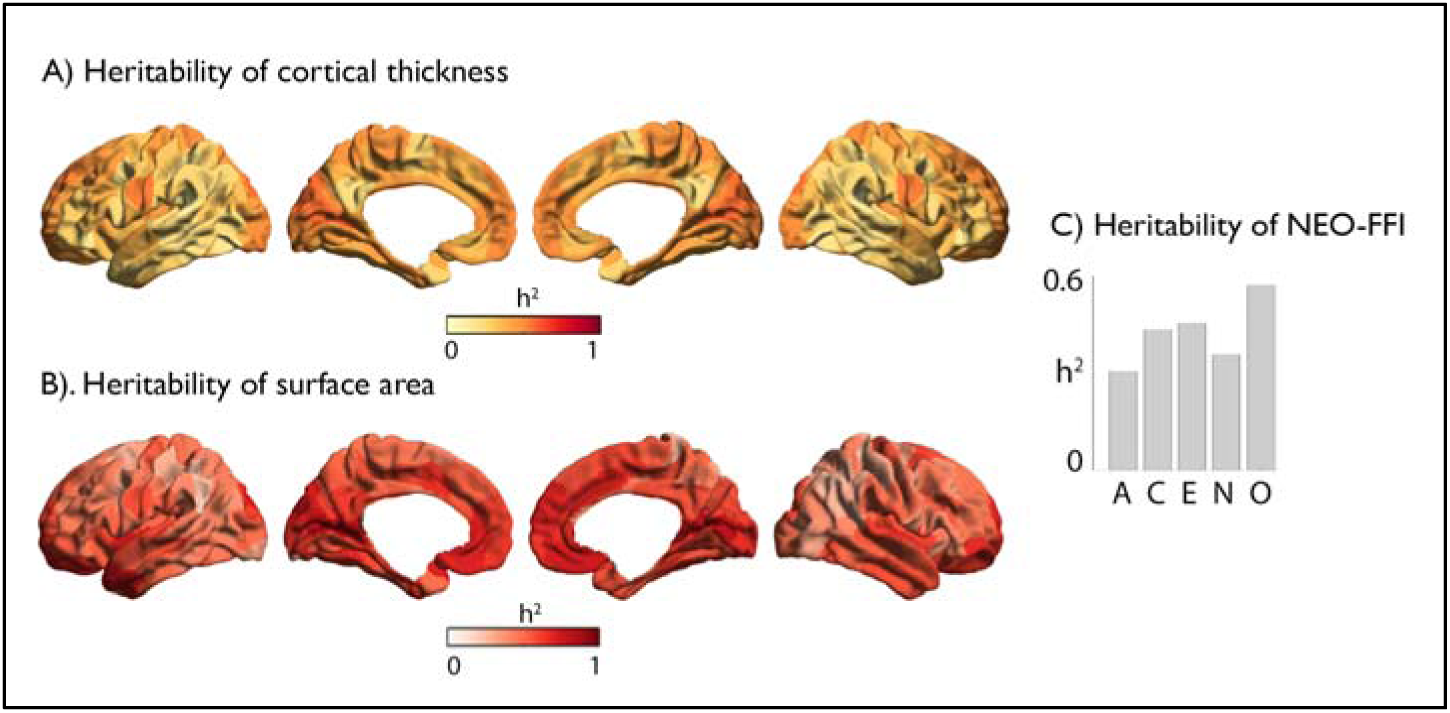
Heritability of local cortical structure and personality traits. A) Heritability of local cortical thickness; B) Heritability of surface area; C) Heritability of NEO-FFI: A= agreeableness, C=conscientiousness, E=extraversion, N=Neuroticism, O=Openness

Following, we assessed genetic correlation between personality traits and cortical structure. We did not observe genetic or environmental associations between personality and global thickness (**Table 5**), however, the phenotypic association between total surface area and conscientiousness was observed to be driven by shared genetic effects (ρ_g_ =-0.12, p<0.03) whereas the association between openness and total surface area was driven by environmental effects (ρ_e_=0.18, p<0.025).

**Table 5.** Genetic and environmental correlation between personality traits and whole brain summaries of surface area and cortical thickness. Genetic and environmental correlations are computed in the HCP sample, and exact p-values are reported, associations that did not show phenotypic correlation at p<0.05 threshold are in bold.

|  | Global thickness |  | Total surface area |  |
| --- | --- | --- | --- | --- |
| **Agreeableness** | $\rho_e = 0.03$, $p = \text{ns}$; | $\rho_g = -0.03$, $p = \text{ns}$ | $\rho_e = -0.08$, $p = \text{ns}$; | $\rho_g = 0.07$, $p = \text{ns}$ |
| **Conscientiousness** | $\rho_e = -0.02$, $p = \text{ns}$; | $\rho_g = 0.05$, $p = \text{ns}$ | **$\rho_e = 0.04$, $p = \text{ns}$;** | **$\rho_g = -0.12$, $p = 0.03$** |
| **Extraversion** | $\rho_e = -0.03$, $p = \text{ns}$; | $\rho_g = 0.06$, $p = \text{ns}$ | $\rho_e = 0.03$, $p = \text{ns}$; | $\rho_g = 0.04$, $p = \text{ns}$ |
| **Neuroticism** | **$\rho_e = 0.10$, $p = \text{ns}$;** | **$\rho_g = 0.02$, $p = \text{ns}$** | **$\rho_e = -0.09$, $p = \text{ns}$;** | **$\rho_g = -0.06$, $p = \text{ns}$** |
| **Openness** | $\rho_e = -0.03$, $p = \text{ns}$; | $\rho_g = -0.03$, $p = \text{ns}$ | **$\rho_e = 0.18$, $p = 0.025$;** | **$\rho_g = 0.08$, $p = \text{ns}$** |

Last, we evaluated the genetic correlation of regions that showed phenotypic correlations between personality and local brain structure. We observed that 10 out of 17 phenotypic correlations showed a genetic correlation (p≤0.05), and 2 out of 17 phenotypic correlates related to an environmental correlation (p≤0.05) (**Table 6**). More specifically, we found a negative genetic correlation between Agreeableness and bilateral superior frontal thickness (p<0.04), a positive genetic correlation between Neuroticism and right superior and lateral frontal cortex thickness (p<0.05) and a positive genetic correlation between right temporal pole thickness and Openness (p<0.005). Neuroticism had a negative genetic correlation between local surface area in left posterior insula, and bilateral superior frontal cortex, and right medial frontal regions (p<0.05). See Supplementary Tables (6-10 and 12-16) for genetic and environmental correlations between personality traits and all parcels.

**Table 6.** Genetic and environmental correlation of personality brain associations in the full HCP sample. Genetic and environmental correlations are computed in the HCP sample, and exact p-values are reported. ** denotes a significant genetic correlation at FDRq<0.05, corrected for the number of ROIs associated with the respective personality trait within the structural marker. * indicated an association of p<0.05. The genetic contribution of phenotypic correlation was computed using the respective heritability of the personality trait and the local parcel as well as their genetic and phenotypic correlation.

| Cortical thickness | ROI | Environmental correlation | Genetic correlation | Genetic contribution to phenotypic correlation |
| --- | --- | --- | --- | --- |
| **Agreeableness** | LH Cont_PFCI_4 | $\rho_e$ -0.12, $p=0.04^*$ | $\rho_g$ -0.08, $p=0.61$ | 20 % |
| | LH Default_PFC_9 | $\rho_e$ -0.05, $p=0.45$ | $\rho_g$ -0.21, $p=0.08$ | 72% |
| | LH Default_PFC_11 | $\rho_e$ 0.01, $p=0.90$ | $\rho_g$ -0.36, $p=0.004^{**}$ | 100% |
| | LH Default_PFC_13 | $\rho_e$ -0.03, $p=0.72$ | $\rho_g$ -0.26, $p=0.04^*$ | 87% |
| | LH Default_PFCm_5 | $\rho_e$ 0.03, $p=0.64$ | $\rho_g$ -0.33, $p=0.005^{**}$ | 100% |
| **Neuroticism** | LH Vis_14 | $\rho_e$ -0.08, $p=0.22$ | $\rho_g$ -0.18, $p=0.12$ | 61% |
| | LH Default_PFC_9 | $\rho_e$ 0.08, $p=0.23$ | $\rho_g$ 0.17, $p=0.13$ | 61% |
| | RH Cont_PFCI_6 | $\rho_e$ 0.03, $p=0.63$ | $\rho_g$ 0.26, $p=0.05^*$ | 82% |
| | RH Default_PFCm_5 | $\rho_e$ 0.04, $p=0.54$ | $\rho_g$ 0.21, $p=0.05^*$ | 79% |
| **Openness** | LH Cont_PFCI_4 | $\rho_e$ -0.13, $p=0.05^*$ | $\rho_g$ -0.15, $p=0.17$ | 46% |
| | RH Limbic_TempPole_1 | $\rho_e$ -0.01, $p=0.94$ | $\rho_g$ 0.37, $p<0.005^{**}$ | 100% |
| **Surface area** |  |  |  |  |
| **Neuroticism** | LH SomMot_3 | $\rho_e$ 0.02, $p=0.78$ | $\rho_g$ -0.28, $p=0.01^*$ | 100% |
| | LH Default_PFC_3 | $\rho_e$ -0.04, $p=0.52$ | $\rho_g$ -0.18, $p=0.12$ | 74% |
| | LH Default_PFC_9 | $\rho_e$ 0.13, $p=0.07$ | $\rho_g$ -0.45, $p=0.0002^{**}$ | 100% |
| | LH Default_PFC_13 | $\rho_e$ -0.03, $p=0.60$ | $\rho_g$ -0.21, $p=0.14$ | 76% |
| | RH Default_PFCm_4 | $\rho_e$ 0.02, $p=0.80$ | $\rho_g$ -0.25, $p=0.02^*$ | 100% |
| | RH Default_PFCm_5 | $\rho_e$ -0.04, $p=0.55$ | $\rho_g$ -0.30, $p=0.02^*$ | 82% |

### Cross-sample reproducibility of the association between personality trait and local brain structure

In the previous analysis steps, we could show that a) there is a significant relationship between local cortical structure and personality traits in a large-scale twin sample (HCP) and that b) this relationship can be, in part, attributed to shared genetic factors. Following, to study whether associations between local brain structure and personality traits are generalizable, we evaluate the phenotypic correlation between personality traits and cortical phenotypes observed in the HPC sample are reproducible in two age-matched samples of young adults (GSP and eNKI). To formalize the level of reproducibility, we computed Bayer Factors (BF) summarizing the evidence of a successful reproduction across samples (Verhagen and Wagenmakers, 2014).

We found moderate to anecdotal evidence of replication for only one personality-brain association in both samples; the relationship between local surface area in right medial frontal cortex and Neuroticism (GSP: t=-1.54, p<0.07; BF=0.68; and eNKI: t=-1.99, p<0.025; BF=0.16). Various associations between local cortical thickness and personality traits could be reproduced in one of both replication samples (**Table 7**). Specifically, in GSP, the association between thickness in right superior frontal cortex and Agreeableness (t=-1.71, p<0.05; BF=0.52), and between thickness of right dorsal lateral PFC and Neuroticism (t=2.08, p<0.02; BF=0.18). In the eNKI sample we observed some evidence of successful replication of the association between left visual cortex and Neuroticism (t=-1.73, p<0.05; BF=0.24), left dorsolateral prefrontal thickness and Openness (t=-0.96, p>0.1, BF=0.82), surface area of left sensory-motor cortex (t=-1.09, p>0.1, BF=0.58) and left prefrontal cortex (t=-1.09, p>0.1, BF=0.58) and Neuroticism. Global measures of cortical thickness and surface area did not replicate out of sample, only in case of the positive association between total surface area and Openness we observed anecdotal evidence of successful replication in the eNKI sample (t=1.55, p<0.1, BF=0.33) (Supplementary Table 17).

**Table 7.**
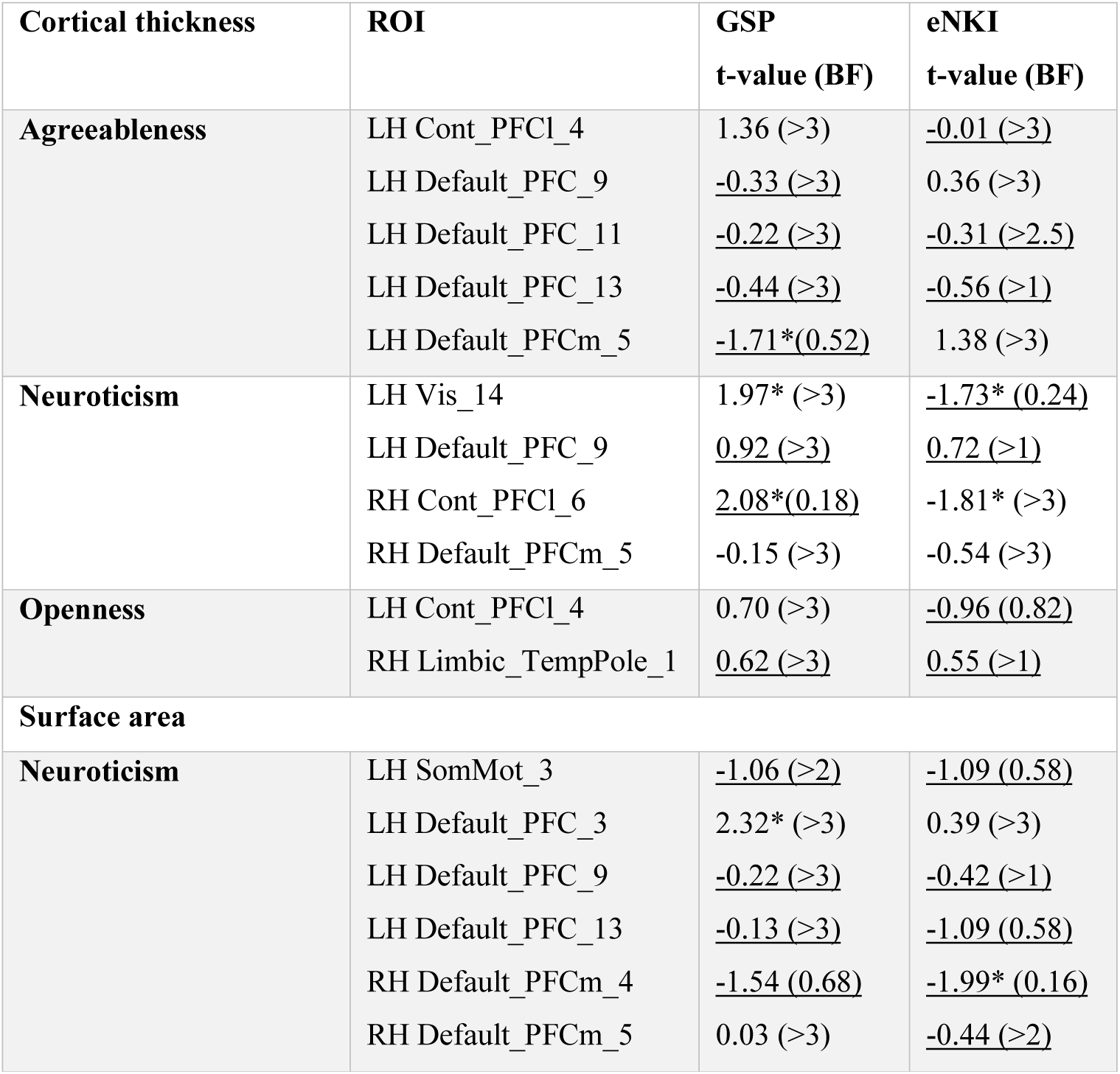
Replication of personality brain associations. Replication in the GSP and eNKI sample of significant associations between personality and local brain structure observed in the HCP sample, t-values as well as Bayes Factors (BF) are reported. If a BF_01_ is between 0 and 1/3 there is a moderate/strong evidence for H1 (replication), between 1/3 and 1 anecdotal evidence for H1, between 1 and 3 anecdotal evidence for H0 (no replication) and >3 moderate to strong evidence of H0. We underlined replications with a correct sign, and made replications bold if there is evidence (anecdotal/moderate/strong) of H1. ** indicates a significant correlation at FDRq<0.05, * is p<0.05.

### Quantitative functional decoding

Last, we performed quantitative functional mapping of the personality – brain relationships for which we observed a) phenotypic and genetic correlation in the HCP sample b) an association (p<0.05) in combination with a BF of <1 in at least one additional sample.

The right medial frontal cortex, where we observed a robust association between surface area and Neuroticism, was functionally involved in various emotional domains, social cognition, and memory, and active in paradigms involving self-reflection, Theory of Mind, and emotion induction (FDRq<0.05) (**Figure 4**).

**Figure 4.**
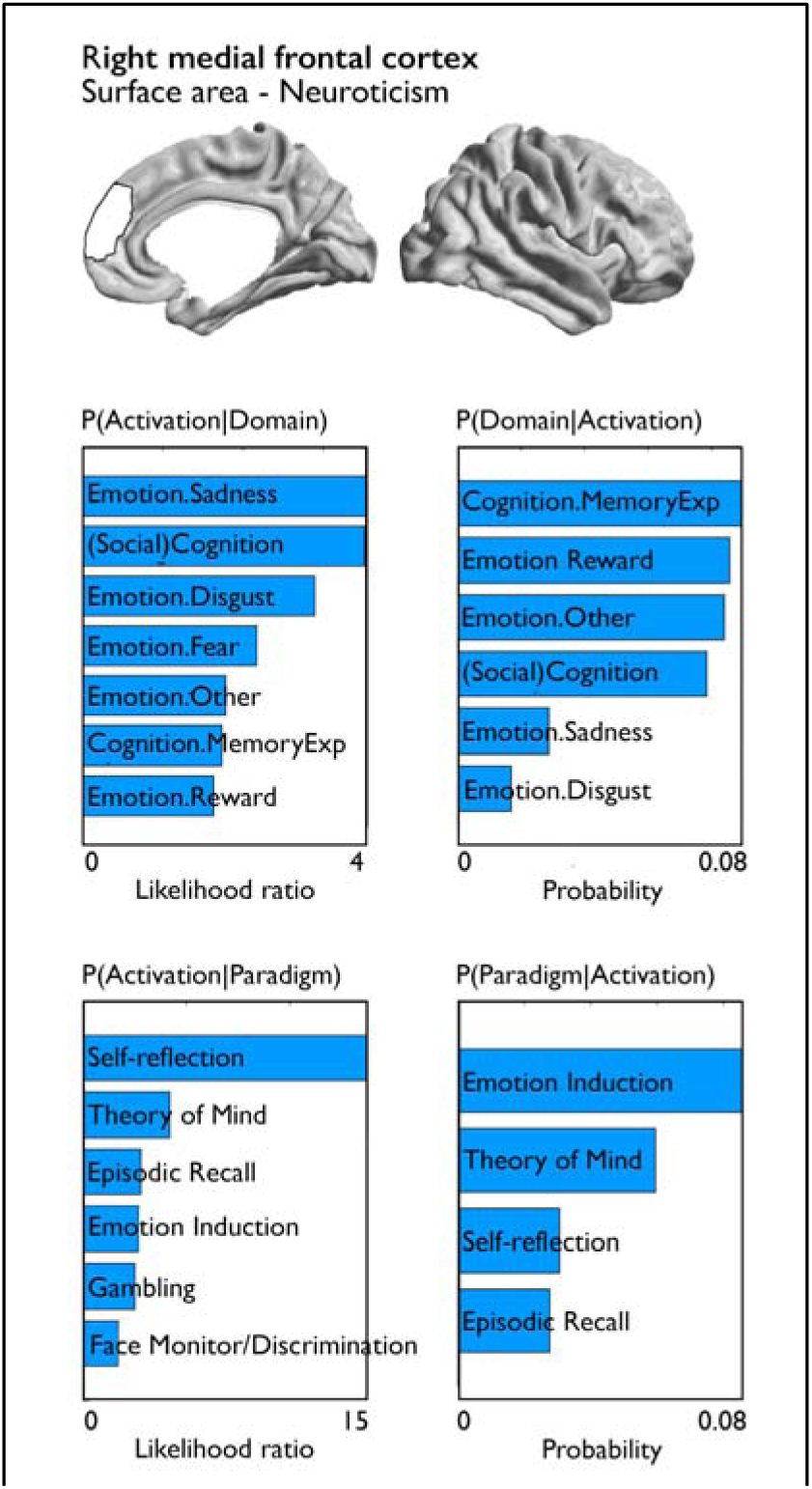
Quantitative functional decoding of consistent associations between personality and local brain structure. Both forward inference and reverse inference of activation-domain and paradigm-domain contrasts are reported for the right medial frontal cortex which showed evidence of successful replication in two samples.

## Discussion

Both local brain structure and personality are heritable. Moreover, a large body of evidence has suggested a relationship between personality and local brain structure. However, effects are weak and vary as a function of sample and effect size. In the current study, we used the large scale and openly available HCP dataset which included monozygotic and dizygotic twins to study whether there is a genetic correlation between local brain structure and personality traits. Second, we evaluated the robustness of personality-brain relationships in two additional age-matched samples.

First, we identified phenotypic associations between personality traits and local cortical structure. Associations between personality on the one hand and cortical thickness and surface area on the other were predominantly observed in frontal cortices. Performing genetic correlation analysis, we found that 10 of 17 phenotypic associations could be explained by shared genetic effects. To evaluate whether observed relationship between personality traits and local brain structure were generalizable to other samples, we additionally studied phenotypic correlations between personality traits and brain structure in two independent age-matched samples of unrelated individuals (GSP and eNKI). Here, we found that surface area in right medial prefrontal cortex was robustly associated with Neuroticism across all three samples. In sum, our findings suggest that part of phenotypic associations between personality and local brain structure can be attributed to shared genetic effects in a large-scale twin sample. However, associations were weak and the association between surface area in right medial prefrontal cortex and Neuroticism replicated in two independent samples.

We assessed the genetic basis of the association between personality and cortical thickness using compressed surface-based MRI data based on the parcellation scheme of Schaefer et al. (2018). Using compressed features of structural MRI has been suggested to both improve signal-to-noise ratio of brain measures (cf. (Eickhoff et al., 2018) and (Genon et al., 2018)), and optimize analysis scalability. The Schaefer parcellation is derived using functional MRI data from ∼1500 subjects and integrates local approaches detecting abrupt transitions in functional connectivity patterns and global approaches that cluster similar functional connectivity patterns (Schaefer et al., 2018). Indeed, a combination of within-area micro circuitry, proxied by brain morphometry, and between-area connectivity enables each area to perform a unique set of computations (Van Essen and Glasser, 2018). Therefore, a parcellation approach that considers both local and global connectivity might benefit structural image analysis, as it reduces signal-to-noise both within and across individuals and makes control for multiple comparisons more straightforward (Genon et al., 2018). Based on the findings in our study, we suggest our approach might be a fruitful first exploratory step to investigate the genetic relation between brain structure and behavior, and locate mechanisms of interest. Future studies can subsequently verify these results by exploring more specific genetic mechanisms, as well as neuroanatomical features.

Though we could establish phenotypic correlations between personality traits and local cortical thickness, associations were weak and phenotypic associations ranged between t-values of 3.5 and −3.5. In the HCP dataset phenotypic correlations between predominantly frontal regions and personality traits of Agreeableness, Neuroticism, and Openness have been previously reported using a non-parcel-based method by Owens and colleagues (Owens et al., 2019). Frontal cortices are functionally involved in a number of tasks involving higher cognitive functioning, such as executive functioning, memory, metacognition and social cognition (Amodio and Frith, 2006; Baird et al., 2013; Bludau et al., 2014; Buckner et al., 2008; Fleming and Dolan, 2012; Valk et al., 2016a). We additionally observed various relationships between global measures of surface area and cortical thickness on the one hand and personality traits on the other. Indeed, we could replicate a recently reported association between total surface area and Neuroticism in phenotypic correlation analysis (Grasby et al., 2020). However, associations between global measures of cortical structure and personality were not consistent across samples.

We extend previously reported phenotypic observations by showing that these phenotypic relationships between personality and local cortical structure are driven, in part, by shared additive genetic effects rather than environmental factors alone. The contribution of genetic effects on phenotypic correlations is dependent on the heritability of each of the correlated markers. In our sample, between 30 % and 60 % (on average 42 %) of variance in personality traits was explained by additive genetic factors. This is in line with previous studies using twin and family samples (Jang et al., 1996) as well as genome-wide approaches (Lo et al., 2017). A recent meta-analysis (Vukasovic and Bratko, 2015) confirmed that on average 40 % of the variance in personality traits is of genetic origin. Also, conform with previous studies (Eyler et al., 2012; Kremen et al., 2010; Panizzon et al., 2009; Strike et al., 2018; Winkler et al., 2010), we observe heritability of local cortical thickness, with highest values in primary sensory areas. Heritability patterns followed previously described patterns with relatively strong genetic influence on cortical thickness in unimodal cortices, whereas variance in association cortices is on average less influenced by genetic factors (Eyler et al., 2012; Grasby et al., 2020; Hofer and al., 2018; Kremen et al., 2010; Panizzon et al., 2009; Strike et al., 2018; Winkler et al., 2010). Also, local surface area was heritable, with lowest heritability values in dorsolateral PFC and temporal-parietal regions (Eyler et al., 2012; Grasby et al., 2020; Hofer and al., 2018; Kremen et al., 2010; Panizzon et al., 2009; Strike et al., 2018; Winkler et al., 2010).

Performing genetic correlation analysis, we observed that the phenotypic correlation between personality and local brain structure in 10 out of 17 regions was driven by genetic factors. These regions were predominantly located in frontal areas, suggesting a genetic link between local structure in frontal cortices and personality. Indeed, various studies have suggested a relationship between personality and the frontal lobe in humans (DeYoung et al., 2010; Owens et al., 2019; Riccelli et al., 2017) and in chimpanzees (Latzman et al., 2015). There are various ways in which a genetic process would affect the relationship between personality and cortical macrostructure and it is likely the observed genetic correlations between local cortical structure and personality traits in the HCP sample are be due to mediated pleiotropy (a gene affects A which affects B). On the one hand, it could be a genetic factor affects grey matter macrostructure and associated function and, as a consequence, personality. On the other hand, it could be that genetic variation affects brain function which in turn modulates both macroscale structure as well as personality, or a genetic mechanism affects an unknown biological factor which in turn affects personality and brain structure. Recent work using GWAS and genetic correlation in a large sample of individuals could found a genetic association between cortical brain structure and various markers of behavior, providing first evidence of a direct link of genes and behavior via cortical brain structure (Grasby et al., 2020). Here, Grasby et al found evidence that genetic regulatory elements influencing local surface area and local cortical thickness stem from different devepemental mechanisms. Whereas surface area is associated with genetic variants active during fetal development, cortical thickness may reflect genetic processes underlying myelination, branching and pruning. Such differential mechanistic timing effects on cortical structure might contribute to the understanding of which biological mechanisms underlie personality, and further dissociate factors that shape personality across the life-span.

As various studies have indicated relationships between local brain structure and psychometric variables are not robust (Avinun et al., bioarXiv; Kharabian Masouleh et al., 2020; Kharabian Masouleh et al., 2019), we further evaluated the robustness of phenotypic associations between personality and local brain structure in two age-matched samples of unrelated individuals. Indeed, though most associations did not replicate across all three samples, the association between medial prefrontal surface area and Neuroticism was observed in all three samples. Functional decoding indicated that this region is functionally involved in (social)-cognitive and emotional processing. Additionally, we found anecdotal to moderate evidence for successful replication of various associations cortical thickness and personality in either GSP or eNKI sample. However, given the inconsistency across samples, these replications are challenging to interpret.

### Limitations and outlook

Moving forward, there are various limitations and challenges in operationalizing personality that might have resulted in a lack of consistent findings across samples. Though our samples all were from WEIRD (Western, educated, industrialized, rich, and democratic) populations (Laajaj et al., 2019), it might be that personality traits probed are not comparable across samples due to challenges to reliably operationalize personality, and that confounding environmental and noise effects vary across samples. For example, it is possible inconsistent or lack of findings with regard to macroscale neuroanatomical associations of personality may be a function of the assessment of personality used (in this case, the NEO-FFI/NEO-PI-R) rather than a true null or unreliable finding (Avinun et al., bioarXiv). The five-factor personality model and the subsequent operationalizations in instruments such as the NEO are based on a lexical approach. Though such an approach might be able to dissociate various personality traits, it is debated whether lexical taxonomy has a direct relation to neurobiology (Perkins et al., 2020; Yarkoni, 2015). Future studies might benefit from using personality instruments developed in concordance with brain structure and function such as Hierarchical Taxonomy of Psychopathology (HiTOP) (Perkins et al., 2020).

Second, a recent review on the neurobiology of personality suggested that rather than focusing on a one-to-one mapping between personality and neurobiology, as done in the current study, studies that seek to identify mechanisms contributing to particular clusters of behaviors might be a more fruitful approach to capture the neurobiological mechanisms underlying personality traits (Yarkoni, 2015). For example, though brain structure is a viable endophenotype of personality, correlation between personality and macro-scale cortical structure is weak. Thus, further study of the relationship between personality and functional activity and functional dynamics might further contribute to understanding the biological basis of personality and other complex traits (Dubois et al., 2018; Kebets et al., 2019; Kong et al., 2019; Wu et al., 2019).

Third, only 40% of personality variance in the current sample could attributed to genetic effects. Environment, such as family environment, peer-groups, and stress have been reported to influence personality (Hopwood et al., 2011; Nakao et al., 2000), and also local cortical structure and associated behavior has been reported to change as a consequence of changing environments in adulthood (Valk et al., 2017). Though genetic and gene by environment effects are not to be excluded in this context, is likely such environmental mechanisms further shape the relation between personality traits and brain structure, above and beyond direct additive genetic effects. Longitudinal designs might help to further understand the environmental relationship between personality and brain structure and function.

Taken together, in the current study we report evidence of a shared genetic basis of personality traits and local brain structure within the HCP sample, and a robust association of local surface area in medial prefrontal regions and Neuroticism across three independent samples. It is of note that our study on the shared genetic basis of personality and brain structure was made possible by the open HCP, GSP, and eNKI neuroimaging repositories. These initiatives offer cognitive neuroimaging communities an unparalleled access to large datasets for the investigation of the brain basis of individual difference. They have also enabled us to highlight variability across samples and validation experiments to verify stability of our observations. Notably, the use of multiple datasets enabled us to test robustness of our findings. Given that reproducibility is increasingly important, our study illustrates the advantages of open data to increase understanding of complex traits.

## Acknowledgements

We would like to thank the various contributors to the open access databases that our data was downloaded from. Specifically; HCP data were provided by the Human Connectome Project, Washington University, the University of Minnesota, and Oxford University Consortium (Principal Investigators: David Van Essen and Kamil Ugurbil;1U54MH091657) funded by the 16 NIH Institutes and Centers that support the NIH Blueprint for Neuroscience Research; and by the McDonnell Center for Systems Neuroscience at Washington University. GSP data were provided by the Brain Genomics Superstruct Project of Harvard University and the Massachusetts General Hospital, (Principal Investigators: Randy Buckner, Joshua Roffman, and Jordan Smoller), with support from the Center for Brain Science Neuroinformatics Research Group, the Athinoula A. Martinos Center for Biomedical Imaging, and the Center for Human Genetic Research. 20 individual investigators at Harvard and MGH generously contributed data to the overall project.

For enhanced NKI, we would like to thank the principal support for the enhanced NKI-RS project is provided by the NIMH BRAINS R01MH094639-01 (PI Milham). Funding for key personnel was also provided in part by the New York State Office of Mental Health and Research Foundation for Mental Hygiene. Funding for the decompression and augmentation of administrative and phenotypic protocols provided by a grant from the Child Mind Institute (1FDN2012-1). Additional personnel support provided by the Center for the Developing Brain at the Child Mind Institute, as well as NIMH R01MH081218, R01MH083246, and R21MH084126. Project support also provided by the NKI Center for Advanced Brain Imaging (CABI), the Brain Research Foundation (Chicago, IL), and the Stavros Niarchos Foundation. Last, we want to thank L.C.Valk for proofreading the manuscript.

