## Supplementary Material for "Personality and local brain structure: their shared genetic basis and reproducibility"

**Supplement**

Sofie L. Valk^1,2*^; Felix Hoffstaedter^1,2^; Julia A. Camilleri^1,2^; Peter Kochunov^3^; B.T. Thomas Yeo^4,5,6^; Simon B. Eickhoff^1,2^

*^1)^ Institute of Neuroscience and Medicine (INM-7: Brain and Behaviour), Research Centre Jülich, 52425 Jülich, Germany*

*^2)^ Institute of Systems Neuroscience, Heinrich Heine University Düsseldorf, 40225 Düsseldorf, Germany*

*^3)^ Maryland Psychiatric Research Center, University of Maryland School of Medicine, Baltimore, Maryland, USA.*

*^4)^ Department of Electrical and Computer Engineering, Centre for Sleep & Cognition, Clinical Imaging Research Centre, and N.1 Institute for Health, National University of Singapore, Singapore, Singapore.*

*^5)^ Athinoula A. Martinos Center for Biomedical Imaging, Massachusetts General Hospital, Charlestown, MA, USA.*

*^6)^ NUS Graduate School for Integrative Sciences and Engineering, National University of Singapore, Singapore, Singapore.*

**Supplementary results**

*Phenotypic and genetic correlation between personality traits.*

We also evaluated whether the phenotypic correlations observed between personality traits in HCP could reflect shared genetic or environmental processes (Supplementary Table 18 and 19). We observed significant genetic correlation between Extraversion and Agreeableness (ρ_g_ = 0.38) and Extraversion and Conscientiousness (ρ_g_ = 0.33). Significant environmental correlations were observed between Agreeableness and Conscientiousness (ρ_e_ = 0.32), Agreeableness and Extraversion (ρ_e_ = 0.26), Agreeableness and Neuroticism (ρ_e_ = -0.41), Conscientiousness and Extraversion (ρ_e_ = 0.24), Conscientiousness and Neuroticism (ρ_e_ = -0.51), and Extraversion and Neuroticism (ρ_e_ = -0.39).

*Whole brain genetic correlation between personality and brain structure*

Additionally, we performed exploratory whole brain analysis of genetic correlations between personality traits and cortical thickness to evaluate whether we could identify regions that showed significant genetic correlation with personality traits. Agreeableness showed genetic correlation (FDRq<0.05, corrected for number of parcels) with cortical thickness in left posterior superior/mid temporal sulcus (*LH_Default_Temp_5*, t=1.11, p=ns; ρ_g_ = 0.60, p<0.0005; ρ_e_ = -0.15, p<0.01), right superior frontal cortex extending to mid cingulate (*RH_SalVentAttn_Med_3*, t=-0.90, p=ns; ρ_g_ = -0.55, p<0.0004; ρ_e_ =0.18, p<0.005), and right dorsolateral frontal cortex (*RH_Default_PFCm_6*, t=-0.01, p=ns; ρ_g_ = -0.51, p<0.0006; ρ_e_ =0.21, p<0.0004). Second, we observed a genetic correlation between Extraversion and left TPJ thickness (*LH_Default_Temp_9*, t=3.10, p<0.001; ρ_g_ = 0.49 p<0.00005; ρ_e_ = -0.11, p<0.05).

Last, we observed significant genetic correlation between Neuroticism and surface area of left superior frontal cortex (*'7Networks_LH_Default_PFC_9'*, t=-3.46, p<0.0001; ρ_g_ = -0.45, p<0.0002; ρ_e_ = 0.13, p=ns). See Supplementary Tables for genetic and environmental correlations between personality traits and all parcels.

**Supplementary Figures**

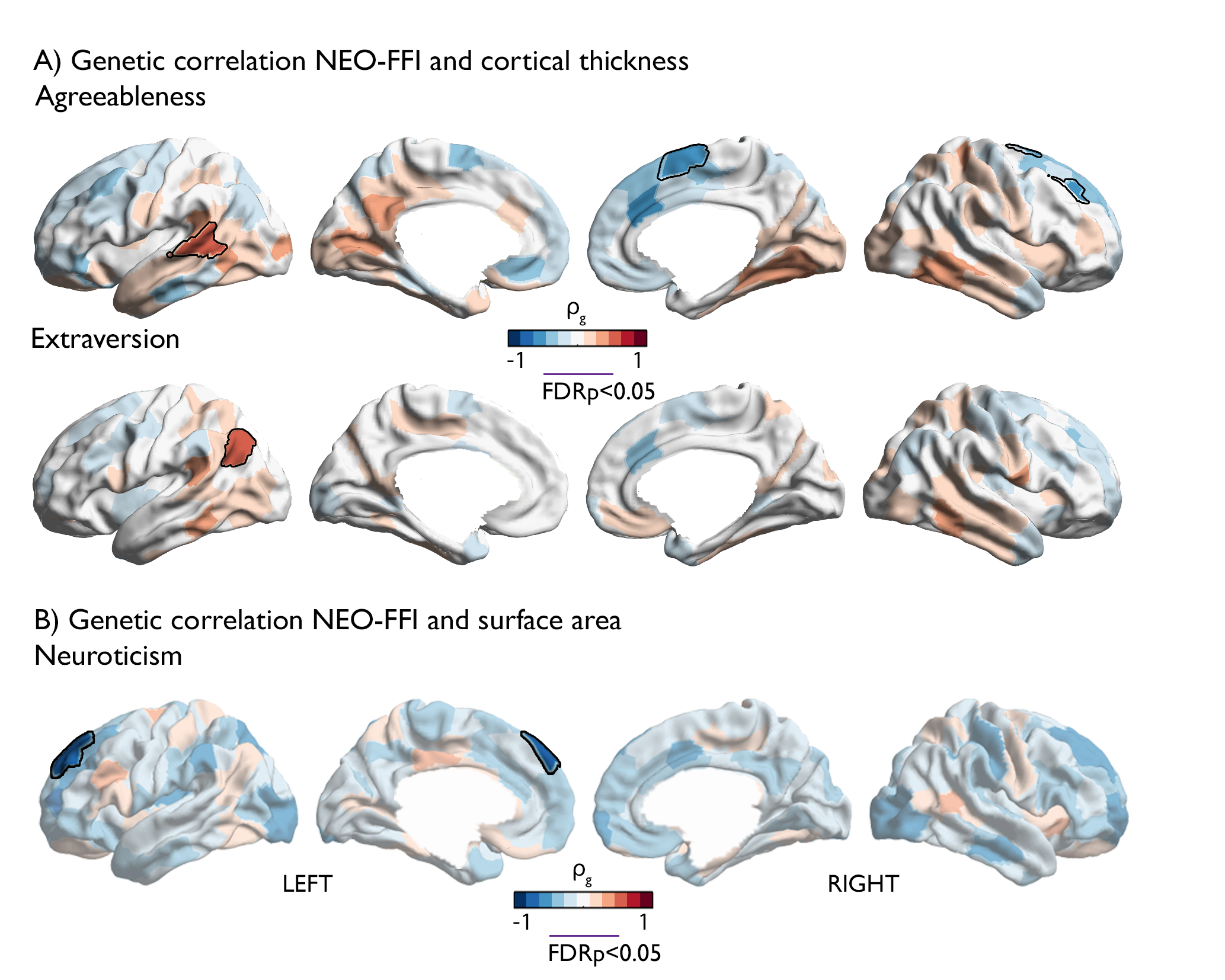

**Supplementary Figure 1. Genetic correlation between personality traits local brain structure.** A) Genetic correlation between cortical thickness and personality traits; B) Genetic correlation between local surface area and personality traits. Red values indicate positive associations whereas blue values indicate negative genetic correlation between parcel-wise thickness and personality traits, FDRq findings have black outline. Only personality-brain associations containing FDRq<0.05 results are displayed. .

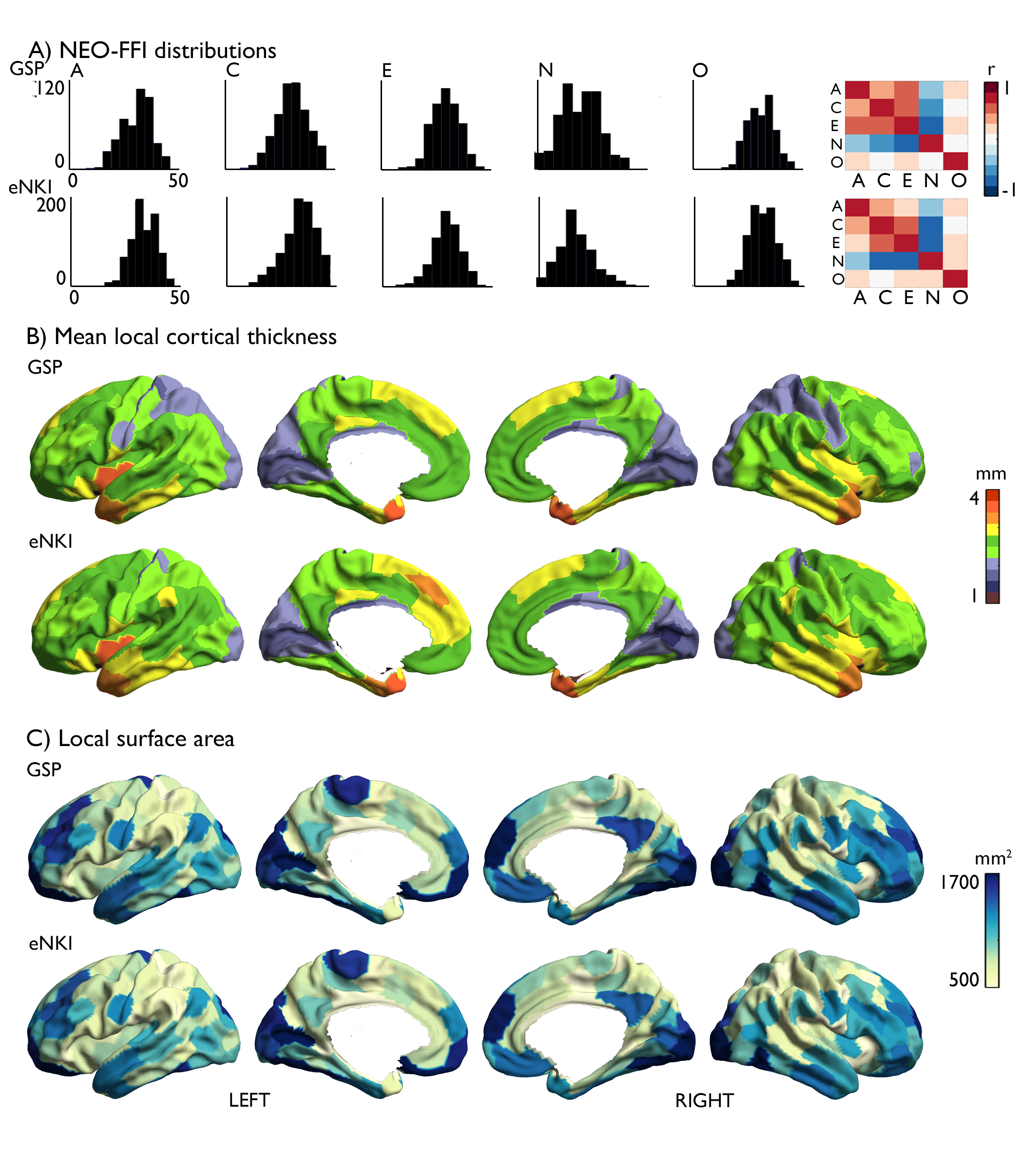

**Supplementary Figure 2. Distribution of NEO-FFI scores, their intercorrelation, and parcel-wise cortical thickness and surface area in two unrelated samples.** A). Distributions of NEO-FFI scores in two unrelated samples (GSP; eNKI); right: Intercorrelation of personality trait scores in each sample. B). Distribution of mean parcel-wise thickness in mm in the two unrelated samples; C).

**Supplementary Tables**

| Personality trait | A | C | E | N | O |
| --- | --- | --- | --- | --- | --- |
| *Agreeableness* | 1 |  |  |  |  |
| *Conscientiousness* | 0.23 ** |  |  |  |  |
| *Extraversion* | 0.28 ** | 0.27 ** |  |  |  |
| *Neuroticism* | -0.28 ** | -0.40 ** | -0.34 ** |  |  |
| *Openness* | 0.09 ** | -0.13 ** | 0.10 ** | 0.01 |  |

**Supplementary Table 1. Phenotypic correlation of personality traits in HCP.** Phenotypic correlation across personality traits. A=Agreeableness, C=Conscientiousness, E=Extraversion, N=Neuroticism, O=Openness. ** indicates corrected at FDRq<0.05, controlled for number of analysis (10), * indicates p<0.05.

|  | Global thickness | | Total surface area | |
| --- | --- | --- | --- | --- |
|  | IQ | Other personality score | IQ | Other personality score |
| Agreeableness | -0.21, p>0.1 | 0.20, p>0.1 | -0.21, p>0.1 | -0.17, p>0.1 |
| Conscientiousness | 0.42, p>0.1 | 1.05, p>0.1 | -1.97, p<0.025 | -3.49, p<0.0005 |
| Extraversion | 0.79, p>0.1 | 1.31, p<0.1 | 1.47, p<0.1 | 0.74, p>0.1 |
| Neuroticism | 1.73, p<0.05 | 2.55, p<0.01 | -1.20, p>0.1 | -2.98, p<0.002 |
| Openness | -1.26, p>0.1 | -1.32, p<0.1 | 0.94, p>0.1 | 2.41, p<0.01 |

**Supplementary Table 2. Phenotypic correlations between personality traits and global brain structure in HCP controlling for IQ and personality scores.** Post-hoc analysis of the association between personality and global brain structure (t-values) when controlling for IQ and NEO-FFI personality scores.

| Cortical thickness | ROI | IQ  t-value, p, (FDR) | Other personality scores |
| --- | --- | --- | --- |
| Agreeableness | LH Cont_PFCl_4  LH Default_PFC_9  LH Default_PFC_11  LH Default_PFC_13  LH Default_PFCm_5 | -3.26, p<0.0005 (0)  -3.29, p<0.0005 (0)  -3.71, p<0.0001 (1)  -3.09, p<0.001 (0)  -2.73, p<0.0025 (0) | -2.65, p<0.005 (0)  -2.41, p<0.005 (0)  -3.76, p<0.0001 (1)  -2.88, p<0.005 (0)  -2.41, p<0.01 (0) |
| Neuroticism | LH Vis_14  LH Default_PFC_9  RH Cont_PFCl_6  RH Default_PFCm_5 | -3.75, p<0.0001 (1)  3.15, p<0.001 (0)  2.84, p<0.0025 (0)  2.48, p<0.01 (0) | -3.23, p<0.001 (0)  2.86, p<0.0025 (0)  2.94, p<0.0025 (0)  2.94, p<0.0025 (0) |
| Openness | LH Cont_PFCl_4  RH Limbic_TempPole_1 | -3.53, p<0.0002 (0)  2.91, p<0.0025 (0) | -3.69, p<0.0001 (1)  3.69, p<0.0001 (1) |
| Surface area | | | |
| Neuroticism | LH SomMot_3  LH Default_PFC_3  LH Default_PFC_9  LH Default_PFC_13  RH Default_PFCm_4  RH Default_PFCm_5 | -3.23, p<0.0005 (0)  -3.11, p<0.001 (0)  -3.41, p<0.0005 (0)  -3.12, p<0.001 (0)  -2.79, p<0.005 (0)  -3.87, p<0.0001 (1) | -3.47, p<0.0005 (1)  -2.18, p<0.025 (0)  -3.73, p<0.0001 (1)  -3.00, p<0.0025 (1)  -3.09, p<0.0001 (1)  -4.28, p<0.0001 (1) |

**Supplementary Table 3. Phenotypic correlations in parcels with FDR-corrected association between personality traits and local brain structure in HCP controlling for IQ and personality scores.** Robustness of phenotypic association between parcels that showed significant (FDRq<0.05) phenotypic correlation in the HCP sample and the corresponding personality trait when controlling for IQ (Total Cognitive score) and other NEO-FFI traits. T-values are reported, as well as p-values. If the result would survive FDR corrections (FDRq<0.05) at whole brain level (1) is reported.

| Personality trait | H^2^ | SE | p |
| --- | --- | --- | --- |
| *Agreeableness* | 0.29 | 0.07 | 0.0000 |
| *Conscientiousness* | 0.43 | 0.06 | 0.0000 |
| *Extraversion* | 0.43 | 0.05 | 0.0000 |
| *Neuroticism* | 0.37 | 0.06 | 0.0000 |
| *Openness* | 0.58 | 0.05 | 0.0000 |

**Supplementary Table 4. Heritability of personality traits.** Heritability of personality traits computed using Solar 8.4.0. Here we report heritability values, standard error, as well as p-values (displayed at 10^-4^ accuracy).

| **ROIS** | **H^2^** | **SE** | **p** |
| --- | --- | --- | --- |
| **7Networks_LH_Vis_1** | 0.3960 | 0.0506 | 0.0000 |
| **7Networks_LH_Vis_2** | 0.3280 | 0.0555 | 0.0000 |
| **7Networks_LH_Vis_3** | 0.1523 | 0.0586 | 0.0036 |
| **7Networks_LH_Vis_4** | 0.4836 | 0.0439 | 0.0000 |
| **7Networks_LH_Vis_5** | 0.4083 | 0.0522 | 0.0000 |
| **7Networks_LH_Vis_6** | 0.4480 | 0.0501 | 0.0000 |
| **7Networks_LH_Vis_7** | 0.4963 | 0.0470 | 0.0000 |
| **7Networks_LH_Vis_8** | 0.2402 | 0.0513 | 0.0000 |
| **7Networks_LH_Vis_9** | 0.4775 | 0.0471 | 0.0000 |
| **7Networks_LH_Vis_10** | 0.5194 | 0.0463 | 0.0000 |
| **7Networks_LH_Vis_11** | 0.3290 | 0.0522 | 0.0000 |
| **7Networks_LH_Vis_12** | 0.4718 | 0.0516 | 0.0000 |
| **7Networks_LH_Vis_13** | 0.5551 | 0.0466 | 0.0000 |
| **7Networks_LH_Vis_14** | 0.4452 | 0.0519 | 0.0000 |
| **7Networks_LH_SomMot_1** | 0.4736 | 0.0497 | 0.0000 |
| **7Networks_LH_SomMot_2** | 0.2755 | 0.0555 | 0.0000 |
| **7Networks_LH_SomMot_3** | 0.5145 | 0.0480 | 0.0000 |
| **7Networks_LH_SomMot_4** | 0.3997 | 0.0531 | 0.0000 |
| **7Networks_LH_SomMot_5** | 0.2452 | 0.0547 | 0.0000 |
| **7Networks_LH_SomMot_6** | 0.5276 | 0.0462 | 0.0000 |
| **7Networks_LH_SomMot_7** | 0.2888 | 0.0575 | 0.0000 |
| **7Networks_LH_SomMot_8** | 0.2808 | 0.0589 | 0.0000 |
| **7Networks_LH_SomMot_9** | 0.3392 | 0.0544 | 0.0000 |
| **7Networks_LH_SomMot_10** | 0.5155 | 0.0479 | 0.0000 |
| **7Networks_LH_SomMot_11** | 0.2151 | 0.0550 | 0.0000 |
| **7Networks_LH_SomMot_12** | 0.3585 | 0.0543 | 0.0000 |
| **7Networks_LH_SomMot_13** | 0.3296 | 0.0584 | 0.0000 |
| **7Networks_LH_SomMot_14** | 0.3638 | 0.0514 | 0.0000 |
| **7Networks_LH_SomMot_15** | 0.4356 | 0.0492 | 0.0000 |
| **7Networks_LH_SomMot_16** | 0.4002 | 0.0578 | 0.0000 |
| **7Networks_LH_DorsAttn_Post_1** | 0.3267 | 0.0586 | 0.0000 |
| **7Networks_LH_DorsAttn_Post_2** | 0.2151 | 0.0535 | 0.0000 |
| **7Networks_LH_DorsAttn_Post_3** | 0.3442 | 0.0544 | 0.0000 |
| **7Networks_LH_DorsAttn_Post_4** | 0.3499 | 0.0561 | 0.0000 |
| **7Networks_LH_DorsAttn_Post_5** | 0.2148 | 0.0550 | 0.0000 |
| **7Networks_LH_DorsAttn_Post_6** | 0.2283 | 0.0590 | 0.0000 |
| **7Networks_LH_DorsAttn_Post_7** | 0.4602 | 0.0527 | 0.0000 |
| **7Networks_LH_DorsAttn_Post_8** | 0.3059 | 0.0551 | 0.0000 |
| **7Networks_LH_DorsAttn_Post_9** | 0.4502 | 0.0506 | 0.0000 |
| **7Networks_LH_DorsAttn_Post_10** | 0.4682 | 0.0529 | 0.0000 |
| **7Networks_LH_DorsAttn_FEF_1** | 0.2673 | 0.0550 | 0.0000 |
| **7Networks_LH_DorsAttn_FEF_2** | 0.3613 | 0.0562 | 0.0000 |
| **7Networks_LH_DorsAttn_PrCv_1** | 0.1866 | 0.0585 | 0.0005 |
| **7Networks_LH_SalVentAttn_ParOper_1** | 0.1303 | 0.0574 | 0.0096 |
| **7Networks_LH_SalVentAttn_ParOper_2** | 0.2800 | 0.0527 | 0.0000 |
| **7Networks_LH_SalVentAttn_ParOper_3** | 0.2185 | 0.0581 | 0.0000 |
| **7Networks_LH_SalVentAttn_FrOper_1** | 0.3500 | 0.0537 | 0.0000 |
| **7Networks_LH_SalVentAttn_FrOper_2** | 0.4083 | 0.0535 | 0.0000 |
| **7Networks_LH_SalVentAttn_FrOper_3** | 0.4316 | 0.0528 | 0.0000 |
| **7Networks_LH_SalVentAttn_FrOper_4** | 0.2426 | 0.0610 | 0.0000 |
| **7Networks_LH_SalVentAttn_PFCl_1** | 0.4391 | 0.0476 | 0.0000 |
| **7Networks_LH_SalVentAttn_Med_1** | 0.3362 | 0.0589 | 0.0000 |
| **7Networks_LH_SalVentAttn_Med_2** | 0.3337 | 0.0577 | 0.0000 |
| **7Networks_LH_SalVentAttn_Med_3** | 0.2462 | 0.0571 | 0.0000 |
| **7Networks_LH_Limbic_OFC_1** | 0.2963 | 0.0532 | 0.0000 |
| **7Networks_LH_Limbic_OFC_2** | 0.3585 | 0.0556 | 0.0000 |
| **7Networks_LH_Limbic_TempPole_1** | 0.3385 | 0.0583 | 0.0000 |
| **7Networks_LH_Limbic_TempPole_2** | 0.3692 | 0.0498 | 0.0000 |
| **7Networks_LH_Limbic_TempPole_3** | 0.2432 | 0.0529 | 0.0000 |
| **7Networks_LH_Limbic_TempPole_4** | 0.4492 | 0.0520 | 0.0000 |
| **7Networks_LH_Cont_Par_1** | 0.2170 | 0.0570 | 0.0000 |
| **7Networks_LH_Cont_Par_2** | 0.2715 | 0.0536 | 0.0000 |
| **7Networks_LH_Cont_Par_3** | 0.0448 | 0.0556 | 0.2068 |
| **7Networks_LH_Cont_Temp_1** | 0.1930 | 0.0574 | 0.0003 |
| **7Networks_LH_Cont_PFCl_1** | 0.2604 | 0.0512 | 0.0000 |
| **7Networks_LH_Cont_PFCl_2** | 0.2964 | 0.0557 | 0.0000 |
| **7Networks_LH_Cont_PFCl_3** | 0.3926 | 0.0532 | 0.0000 |
| **7Networks_LH_Cont_PFCl_4** | 0.2659 | 0.0547 | 0.0000 |
| **7Networks_LH_Cont_PFCl_5** | 0.4091 | 0.0501 | 0.0000 |
| **7Networks_LH_Cont_PFCl_6** | 0.2063 | 0.0572 | 0.0001 |
| **7Networks_LH_Cont_pCun_1** | 0.3744 | 0.0516 | 0.0000 |
| **7Networks_LH_Cont_Cing_1** | 0.5313 | 0.0487 | 0.0000 |
| **7Networks_LH_Cont_Cing_2** | 0.3678 | 0.0568 | 0.0000 |
| **7Networks_LH_Default_Temp_1** | 0.2854 | 0.0595 | 0.0000 |
| **7Networks_LH_Default_Temp_2** | 0.2831 | 0.0547 | 0.0000 |
| **7Networks_LH_Default_Temp_3** | 0.3428 | 0.0543 | 0.0000 |
| **7Networks_LH_Default_Temp_4** | 0.1930 | 0.0585 | 0.0004 |
| **7Networks_LH_Default_Temp_5** | 0.2131 | 0.0585 | 0.0001 |
| **7Networks_LH_Default_Temp_6** | 0.2821 | 0.0539 | 0.0000 |
| **7Networks_LH_Default_Temp_7** | 0.3019 | 0.0572 | 0.0000 |
| **7Networks_LH_Default_Temp_8** | 0.2422 | 0.0561 | 0.0000 |
| **7Networks_LH_Default_Temp_9** | 0.2643 | 0.0520 | 0.0000 |
| **7Networks_LH_Default_PFC_1** | 0.3635 | 0.0525 | 0.0000 |
| **7Networks_LH_Default_PFC_2** | 0.2708 | 0.0581 | 0.0000 |
| **7Networks_LH_Default_PFC_3** | 0.1493 | 0.0560 | 0.0032 |
| **7Networks_LH_Default_PFC_4** | 0.5135 | 0.0459 | 0.0000 |
| **7Networks_LH_Default_PFC_5** | 0.3168 | 0.0578 | 0.0000 |
| **7Networks_LH_Default_PFC_6** | 0.3310 | 0.0538 | 0.0000 |
| **7Networks_LH_Default_PFC_7** | 0.5053 | 0.0483 | 0.0000 |
| **7Networks_LH_Default_PFC_8** | 0.3889 | 0.0572 | 0.0000 |
| **7Networks_LH_Default_PFC_9** | 0.4656 | 0.0478 | 0.0000 |
| **7Networks_LH_Default_PFC_10** | 0.3767 | 0.0515 | 0.0000 |
| **7Networks_LH_Default_PFC_11** | 0.4263 | 0.0481 | 0.0000 |
| **7Networks_LH_Default_PFC_12** | 0.4694 | 0.0486 | 0.0000 |
| **7Networks_LH_Default_PFC_13** | 0.4649 | 0.0493 | 0.0000 |
| **7Networks_LH_Default_PCC_1** | 0.3913 | 0.0550 | 0.0000 |
| **7Networks_LH_Default_PCC_2** | 0.2794 | 0.0537 | 0.0000 |
| **7Networks_LH_Default_PCC_3** | 0.3900 | 0.0539 | 0.0000 |
| **7Networks_LH_Default_PCC_4** | 0.3922 | 0.0556 | 0.0000 |
| **7Networks_LH_Default_PHC_1** | 0.4640 | 0.0536 | 0.0000 |
| **7Networks_RH_Vis_1** | 0.3649 | 0.0531 | 0.0000 |
| **7Networks_RH_Vis_2** | 0.3496 | 0.0569 | 0.0000 |
| **7Networks_RH_Vis_3** | 0.3773 | 0.0566 | 0.0000 |
| **7Networks_RH_Vis_4** | 0.5840 | 0.0417 | 0.0000 |
| **7Networks_RH_Vis_5** | 0.2794 | 0.0551 | 0.0000 |
| **7Networks_RH_Vis_6** | 0.4662 | 0.0502 | 0.0000 |
| **7Networks_RH_Vis_7** | 0.5079 | 0.0463 | 0.0000 |
| **7Networks_RH_Vis_8** | 0.4341 | 0.0539 | 0.0000 |
| **7Networks_RH_Vis_9** | 0.5773 | 0.0433 | 0.0000 |
| **7Networks_RH_Vis_10** | 0.4768 | 0.0535 | 0.0000 |
| **7Networks_RH_Vis_11** | 0.2379 | 0.0536 | 0.0000 |
| **7Networks_RH_Vis_12** | 0.5188 | 0.0479 | 0.0000 |
| **7Networks_RH_Vis_13** | 0.4779 | 0.0490 | 0.0000 |
| **7Networks_RH_Vis_14** | 0.4356 | 0.0522 | 0.0000 |
| **7Networks_RH_Vis_15** | 0.2731 | 0.0558 | 0.0000 |
| **7Networks_RH_SomMot_1** | 0.4370 | 0.0532 | 0.0000 |
| **7Networks_RH_SomMot_2** | 0.3421 | 0.0590 | 0.0000 |
| **7Networks_RH_SomMot_3** | 0.3498 | 0.0573 | 0.0000 |
| **7Networks_RH_SomMot_4** | 0.3403 | 0.0554 | 0.0000 |
| **7Networks_RH_SomMot_5** | 0.1911 | 0.0576 | 0.0003 |
| **7Networks_RH_SomMot_6** | 0.2971 | 0.0572 | 0.0000 |
| **7Networks_RH_SomMot_7** | 0.4346 | 0.0501 | 0.0000 |
| **7Networks_RH_SomMot_8** | 0.1921 | 0.0546 | 0.0001 |
| **7Networks_RH_SomMot_9** | 0.3351 | 0.0603 | 0.0000 |
| **7Networks_RH_SomMot_10** | 0.1718 | 0.0602 | 0.0016 |
| **7Networks_RH_SomMot_11** | 0.3010 | 0.0532 | 0.0000 |
| **7Networks_RH_SomMot_12** | 0.4576 | 0.0501 | 0.0000 |
| **7Networks_RH_SomMot_13** | 0.3075 | 0.0531 | 0.0000 |
| **7Networks_RH_SomMot_14** | 0.3848 | 0.0524 | 0.0000 |
| **7Networks_RH_SomMot_15** | 0.3496 | 0.0590 | 0.0000 |
| **7Networks_RH_SomMot_16** | 0.2815 | 0.0593 | 0.0000 |
| **7Networks_RH_SomMot_17** | 0.4880 | 0.0494 | 0.0000 |
| **7Networks_RH_SomMot_18** | 0.3781 | 0.0504 | 0.0000 |
| **7Networks_RH_SomMot_19** | 0.4382 | 0.0534 | 0.0000 |
| **7Networks_RH_DorsAttn_Post_1** | 0.2420 | 0.0563 | 0.0000 |
| **7Networks_RH_DorsAttn_Post_2** | 0.2431 | 0.0565 | 0.0000 |
| **7Networks_RH_DorsAttn_Post_3** | 0.3086 | 0.0533 | 0.0000 |
| **7Networks_RH_DorsAttn_Post_4** | 0.2197 | 0.0572 | 0.0000 |
| **7Networks_RH_DorsAttn_Post_5** | 0.1940 | 0.0576 | 0.0002 |
| **7Networks_RH_DorsAttn_Post_6** | 0.3929 | 0.0523 | 0.0000 |
| **7Networks_RH_DorsAttn_Post_7** | 0.1424 | 0.0564 | 0.0047 |
| **7Networks_RH_DorsAttn_Post_8** | 0.3550 | 0.0542 | 0.0000 |
| **7Networks_RH_DorsAttn_Post_9** | 0.3334 | 0.0539 | 0.0000 |
| **7Networks_RH_DorsAttn_Post_10** | 0.4139 | 0.0513 | 0.0000 |
| **7Networks_RH_DorsAttn_FEF_1** | 0.3090 | 0.0586 | 0.0000 |
| **7Networks_RH_DorsAttn_FEF_2** | 0.2329 | 0.0569 | 0.0000 |
| **7Networks_RH_DorsAttn_PrCv_1** | 0.3154 | 0.0553 | 0.0000 |
| **7Networks_RH_SalVentAttn_TempOccPar_1** | 0.2310 | 0.0572 | 0.0000 |
| **7Networks_RH_SalVentAttn_TempOccPar_2** | 0.1702 | 0.0549 | 0.0008 |
| **7Networks_RH_SalVentAttn_TempOccPar_3** | 0.2479 | 0.0547 | 0.0000 |
| **7Networks_RH_SalVentAttn_PrC_1** | 0.1571 | 0.0568 | 0.0022 |
| **7Networks_RH_SalVentAttn_FrOper_1** | 0.2728 | 0.0570 | 0.0000 |
| **7Networks_RH_SalVentAttn_FrOper_2** | 0.5003 | 0.0520 | 0.0000 |
| **7Networks_RH_SalVentAttn_FrOper_3** | 0.3183 | 0.0530 | 0.0000 |
| **7Networks_RH_SalVentAttn_FrOper_4** | 0.2536 | 0.0567 | 0.0000 |
| **7Networks_RH_SalVentAttn_Med_1** | 0.2638 | 0.0558 | 0.0000 |
| **7Networks_RH_SalVentAttn_Med_2** | 0.4131 | 0.0515 | 0.0000 |
| **7Networks_RH_SalVentAttn_Med_3** | 0.3440 | 0.0594 | 0.0000 |
| **7Networks_RH_Limbic_OFC_1** | 0.4121 | 0.0566 | 0.0000 |
| **7Networks_RH_Limbic_OFC_2** | 0.3526 | 0.0496 | 0.0000 |
| **7Networks_RH_Limbic_OFC_3** | 0.3785 | 0.0545 | 0.0000 |
| **7Networks_RH_Limbic_TempPole_1** | 0.1799 | 0.0560 | 0.0005 |
| **7Networks_RH_Limbic_TempPole_2** | 0.3495 | 0.0556 | 0.0000 |
| **7Networks_RH_Limbic_TempPole_3** | 0.5059 | 0.0498 | 0.0000 |
| **7Networks_RH_Cont_Par_1** | 0.1538 | 0.0562 | 0.0021 |
| **7Networks_RH_Cont_Par_2** | 0.1959 | 0.0554 | 0.0001 |
| **7Networks_RH_Cont_Par_3** | 0.1328 | 0.0566 | 0.0081 |
| **7Networks_RH_Cont_Temp_1** | 0.2258 | 0.0600 | 0.0000 |
| **7Networks_RH_Cont_PFCv_1** | 0.2109 | 0.0579 | 0.0001 |
| **7Networks_RH_Cont_PFCl_1** | 0.2699 | 0.0546 | 0.0000 |
| **7Networks_RH_Cont_PFCl_2** | 0.3055 | 0.0549 | 0.0000 |
| **7Networks_RH_Cont_PFCl_3** | 0.3266 | 0.0515 | 0.0000 |
| **7Networks_RH_Cont_PFCl_4** | 0.4177 | 0.0506 | 0.0000 |
| **7Networks_RH_Cont_PFCl_5** | 0.3271 | 0.0519 | 0.0000 |
| **7Networks_RH_Cont_PFCl_6** | 0.3110 | 0.0512 | 0.0000 |
| **7Networks_RH_Cont_PFCl_7** | 0.3489 | 0.0573 | 0.0000 |
| **7Networks_RH_Cont_pCun_1** | 0.2582 | 0.0545 | 0.0000 |
| **7Networks_RH_Cont_PFCmp_1** | 0.4633 | 0.0479 | 0.0000 |
| **7Networks_RH_Cont_PFCmp_2** | 0.4163 | 0.0537 | 0.0000 |
| **7Networks_RH_Cont_PFCmp_3** | 0.2755 | 0.0573 | 0.0000 |
| **7Networks_RH_Cont_PFCmp_4** | 0.4503 | 0.0546 | 0.0000 |
| **7Networks_RH_Default_Par_1** | 0.2263 | 0.0589 | 0.0000 |
| **7Networks_RH_Default_Par_2** | 0.2045 | 0.0563 | 0.0001 |
| **7Networks_RH_Default_Par_3** | 0.1980 | 0.0561 | 0.0001 |
| **7Networks_RH_Default_Temp_1** | 0.3255 | 0.0543 | 0.0000 |
| **7Networks_RH_Default_Temp_2** | 0.3169 | 0.0556 | 0.0000 |
| **7Networks_RH_Default_Temp_3** | 0.4308 | 0.0562 | 0.0000 |
| **7Networks_RH_Default_Temp_4** | 0.2967 | 0.0533 | 0.0000 |
| **7Networks_RH_Default_Temp_5** | 0.2504 | 0.0595 | 0.0000 |
| **7Networks_RH_Default_PFCv_1** | 0.3596 | 0.0550 | 0.0000 |
| **7Networks_RH_Default_PFCm_1** | 0.2831 | 0.0532 | 0.0000 |
| **7Networks_RH_Default_PFCm_2** | 0.2488 | 0.0552 | 0.0000 |
| **7Networks_RH_Default_PFCm_3** | 0.3565 | 0.0609 | 0.0000 |
| **7Networks_RH_Default_PFCm_4** | 0.5538 | 0.0463 | 0.0000 |
| **7Networks_RH_Default_PFCm_5** | 0.4769 | 0.0474 | 0.0000 |
| **7Networks_RH_Default_PFCm_6** | 0.3159 | 0.0539 | 0.0000 |
| **7Networks_RH_Default_PFCm_7** | 0.1779 | 0.0539 | 0.0003 |
| **7Networks_RH_Default_PCC_1** | 0.3857 | 0.0577 | 0.0000 |
| **7Networks_RH_Default_PCC_2** | 0.4159 | 0.0549 | 0.0000 |
| **7Networks_RH_Default_PCC_3** | 0.2319 | 0.0547 | 0.0000 |

**Supplementary Table 5. Heritability of cortical thickness.** Heritability of cortical thickness calculated using solar 8.4.0. regions are named according to the Schaefer 200 atlas, based on 7-networks. Here we report heritability values, standard errors (SE), and p-values. All regions were significantly heritable at FDRq<0.05, with the exception of *7Networks_LH_Cont_Par_3,* which is highlighted in yellow. Nomenclature of the regions is based on the official parcel names of the Schaefer 200 – 7 networks parcel solution.

| ROI | ρ_e_ | p | ρ_g_ | p |
| --- | --- | --- | --- | --- |
| 7Networks_LH_Vis_1 | -0,079052 | 0,19729 | 0,18676 | 0,14884 |
| 7Networks_LH_Vis_2 | 0,012668 | 0,83697 | 0,063927 | 0,66213 |
| 7Networks_LH_Vis_3 | -0,065285 | 0,26767 | 0,16731 | 0,44393 |
| 7Networks_LH_Vis_4 | -0,11466 | 0,061538 | 0,2425 | 0,034865 |
| 7Networks_LH_Vis_5 | 0,075226 | 0,23203 | -0,0003708 | 0,99701 |
| 7Networks_LH_Vis_6 | -0,062598 | 0,32606 | 0,020191 | 0,87028 |
| 7Networks_LH_Vis_7 | -0,0004927 | 0,99382 | 0,1427 | 0,21894 |
| 7Networks_LH_Vis_8 | -0,057216 | 0,31688 | 0,14571 | 0,35828 |
| 7Networks_LH_Vis_9 | -0,050569 | 0,42351 | 0,28583 | 0,014601 |
| 7Networks_LH_Vis_10 | -0,18542 | 0,0039403 | 0,35907 | 0,0013726 |
| 7Networks_LH_Vis_11 | 0,055302 | 0,35692 | 0,036676 | 0,79524 |
| 7Networks_LH_Vis_12 | -0,058799 | 0,37029 | 0,15021 | 0,2271 |
| 7Networks_LH_Vis_13 | 0,054723 | 0,42117 | 0,033513 | 0,76768 |
| 7Networks_LH_Vis_14 | 0,097218 | 0,1308 | 0,026523 | 0,83438 |
| 7Networks_LH_SomMot_1 | 0,025695 | 0,69266 | 0,0049152 | 0,96785 |
| 7Networks_LH_SomMot_2 | 0,081928 | 0,17203 | 0,018946 | 0,90512 |
| 7Networks_LH_SomMot_3 | 0,031826 | 0,63064 | 0,077473 | 0,5073 |
| 7Networks_LH_SomMot_4 | 0,011344 | 0,85787 | -0,048262 | 0,71566 |
| 7Networks_LH_SomMot_5 | -0,019742 | 0,73747 | -0,025311 | 0,87739 |
| 7Networks_LH_SomMot_6 | 0,0066233 | 0,91986 | 0,12049 | 0,29016 |
| 7Networks_LH_SomMot_7 | 0,015863 | 0,79591 | -0,18177 | 0,25798 |
| 7Networks_LH_SomMot_8 | -0,033367 | 0,58906 | 0,0038403 | 0,98099 |
| 7Networks_LH_SomMot_9 | -0,048875 | 0,42583 | 0,046745 | 0,74453 |
| 7Networks_LH_SomMot_10 | 0,035033 | 0,59616 | 0,096343 | 0,41044 |
| 7Networks_LH_SomMot_11 | -0,05484 | 0,34707 | -0,0005635 | 0,99748 |
| 7Networks_LH_SomMot_12 | -0,0072036 | 0,90763 | -0,1773 | 0,21072 |
| 7Networks_LH_SomMot_13 | -0,063163 | 0,31636 | 0,039682 | 0,79395 |
| 7Networks_LH_SomMot_14 | 0,0032122 | 0,95786 | -0,12451 | 0,35643 |
| 7Networks_LH_SomMot_15 | -0,03197 | 0,6117 | 0,034414 | 0,78044 |
| 7Networks_LH_SomMot_16 | 0,059943 | 0,3715 | -0,099077 | 0,47536 |
| 7Networks_LH_DorsAttn_Post_1 | -0,05155 | 0,4141 | 0,12269 | 0,41534 |
| 7Networks_LH_DorsAttn_Post_2 | -0,083855 | 0,14368 | 0,27419 | 0,11542 |
| 7Networks_LH_DorsAttn_Post_3 | -0,059709 | 0,33232 | 0,12045 | 0,3938 |
| 7Networks_LH_DorsAttn_Post_4 | 0,06774 | 0,27799 | 0,074927 | 0,60125 |
| 7Networks_LH_DorsAttn_Post_5 | -0,024555 | 0,67318 | 0,14005 | 0,42657 |
| 7Networks_LH_DorsAttn_Post_6 | -0,025155 | 0,67756 | 0,15936 | 0,36807 |
| 7Networks_LH_DorsAttn_Post_7 | 0,011903 | 0,85534 | -0,24582 | 0,048949 |
| 7Networks_LH_DorsAttn_Post_8 | -0,072651 | 0,23102 | -0,053424 | 0,72122 |
| 7Networks_LH_DorsAttn_Post_9 | 0,063859 | 0,32607 | -0,26818 | 0,025986 |
| 7Networks_LH_DorsAttn_Post_10 | 0,026982 | 0,68308 | -0,071597 | 0,56751 |
| 7Networks_LH_DorsAttn_FEF_1 | -0,039598 | 0,50895 | -0,012733 | 0,93624 |
| 7Networks_LH_DorsAttn_FEF_2 | 0,0066935 | 0,91562 | -0,25164 | 0,073418 |
| 7Networks_LH_DorsAttn_PrCv_1 | 0,015117 | 0,79866 | -0,22564 | 0,2397 |
| 7Networks_LH_SalVentAttn_ParOper_1 | -0,0025321 | 0,96536 | 0,32611 | 0,14177 |
| 7Networks_LH_SalVentAttn_ParOper_2 | 0,032171 | 0,5844 | -0,0705 | 0,64219 |
| 7Networks_LH_SalVentAttn_ParOper_3 | 0,0063434 | 0,91621 | -0,009796 | 0,9559 |
| 7Networks_LH_SalVentAttn_FrOper_1 | 0,0027538 | 0,96426 | 0,0071485 | 0,95907 |
| 7Networks_LH_SalVentAttn_FrOper_2 | 0,040815 | 0,5263 | 0,038533 | 0,77339 |
| 7Networks_LH_SalVentAttn_FrOper_3 | -0,002797 | 0,96532 | 0,10839 | 0,39938 |
| 7Networks_LH_SalVentAttn_FrOper_4 | 0,0055092 | 0,92869 | -0,078883 | 0,6509 |
| 7Networks_LH_SalVentAttn_PFCl_1 | 0,02871 | 0,64372 | -0,1678 | 0,16588 |
| 7Networks_LH_SalVentAttn_Med_1 | 0,0555 | 0,38128 | -0,036957 | 0,80586 |
| 7Networks_LH_SalVentAttn_Med_2 | -0,046889 | 0,45594 | 0,1865 | 0,20748 |
| 7Networks_LH_SalVentAttn_Med_3 | 0,11748 | 0,051188 | -0,358 | 0,034025 |
| 7Networks_LH_Limbic_OFC_1 | 0,10849 | 0,065906 | -0,27566 | 0,062208 |
| 7Networks_LH_Limbic_OFC_2 | -0,0020436 | 0,97408 | 0,098323 | 0,49165 |
| 7Networks_LH_Limbic_TempPole_1 | 0,064778 | 0,30702 | -0,038212 | 0,79566 |
| 7Networks_LH_Limbic_TempPole_2 | -0,0096495 | 0,87326 | 0,091728 | 0,49251 |
| 7Networks_LH_Limbic_TempPole_3 | -0,0067672 | 0,90693 | 0,10621 | 0,50931 |
| 7Networks_LH_Limbic_TempPole_4 | 0,0095145 | 0,88315 | -0,0074987 | 0,95256 |
| 7Networks_LH_Cont_Par_1 | 0,014026 | 0,81303 | -0,044135 | 0,80398 |
| 7Networks_LH_Cont_Par_2 | 0,037755 | 0,52275 | -0,17838 | 0,24893 |
| 7Networks_LH_Cont_Par_3 | -0,02525 | 0,6534 | 0,37426 | 0,32587 |
| 7Networks_LH_Cont_Temp_1 | 0,084383 | 0,15001 | -0,14186 | 0,44977 |
| 7Networks_LH_Cont_PFCl_1 | 0,022654 | 0,69314 | 0,075481 | 0,62386 |
| 7Networks_LH_Cont_PFCl_2 | 0,052698 | 0,38374 | -0,22256 | 0,14458 |
| 7Networks_LH_Cont_PFCl_3 | -0,0023602 | 0,97009 | -0,044528 | 0,73868 |
| 7Networks_LH_Cont_PFCl_4 | -0,12144 | 0,040379 | -0,080271 | 0,61394 |
| 7Networks_LH_Cont_PFCl_5 | -0,041619 | 0,50252 | -0,12044 | 0,34663 |
| 7Networks_LH_Cont_PFCl_6 | -0,010749 | 0,85558 | -0,095273 | 0,60287 |
| 7Networks_LH_Cont_pCun_1 | -0,016957 | 0,78178 | -0,098956 | 0,4593 |
| 7Networks_LH_Cont_Cing_1 | -0,079363 | 0,24079 | 0,017564 | 0,881 |
| 7Networks_LH_Cont_Cing_2 | 0,0075894 | 0,90506 | -0,08263 | 0,55828 |
| 7Networks_LH_Default_Temp_1 | -0,069696 | 0,26057 | 0,25444 | 0,11437 |
| 7Networks_LH_Default_Temp_2 | 0,16512 | 0,0050416 | -0,2983 | 0,055193 |
| 7Networks_LH_Default_Temp_3 | 0,0001054 | 0,99805 | 0,19144 | 0,17682 |
| 7Networks_LH_Default_Temp_4 | 0,043015 | 0,46868 | 0,13858 | 0,4665 |
| 7Networks_LH_Default_Temp_5 | -0,15362 | 0,011314 | 0,59788 | 0,0004934 |
| 7Networks_LH_Default_Temp_6 | -0,040484 | 0,49612 | 0,18066 | 0,23753 |
| 7Networks_LH_Default_Temp_7 | -0,0061987 | 0,91969 | -0,098815 | 0,52125 |
| 7Networks_LH_Default_Temp_8 | -0,058035 | 0,33503 | 0,19464 | 0,23756 |
| 7Networks_LH_Default_Temp_9 | 0,0012862 | 0,9823 | -0,094239 | 0,54007 |
| 7Networks_LH_Default_PFC_1 | -0,026717 | 0,66231 | 0,13345 | 0,32796 |
| 7Networks_LH_Default_PFC_2 | 0,14148 | 0,019489 | -0,28872 | 0,077282 |
| 7Networks_LH_Default_PFC_3 | 0,097464 | 0,088585 | -0,39062 | 0,061042 |
| 7Networks_LH_Default_PFC_4 | 0,033147 | 0,60829 | -0,17588 | 0,12119 |
| 7Networks_LH_Default_PFC_5 | -0,10377 | 0,095584 | -0,017451 | 0,90815 |
| 7Networks_LH_Default_PFC_6 | -0,018674 | 0,75928 | -0,023309 | 0,87077 |
| 7Networks_LH_Default_PFC_7 | -0,013331 | 0,83922 | -0,22214 | 0,061045 |
| 7Networks_LH_Default_PFC_8 | -0,045787 | 0,47826 | 0,11445 | 0,41756 |
| 7Networks_LH_Default_PFC_9 | -0,048351 | 0,44487 | -0,21022 | 0,078581 |
| 7Networks_LH_Default_PFC_10 | -0,046104 | 0,45976 | -0,039388 | 0,77116 |
| 7Networks_LH_Default_PFC_11 | 0,0078701 | 0,89816 | -0,35606 | 0,0037521 |
| 7Networks_LH_Default_PFC_12 | -0,039602 | 0,53482 | -0,16614 | 0,16671 |
| 7Networks_LH_Default_PFC_13 | -0,0231 | 0,71745 | -0,2591 | 0,033329 |
| 7Networks_LH_Default_PCC_1 | -0,095588 | 0,13269 | 0,15406 | 0,27152 |
| 7Networks_LH_Default_PCC_2 | -0,099767 | 0,092117 | 0,30689 | 0,044508 |
| 7Networks_LH_Default_PCC_3 | -0,064521 | 0,30722 | 0,099199 | 0,46033 |
| 7Networks_LH_Default_PCC_4 | -0,053317 | 0,40417 | 0,12529 | 0,36301 |
| 7Networks_LH_Default_PHC_1 | 0,06492 | 0,32866 | -0,11549 | 0,36047 |
| 7Networks_RH_Vis_1 | 0,01526 | 0,80482 | 0,16498 | 0,23323 |
| 7Networks_RH_Vis_2 | -0,040559 | 0,51608 | 0,3075 | 0,0359 |
| 7Networks_RH_Vis_3 | -0,1453 | 0,021953 | 0,33979 | 0,015079 |
| 7Networks_RH_Vis_4 | -0,12988 | 0,049683 | 0,31992 | 0,0028168 |
| 7Networks_RH_Vis_5 | 0,033343 | 0,57817 | 0,013407 | 0,93125 |
| 7Networks_RH_Vis_6 | -0,0071607 | 0,91131 | 0,1408 | 0,25239 |
| 7Networks_RH_Vis_7 | -0,12116 | 0,060626 | 0,049553 | 0,66559 |
| 7Networks_RH_Vis_8 | -0,040656 | 0,52823 | 0,20938 | 0,10582 |
| 7Networks_RH_Vis_9 | -0,080138 | 0,22911 | 0,26415 | 0,016985 |
| 7Networks_RH_Vis_10 | -0,028164 | 0,67088 | 0,1684 | 0,1765 |
| 7Networks_RH_Vis_11 | 0,025049 | 0,66596 | 0,1055 | 0,52108 |
| 7Networks_RH_Vis_12 | 0,035559 | 0,59216 | 0,1063 | 0,36182 |
| 7Networks_RH_Vis_13 | 0,04067 | 0,52977 | 0,074824 | 0,53406 |
| 7Networks_RH_Vis_14 | 0,07806 | 0,22609 | -0,26922 | 0,03158 |
| 7Networks_RH_Vis_15 | 0,078028 | 0,19353 | -0,024067 | 0,87919 |
| 7Networks_RH_SomMot_1 | 0,11893 | 0,066284 | -0,090136 | 0,48669 |
| 7Networks_RH_SomMot_2 | 0,097147 | 0,12609 | -0,022086 | 0,88268 |
| 7Networks_RH_SomMot_3 | -0,0080235 | 0,89874 | 0,08856 | 0,54096 |
| 7Networks_RH_SomMot_4 | 0,029742 | 0,63395 | 0,14224 | 0,32241 |
| 7Networks_RH_SomMot_5 | 0,0098532 | 0,86777 | 0,18981 | 0,31908 |
| 7Networks_RH_SomMot_6 | 0,028825 | 0,63982 | -0,073608 | 0,63328 |
| 7Networks_RH_SomMot_7 | 0,052779 | 0,40199 | 0,14991 | 0,22949 |
| 7Networks_RH_SomMot_8 | 0,13449 | 0,019246 | -0,22895 | 0,21083 |
| 7Networks_RH_SomMot_9 | 0,010379 | 0,87072 | 0,19935 | 0,18581 |
| 7Networks_RH_SomMot_10 | 0,077136 | 0,19962 | -0,043387 | 0,83579 |
| 7Networks_RH_SomMot_11 | 0,19243 | 0,0011463 | -0,31677 | 0,031319 |
| 7Networks_RH_SomMot_12 | 0,064576 | 0,31419 | 0,075429 | 0,54191 |
| 7Networks_RH_SomMot_13 | -0,015987 | 0,78848 | 0,12894 | 0,37851 |
| 7Networks_RH_SomMot_14 | -0,034907 | 0,5747 | -0,093961 | 0,48366 |
| 7Networks_RH_SomMot_15 | -0,11791 | 0,06322 | 0,27408 | 0,059718 |
| 7Networks_RH_SomMot_16 | 0,0019755 | 0,9746 | 0,0013636 | 0,99314 |
| 7Networks_RH_SomMot_17 | -0,01335 | 0,8384 | 0,019774 | 0,86923 |
| 7Networks_RH_SomMot_18 | 0,069773 | 0,25036 | -0,088504 | 0,49984 |
| 7Networks_RH_SomMot_19 | 0,13469 | 0,038284 | -0,21184 | 0,10014 |
| 7Networks_RH_DorsAttn_Post_1 | -0,05627 | 0,34214 | 0,34532 | 0,042578 |
| 7Networks_RH_DorsAttn_Post_2 | 0,0057328 | 0,92329 | 0,13699 | 0,41797 |
| 7Networks_RH_DorsAttn_Post_3 | 0,097877 | 0,1024 | -0,054596 | 0,70936 |
| 7Networks_RH_DorsAttn_Post_4 | -0,039292 | 0,50859 | 0,11921 | 0,50315 |
| 7Networks_RH_DorsAttn_Post_5 | 0,031397 | 0,59703 | 0,15275 | 0,41654 |
| 7Networks_RH_DorsAttn_Post_6 | 0,079684 | 0,20391 | -0,21119 | 0,1034 |
| 7Networks_RH_DorsAttn_Post_7 | -0,0195 | 0,73558 | -0,02258 | 0,91652 |
| 7Networks_RH_DorsAttn_Post_8 | -0,10281 | 0,10431 | 0,22073 | 0,10718 |
| 7Networks_RH_DorsAttn_Post_9 | 0,021472 | 0,72434 | -0,11398 | 0,42314 |
| 7Networks_RH_DorsAttn_Post_10 | -0,047737 | 0,44655 | 0,061705 | 0,6305 |
| 7Networks_RH_DorsAttn_FEF_1 | -0,087941 | 0,16012 | 0,08259 | 0,58942 |
| 7Networks_RH_DorsAttn_FEF_2 | 0,059472 | 0,31442 | -0,35923 | 0,03744 |
| 7Networks_RH_DorsAttn_PrCv_1 | -0,023558 | 0,69956 | -0,085148 | 0,56629 |
| 7Networks_RH_SalVentAttn_TempOccPar_1 | -0,0030163 | 0,95955 | 0,24926 | 0,14885 |
| 7Networks_RH_SalVentAttn_TempOccPar_2 | 0,06984 | 0,22222 | -0,10182 | 0,60233 |
| 7Networks_RH_SalVentAttn_TempOccPar_3 | 0,014644 | 0,80411 | 0,12575 | 0,43957 |
| 7Networks_RH_SalVentAttn_PrC_1 | -0,032073 | 0,58046 | -0,17922 | 0,38742 |
| 7Networks_RH_SalVentAttn_FrOper_1 | 0,073802 | 0,21908 | -0,24837 | 0,12224 |
| 7Networks_RH_SalVentAttn_FrOper_2 | 0,033102 | 0,62273 | -0,058355 | 0,63411 |
| 7Networks_RH_SalVentAttn_FrOper_3 | -0,017422 | 0,77187 | 0,16148 | 0,26407 |
| 7Networks_RH_SalVentAttn_FrOper_4 | -0,047879 | 0,42419 | 0,15661 | 0,34801 |
| 7Networks_RH_SalVentAttn_Med_1 | 0,081997 | 0,17022 | -0,1724 | 0,28292 |
| 7Networks_RH_SalVentAttn_Med_2 | 0,056308 | 0,36997 | 0,0088976 | 0,94468 |
| 7Networks_RH_SalVentAttn_Med_3 | 0,18492 | 0,0028243 | -0,55335 | 0,0004109 |
| 7Networks_RH_Limbic_OFC_1 | 0,000525 | 0,99352 | 0,08355 | 0,53641 |
| 7Networks_RH_Limbic_OFC_2 | 0,016288 | 0,78382 | 0,070742 | 0,59627 |
| 7Networks_RH_Limbic_OFC_3 | 0,093422 | 0,13691 | -0,20322 | 0,13794 |
| 7Networks_RH_Limbic_TempPole_1 | 0,025827 | 0,65655 | 0,061589 | 0,74812 |
| 7Networks_RH_Limbic_TempPole_2 | -0,019268 | 0,75829 | 0,18356 | 0,19993 |
| 7Networks_RH_Limbic_TempPole_3 | 0,074456 | 0,26554 | -0,031963 | 0,78812 |
| 7Networks_RH_Cont_Par_1 | 0,013852 | 0,81185 | -0,084165 | 0,68608 |
| 7Networks_RH_Cont_Par_2 | 0,072785 | 0,21067 | -0,069621 | 0,70206 |
| 7Networks_RH_Cont_Par_3 | 0,050633 | 0,38055 | -0,18529 | 0,402 |
| 7Networks_RH_Cont_Temp_1 | -0,089824 | 0,13905 | 0,35894 | 0,05401 |
| 7Networks_RH_Cont_PFCv_1 | -0,045812 | 0,44105 | 0,19838 | 0,28585 |
| 7Networks_RH_Cont_PFCl_1 | 0,042174 | 0,47755 | 0,045826 | 0,77041 |
| 7Networks_RH_Cont_PFCl_2 | -0,014437 | 0,81156 | 0,032326 | 0,82868 |
| 7Networks_RH_Cont_PFCl_3 | -0,12094 | 0,04088 | 0,13941 | 0,32564 |
| 7Networks_RH_Cont_PFCl_4 | 0,024286 | 0,69911 | -0,059137 | 0,6466 |
| 7Networks_RH_Cont_PFCl_5 | -0,051934 | 0,38582 | -0,069729 | 0,62116 |
| 7Networks_RH_Cont_PFCl_6 | -0,0091309 | 0,87687 | -0,046093 | 0,74684 |
| 7Networks_RH_Cont_PFCl_7 | 0,01624 | 0,79713 | -0,069193 | 0,63339 |
| 7Networks_RH_Cont_pCun_1 | -0,062743 | 0,28735 | 0,17131 | 0,28731 |
| 7Networks_RH_Cont_PFCmp_1 | -0,064961 | 0,30436 | 0,039802 | 0,74008 |
| 7Networks_RH_Cont_PFCmp_2 | 0,048745 | 0,44644 | -0,14787 | 0,25835 |
| 7Networks_RH_Cont_PFCmp_3 | 0,14451 | 0,016821 | -0,48906 | 0,0023123 |
| 7Networks_RH_Cont_PFCmp_4 | 0,046628 | 0,47516 | -0,30197 | 0,019916 |
| 7Networks_RH_Default_Par_1 | 0,095142 | 0,11799 | -0,075421 | 0,66797 |
| 7Networks_RH_Default_Par_2 | 0,044959 | 0,44645 | 0,059015 | 0,74643 |
| 7Networks_RH_Default_Par_3 | -0,026585 | 0,64961 | -0,062762 | 0,73235 |
| 7Networks_RH_Default_Temp_1 | 0,028678 | 0,63731 | -0,12972 | 0,36886 |
| 7Networks_RH_Default_Temp_2 | -0,03071 | 0,61712 | 0,1155 | 0,43617 |
| 7Networks_RH_Default_Temp_3 | -0,052706 | 0,42281 | 0,16048 | 0,23618 |
| 7Networks_RH_Default_Temp_4 | -0,094782 | 0,10756 | 0,28412 | 0,063133 |
| 7Networks_RH_Default_Temp_5 | 0,12176 | 0,047406 | -0,049152 | 0,77162 |
| 7Networks_RH_Default_PFCv_1 | -0,063573 | 0,30782 | 0,015555 | 0,91186 |
| 7Networks_RH_Default_PFCm_1 | 0,087377 | 0,13796 | -0,09607 | 0,52614 |
| 7Networks_RH_Default_PFCm_2 | 0,025903 | 0,66185 | -0,026321 | 0,87224 |
| 7Networks_RH_Default_PFCm_3 | -0,047054 | 0,47151 | 0,0038185 | 0,97947 |
| 7Networks_RH_Default_PFCm_4 | 0,003877 | 0,95457 | -0,13584 | 0,22847 |
| 7Networks_RH_Default_PFCm_5 | 0,029688 | 0,64304 | -0,33171 | 0,0049967 |
| 7Networks_RH_Default_PFCm_6 | 0,21399 | 0,0003541 | -0,50618 | 0,0005788 |
| 7Networks_RH_Default_PFCm_7 | -0,0080771 | 0,88762 | -0,12891 | 0,49347 |
| 7Networks_RH_Default_PCC_1 | -0,12127 | 0,057177 | 0,22042 | 0,12134 |
| 7Networks_RH_Default_PCC_2 | -0,035063 | 0,5869 | 0,076935 | 0,56602 |
| 7Networks_RH_Default_PCC_3 | 0,082602 | 0,15637 | -0,076839 | 0,64896 |

**Supplementary Table 6. Genetic and environmental correlation between Agreeableness and cortical thickness in HCP.** Genetic correlation between Agreeableness and local cortical thickness calculated using solar 8.4.0. Regions are named according to the Schaefer 200 atlas, based on 7-networks. Here we report environmental correlation (ρ_e_) and genetic correlation (ρ_g_) and the associated p-values.

| ROI name | ρ_e_ | p | ρ_g_ | p |
| --- | --- | --- | --- | --- |
| 7Networks_LH_Vis_1 | -0.077319 | 0.21739 | 0.031867 | 0.76213 |
| 7Networks_LH_Vis_2 | 0.020231 | 0.74679 | -0.021573 | 0.85557 |
| 7Networks_LH_Vis_3 | -0.019318 | 0.74977 | -0.16548 | 0.33899 |
| 7Networks_LH_Vis_4 | 0.030785 | 0.62362 | -0.037492 | 0.68262 |
| 7Networks_LH_Vis_5 | -0.045422 | 0.47837 | -0.0060919 | 0.95433 |
| 7Networks_LH_Vis_6 | 0.0020273 | 0.975 | -0.0084422 | 0.93314 |
| 7Networks_LH_Vis_7 | 0.018961 | 0.77164 | 0.023085 | 0.80733 |
| 7Networks_LH_Vis_8 | 0.061134 | 0.29104 | -0.10968 | 0.40093 |
| 7Networks_LH_Vis_9 | -0.052112 | 0.42324 | 0.086626 | 0.36478 |
| 7Networks_LH_Vis_10 | -0.0001018 | 0.99887 | -0.023673 | 0.7997 |
| 7Networks_LH_Vis_11 | 0.019588 | 0.75194 | 0.11301 | 0.32772 |
| 7Networks_LH_Vis_12 | -0.010871 | 0.87054 | -0.017826 | 0.85943 |
| 7Networks_LH_Vis_13 | 0.051006 | 0.45749 | 0.021795 | 0.813 |
| 7Networks_LH_Vis_14 | 0.097985 | 0.13338 | -0.013031 | 0.89892 |
| 7Networks_LH_SomMot_1 | -0.022766 | 0.72929 | -0.0059826 | 0.95069 |
| 7Networks_LH_SomMot_2 | -0.0047647 | 0.93819 | 0.037657 | 0.76946 |
| 7Networks_LH_SomMot_3 | -0.034172 | 0.60967 | -0.092592 | 0.32895 |
| 7Networks_LH_SomMot_4 | -0.040704 | 0.52632 | -0.047306 | 0.66143 |
| 7Networks_LH_SomMot_5 | -0.09301 | 0.12396 | 0.12071 | 0.36274 |
| 7Networks_LH_SomMot_6 | -0.061314 | 0.35693 | 0.0008657 | 0.99243 |
| 7Networks_LH_SomMot_7 | 0.051566 | 0.41174 | -0.0005182 | 0.99643 |
| 7Networks_LH_SomMot_8 | -0.064761 | 0.30999 | 0.18022 | 0.1665 |
| 7Networks_LH_SomMot_9 | 0.067279 | 0.28354 | -0.14346 | 0.21244 |
| 7Networks_LH_SomMot_10 | -0.0019795 | 0.97631 | -0.1657 | 0.079732 |
| 7Networks_LH_SomMot_11 | 0.045638 | 0.44984 | -0.11944 | 0.39923 |
| 7Networks_LH_SomMot_12 | 0.030527 | 0.62916 | -0.0062447 | 0.95609 |
| 7Networks_LH_SomMot_13 | 0.014934 | 0.81601 | -0.019287 | 0.87374 |
| 7Networks_LH_SomMot_14 | -0.062096 | 0.31446 | 0.063622 | 0.56204 |
| 7Networks_LH_SomMot_15 | -0.014165 | 0.82361 | -0.087766 | 0.38355 |
| 7Networks_LH_SomMot_16 | 0.10419 | 0.11947 | -0.13423 | 0.22497 |
| 7Networks_LH_DorsAttn_Post_1 | 0.16223 | 0.0097636 | -0.41593 | 0.0006824 |
| 7Networks_LH_DorsAttn_Post_2 | 0.0079147 | 0.8933 | -0.028017 | 0.8417 |
| 7Networks_LH_DorsAttn_Post_3 | -0.0016595 | 0.97895 | 0.052246 | 0.65202 |
| 7Networks_LH_DorsAttn_Post_4 | 0.018212 | 0.77597 | -0.045804 | 0.69585 |
| 7Networks_LH_DorsAttn_Post_5 | 0.0090312 | 0.88023 | 0.046217 | 0.74586 |
| 7Networks_LH_DorsAttn_Post_6 | -0.050747 | 0.41443 | 0.14445 | 0.31342 |
| 7Networks_LH_DorsAttn_Post_7 | -0.062537 | 0.34788 | 0.061882 | 0.54495 |
| 7Networks_LH_DorsAttn_Post_8 | -0.082318 | 0.18385 | 0.11137 | 0.36049 |
| 7Networks_LH_DorsAttn_Post_9 | -0.033359 | 0.60768 | 0.012096 | 0.90459 |
| 7Networks_LH_DorsAttn_Post_10 | 0.0044846 | 0.94713 | -0.023911 | 0.81669 |
| 7Networks_LH_DorsAttn_FEF_1 | -0.11762 | 0.054424 | 0.27643 | 0.029121 |
| 7Networks_LH_DorsAttn_FEF_2 | 0.037359 | 0.56248 | 0.0037484 | 0.974 |
| 7Networks_LH_DorsAttn_PrCv_1 | -0.0906 | 0.132 | 0.35233 | 0.026929 |
| 7Networks_LH_SalVentAttn_ParOper_1 | -0.082851 | 0.16615 | 0.13334 | 0.47023 |
| 7Networks_LH_SalVentAttn_ParOper_2 | -0.0074933 | 0.90029 | -0.11113 | 0.36781 |
| 7Networks_LH_SalVentAttn_ParOper_3 | -0.0144 | 0.81558 | 0.011076 | 0.93966 |
| 7Networks_LH_SalVentAttn_FrOper_1 | -0.068857 | 0.27057 | -0.045872 | 0.68752 |
| 7Networks_LH_SalVentAttn_FrOper_2 | 0.04249 | 0.5115 | -0.1597 | 0.13611 |
| 7Networks_LH_SalVentAttn_FrOper_3 | 0.07889 | 0.22347 | -0.33271 | 0.0013496 |
| 7Networks_LH_SalVentAttn_FrOper_4 | -0.016992 | 0.78685 | 0.20297 | 0.15611 |
| 7Networks_LH_SalVentAttn_PFCl_1 | -0.03721 | 0.55386 | 0.17134 | 0.079832 |
| 7Networks_LH_SalVentAttn_Med_1 | -0.054133 | 0.40252 | 0.2731 | 0.025194 |
| 7Networks_LH_SalVentAttn_Med_2 | 0.046377 | 0.46887 | -0.01192 | 0.92117 |
| 7Networks_LH_SalVentAttn_Med_3 | 0.1825 | 0.0031986 | -0.12789 | 0.35265 |
| 7Networks_LH_Limbic_OFC_1 | 0.026823 | 0.65874 | 0.063344 | 0.60155 |
| 7Networks_LH_Limbic_OFC_2 | 0.017305 | 0.78904 | -0.016377 | 0.88751 |
| 7Networks_LH_Limbic_TempPole_1 | -0.039704 | 0.53917 | 0.038064 | 0.75364 |
| 7Networks_LH_Limbic_TempPole_2 | -0.058372 | 0.3406 | 0.021006 | 0.8443 |
| 7Networks_LH_Limbic_TempPole_3 | -0.011536 | 0.84501 | -0.11831 | 0.36935 |
| 7Networks_LH_Limbic_TempPole_4 | -0.037723 | 0.56597 | 0.029318 | 0.77513 |
| 7Networks_LH_Cont_Par_1 | -0.027188 | 0.6545 | 0.084068 | 0.56199 |
| 7Networks_LH_Cont_Par_2 | -0.12694 | 0.033798 | 0.16236 | 0.20299 |
| 7Networks_LH_Cont_Par_3 | -0.053117 | 0.35935 | 0.18933 | 0.53464 |
| 7Networks_LH_Cont_Temp_1 | 0.055774 | 0.35649 | 0.0015069 | 0.99194 |
| 7Networks_LH_Cont_PFCl_1 | 0.018594 | 0.75069 | 0.023251 | 0.85269 |
| 7Networks_LH_Cont_PFCl_2 | 0.0060743 | 0.92192 | -0.084399 | 0.49749 |
| 7Networks_LH_Cont_PFCl_3 | 0.0021952 | 0.97253 | 0.080574 | 0.45774 |
| 7Networks_LH_Cont_PFCl_4 | -0.09488 | 0.12077 | 0.22866 | 0.070802 |
| 7Networks_LH_Cont_PFCl_5 | -0.001802 | 0.9771 | 0.21568 | 0.035678 |
| 7Networks_LH_Cont_PFCl_6 | -0.043499 | 0.4709 | 0.26095 | 0.075109 |
| 7Networks_LH_Cont_pCun_1 | -0.0087962 | 0.88796 | 0.016218 | 0.88139 |
| 7Networks_LH_Cont_Cing_1 | -0.063993 | 0.35004 | 0.070036 | 0.46001 |
| 7Networks_LH_Cont_Cing_2 | 0.034948 | 0.5915 | -0.13962 | 0.2225 |
| 7Networks_LH_Default_Temp_1 | -0.010139 | 0.87328 | 0.044788 | 0.73194 |
| 7Networks_LH_Default_Temp_2 | 0.098493 | 0.10614 | -0.036615 | 0.77177 |
| 7Networks_LH_Default_Temp_3 | 0.01818 | 0.77103 | -0.095804 | 0.40463 |
| 7Networks_LH_Default_Temp_4 | -0.029639 | 0.62657 | -0.020403 | 0.89475 |
| 7Networks_LH_Default_Temp_5 | -0.091016 | 0.13879 | -0.041834 | 0.77842 |
| 7Networks_LH_Default_Temp_6 | 0.031056 | 0.60835 | -0.099433 | 0.42615 |
| 7Networks_LH_Default_Temp_7 | -0.023026 | 0.71442 | 0.015706 | 0.90017 |
| 7Networks_LH_Default_Temp_8 | -0.082184 | 0.17506 | 0.21425 | 0.12269 |
| 7Networks_LH_Default_Temp_9 | -0.014934 | 0.80074 | 0.03788 | 0.76324 |
| 7Networks_LH_Default_PFC_1 | 0.024343 | 0.69604 | -0.14245 | 0.20001 |
| 7Networks_LH_Default_PFC_2 | -0.056921 | 0.36862 | 0.22041 | 0.091025 |
| 7Networks_LH_Default_PFC_3 | -0.036084 | 0.53982 | -0.1038 | 0.54295 |
| 7Networks_LH_Default_PFC_4 | -0.019404 | 0.76718 | 0.034188 | 0.71211 |
| 7Networks_LH_Default_PFC_5 | -0.10294 | 0.10452 | 0.19727 | 0.10704 |
| 7Networks_LH_Default_PFC_6 | -0.064973 | 0.29716 | 0.15564 | 0.17843 |
| 7Networks_LH_Default_PFC_7 | -0.057612 | 0.38418 | 0.13196 | 0.16699 |
| 7Networks_LH_Default_PFC_8 | 0.043272 | 0.51197 | -0.02326 | 0.83834 |
| 7Networks_LH_Default_PFC_9 | -0.042367 | 0.50986 | 0.043037 | 0.66124 |
| 7Networks_LH_Default_PFC_10 | 0.0052061 | 0.93367 | 0.04063 | 0.7078 |
| 7Networks_LH_Default_PFC_11 | 0.0036174 | 0.9536 | 0.18172 | 0.069187 |
| 7Networks_LH_Default_PFC_12 | -0.0028771 | 0.96396 | 0.012089 | 0.90156 |
| 7Networks_LH_Default_PFC_13 | -0.056134 | 0.38584 | 0.09851 | 0.31969 |
| 7Networks_LH_Default_PCC_1 | -0.055093 | 0.3967 | 0.060109 | 0.58533 |
| 7Networks_LH_Default_PCC_2 | -0.030578 | 0.61521 | 0.078073 | 0.53121 |
| 7Networks_LH_Default_PCC_3 | -0.037161 | 0.56301 | 0.037023 | 0.73463 |
| 7Networks_LH_Default_PCC_4 | -0.010899 | 0.8678 | -0.046364 | 0.67889 |
| 7Networks_LH_Default_PHC_1 | -0.08159 | 0.22444 | 0.073602 | 0.48017 |
| 7Networks_RH_Vis_1 | 0.057201 | 0.36663 | -0.10737 | 0.33181 |
| 7Networks_RH_Vis_2 | -0.033924 | 0.5973 | 0.038809 | 0.74153 |
| 7Networks_RH_Vis_3 | 0.046263 | 0.47867 | -0.11764 | 0.30064 |
| 7Networks_RH_Vis_4 | -0.16016 | 0.018567 | 0.13707 | 0.10772 |
| 7Networks_RH_Vis_5 | 0.058668 | 0.3427 | -0.1222 | 0.32994 |
| 7Networks_RH_Vis_6 | 0.014021 | 0.83055 | -0.068484 | 0.49273 |
| 7Networks_RH_Vis_7 | -0.022471 | 0.73405 | 0.025383 | 0.78663 |
| 7Networks_RH_Vis_8 | -0.037437 | 0.57282 | -0.022487 | 0.83179 |
| 7Networks_RH_Vis_9 | -0.047876 | 0.48225 | 0.092151 | 0.29541 |
| 7Networks_RH_Vis_10 | 0.016975 | 0.80232 | -0.042458 | 0.67763 |
| 7Networks_RH_Vis_11 | 0.054493 | 0.35708 | -0.067285 | 0.61478 |
| 7Networks_RH_Vis_12 | 0.014776 | 0.82598 | 0.010443 | 0.91209 |
| 7Networks_RH_Vis_13 | -0.031754 | 0.62904 | -0.031339 | 0.74954 |
| 7Networks_RH_Vis_14 | 0.012977 | 0.8418 | -0.14005 | 0.17876 |
| 7Networks_RH_Vis_15 | 0.026938 | 0.66212 | -0.068086 | 0.59664 |
| 7Networks_RH_SomMot_1 | 0.037501 | 0.56873 | -0.086867 | 0.40558 |
| 7Networks_RH_SomMot_2 | 0.030815 | 0.63649 | -0.044098 | 0.71403 |
| 7Networks_RH_SomMot_3 | 0.11773 | 0.066815 | -0.28746 | 0.014132 |
| 7Networks_RH_SomMot_4 | 0.017257 | 0.78505 | -0.020402 | 0.86196 |
| 7Networks_RH_SomMot_5 | -0.04032 | 0.5074 | -0.0014048 | 0.99269 |
| 7Networks_RH_SomMot_6 | 0.045523 | 0.46724 | -0.11774 | 0.35075 |
| 7Networks_RH_SomMot_7 | -0.06929 | 0.27877 | -0.08431 | 0.40853 |
| 7Networks_RH_SomMot_8 | 0.031813 | 0.5918 | 0.034054 | 0.82033 |
| 7Networks_RH_SomMot_9 | 0.027008 | 0.68193 | -0.038093 | 0.75679 |
| 7Networks_RH_SomMot_10 | 0.040142 | 0.51509 | -0.007268 | 0.96512 |
| 7Networks_RH_SomMot_11 | 0.04506 | 0.45817 | 0.11781 | 0.32939 |
| 7Networks_RH_SomMot_12 | 0.0006033 | 0.99252 | -0.11943 | 0.23107 |
| 7Networks_RH_SomMot_13 | 0.01692 | 0.78108 | -0.022586 | 0.84963 |
| 7Networks_RH_SomMot_14 | 0.12475 | 0.048685 | -0.14817 | 0.17146 |
| 7Networks_RH_SomMot_15 | 0.015389 | 0.81403 | 0.025695 | 0.83007 |
| 7Networks_RH_SomMot_16 | 0.017261 | 0.78542 | 0.060817 | 0.64336 |
| 7Networks_RH_SomMot_17 | 0.063942 | 0.3294 | -0.26003 | 0.0081624 |
| 7Networks_RH_SomMot_18 | 0.06379 | 0.30556 | -0.078342 | 0.46215 |
| 7Networks_RH_SomMot_19 | -0.012906 | 0.84477 | -0.049674 | 0.63496 |
| 7Networks_RH_DorsAttn_Post_1 | -0.028847 | 0.63632 | -0.014462 | 0.91569 |
| 7Networks_RH_DorsAttn_Post_2 | 0.065433 | 0.28324 | -0.29619 | 0.02782 |
| 7Networks_RH_DorsAttn_Post_3 | 0.069326 | 0.25789 | -0.14135 | 0.23337 |
| 7Networks_RH_DorsAttn_Post_4 | 0.10875 | 0.074748 | -0.21554 | 0.13004 |
| 7Networks_RH_DorsAttn_Post_5 | 0.021496 | 0.72454 | -0.025668 | 0.86743 |
| 7Networks_RH_DorsAttn_Post_6 | -0.013621 | 0.83005 | 0.0036559 | 0.97039 |
| 7Networks_RH_DorsAttn_Post_7 | 0.012981 | 0.82677 | -0.037657 | 0.82965 |
| 7Networks_RH_DorsAttn_Post_8 | -0.0084823 | 0.8934 | 0.038211 | 0.73724 |
| 7Networks_RH_DorsAttn_Post_9 | -0.038642 | 0.53678 | 0.03108 | 0.78929 |
| 7Networks_RH_DorsAttn_Post_10 | 0.06167 | 0.33431 | -0.043208 | 0.67943 |
| 7Networks_RH_DorsAttn_FEF_1 | 0.05782 | 0.36785 | -0.049904 | 0.69078 |
| 7Networks_RH_DorsAttn_FEF_2 | 0.064173 | 0.28964 | -0.13362 | 0.33989 |
| 7Networks_RH_DorsAttn_PrCv_1 | -0.012223 | 0.84537 | 0.065481 | 0.58746 |
| 7Networks_RH_SalVentAttn_TempOccPar_1 | 0.034713 | 0.56938 | -0.22452 | 0.11504 |
| 7Networks_RH_SalVentAttn_TempOccPar_2 | 0.0035157 | 0.95216 | 0.044896 | 0.77571 |
| 7Networks_RH_SalVentAttn_TempOccPar_3 | 0.0049294 | 0.93459 | -0.11048 | 0.40572 |
| 7Networks_RH_SalVentAttn_PrC_1 | -0.027791 | 0.64229 | 0.11681 | 0.49596 |
| 7Networks_RH_SalVentAttn_FrOper_1 | -0.001387 | 0.98184 | 0.011737 | 0.92829 |
| 7Networks_RH_SalVentAttn_FrOper_2 | 0.063812 | 0.35037 | -0.020277 | 0.83769 |
| 7Networks_RH_SalVentAttn_FrOper_3 | -0.0069686 | 0.90936 | -0.026862 | 0.81893 |
| 7Networks_RH_SalVentAttn_FrOper_4 | -0.015541 | 0.80051 | -0.16973 | 0.20346 |
| 7Networks_RH_SalVentAttn_Med_1 | -0.05008 | 0.41361 | 0.1985 | 0.12697 |
| 7Networks_RH_SalVentAttn_Med_2 | -0.0030423 | 0.96217 | 0.055779 | 0.59487 |
| 7Networks_RH_SalVentAttn_Med_3 | 0.029224 | 0.65494 | 0.0013009 | 0.99141 |
| 7Networks_RH_Limbic_OFC_1 | -0.046644 | 0.48393 | 0.14545 | 0.18315 |
| 7Networks_RH_Limbic_OFC_2 | 0.041184 | 0.49742 | -0.012827 | 0.90721 |
| 7Networks_RH_Limbic_OFC_3 | 0.058736 | 0.35898 | -0.0058573 | 0.95804 |
| 7Networks_RH_Limbic_TempPole_1 | 0.020697 | 0.72865 | -0.16974 | 0.27194 |
| 7Networks_RH_Limbic_TempPole_2 | -0.039484 | 0.53512 | 0.021472 | 0.85333 |
| 7Networks_RH_Limbic_TempPole_3 | -0.064371 | 0.33849 | 0.05844 | 0.54706 |
| 7Networks_RH_Cont_Par_1 | 0.0052287 | 0.93065 | 0.018464 | 0.91404 |
| 7Networks_RH_Cont_Par_2 | 0.11953 | 0.044264 | -0.107 | 0.47125 |
| 7Networks_RH_Cont_Par_3 | -0.0095611 | 0.87139 | 0.027634 | 0.87846 |
| 7Networks_RH_Cont_Temp_1 | -0.0224 | 0.71882 | 0.26487 | 0.072585 |
| 7Networks_RH_Cont_PFCv_1 | 0.025652 | 0.67507 | -0.034168 | 0.81645 |
| 7Networks_RH_Cont_PFCl_1 | 0.10067 | 0.095965 | -0.048468 | 0.70801 |
| 7Networks_RH_Cont_PFCl_2 | 0.045089 | 0.46641 | -0.053835 | 0.65743 |
| 7Networks_RH_Cont_PFCl_3 | 0.0065973 | 0.91339 | 0.033812 | 0.76753 |
| 7Networks_RH_Cont_PFCl_4 | 0.05794 | 0.36176 | 0.13494 | 0.19301 |
| 7Networks_RH_Cont_PFCl_5 | -0.018524 | 0.76053 | 0.072774 | 0.52668 |
| 7Networks_RH_Cont_PFCl_6 | 0.037125 | 0.5393 | -0.12707 | 0.2743 |
| 7Networks_RH_Cont_PFCl_7 | 0.037633 | 0.5587 | 0.11287 | 0.33879 |
| 7Networks_RH_Cont_pCun_1 | -0.060723 | 0.31192 | 0.21667 | 0.095314 |
| 7Networks_RH_Cont_PFCmp_1 | -0.034261 | 0.59357 | 0.098281 | 0.31428 |
| 7Networks_RH_Cont_PFCmp_2 | 0.0073659 | 0.91043 | -0.072901 | 0.49474 |
| 7Networks_RH_Cont_PFCmp_3 | 0.032848 | 0.59758 | -0.109 | 0.40345 |
| 7Networks_RH_Cont_PFCmp_4 | -0.0082365 | 0.9027 | 0.049384 | 0.63757 |
| 7Networks_RH_Default_Par_1 | 0.04386 | 0.47892 | -0.1613 | 0.26073 |
| 7Networks_RH_Default_Par_2 | 0.10623 | 0.077322 | -0.16732 | 0.2688 |
| 7Networks_RH_Default_Par_3 | 0.017001 | 0.77731 | -0.043425 | 0.7711 |
| 7Networks_RH_Default_Temp_1 | -0.057436 | 0.35624 | -0.030608 | 0.79545 |
| 7Networks_RH_Default_Temp_2 | -0.019443 | 0.75644 | 0.013218 | 0.91278 |
| 7Networks_RH_Default_Temp_3 | 0.086733 | 0.19513 | -0.084617 | 0.43547 |
| 7Networks_RH_Default_Temp_4 | -0.041545 | 0.49468 | -0.10176 | 0.40342 |
| 7Networks_RH_Default_Temp_5 | 0.04861 | 0.44075 | -0.27408 | 0.048471 |
| 7Networks_RH_Default_PFCv_1 | -0.083913 | 0.19125 | 0.12837 | 0.255 |
| 7Networks_RH_Default_PFCm_1 | -0.054706 | 0.36494 | 0.27425 | 0.02385 |
| 7Networks_RH_Default_PFCm_2 | 0.058903 | 0.32953 | 0.11629 | 0.38274 |
| 7Networks_RH_Default_PFCm_3 | 0.02927 | 0.65944 | -0.077827 | 0.51683 |
| 7Networks_RH_Default_PFCm_4 | -0.017048 | 0.80289 | 0.058069 | 0.52556 |
| 7Networks_RH_Default_PFCm_5 | 0.064147 | 0.3257 | -0.019145 | 0.84249 |
| 7Networks_RH_Default_PFCm_6 | 0.18828 | 0.0021595 | -0.046963 | 0.69211 |
| 7Networks_RH_Default_PFCm_7 | 0.030791 | 0.59863 | 0.02853 | 0.85299 |
| 7Networks_RH_Default_PCC_1 | 0.0070305 | 0.91539 | -0.092366 | 0.4163 |
| 7Networks_RH_Default_PCC_2 | -0.084853 | 0.19643 | 0.15324 | 0.15821 |
| 7Networks_RH_Default_PCC_3 | 0.053687 | 0.36939 | 0.045433 | 0.74041 |

**Supplementary Table 7. Genetic and environmental correlation between Conscientiousness and cortical thickness in HCP.** Genetic correlation between Conscientiousness and local cortical thickness calculated using solar 8.4.0. Regions are named according to the Schaefer 200 atlas, based on 7-networks. Here we report environmental correlation (ρ_e_) and genetic correlation (ρ_g_) and the associated p-values.

| ROI name | ρ_e_ | p | ρ_g_ | p |
| --- | --- | --- | --- | --- |
| 7Networks_LH_Vis_1 | -0.064948 | 0.28488 | 0.1025 | 0.31847 |
| 7Networks_LH_Vis_2 | -0.02984 | 0.62517 | 0.019301 | 0.86777 |
| 7Networks_LH_Vis_3 | -0.04565 | 0.44006 | 0.11881 | 0.479 |
| 7Networks_LH_Vis_4 | -0.054817 | 0.36709 | 0.062505 | 0.48675 |
| 7Networks_LH_Vis_5 | -0.066132 | 0.28586 | 0.049089 | 0.63462 |
| 7Networks_LH_Vis_6 | 0.030085 | 0.63056 | -0.10719 | 0.27071 |
| 7Networks_LH_Vis_7 | 0.10483 | 0.095888 | -0.17563 | 0.054484 |
| 7Networks_LH_Vis_8 | -0.037894 | 0.50597 | 0.11199 | 0.37415 |
| 7Networks_LH_Vis_9 | 0.02718 | 0.66247 | 0.048604 | 0.60027 |
| 7Networks_LH_Vis_10 | 0.006716 | 0.91632 | -0.039026 | 0.66474 |
| 7Networks_LH_Vis_11 | 0.011557 | 0.84615 | 0.11806 | 0.28916 |
| 7Networks_LH_Vis_12 | 0.038152 | 0.55275 | 0.056155 | 0.56363 |
| 7Networks_LH_Vis_13 | 0.03894 | 0.55583 | 0.083177 | 0.35067 |
| 7Networks_LH_Vis_14 | -0.0097543 | 0.87756 | 0.040093 | 0.68631 |
| 7Networks_LH_SomMot_1 | -0.057818 | 0.36342 | -0.012428 | 0.89643 |
| 7Networks_LH_SomMot_2 | 0.022158 | 0.71025 | 0.0030509 | 0.9803 |
| 7Networks_LH_SomMot_3 | 0.013214 | 0.83743 | -0.075175 | 0.41197 |
| 7Networks_LH_SomMot_4 | -0.038026 | 0.54074 | -0.052692 | 0.61323 |
| 7Networks_LH_SomMot_5 | -0.042224 | 0.47148 | -0.011526 | 0.92927 |
| 7Networks_LH_SomMot_6 | -0.023461 | 0.71585 | 0.054897 | 0.53905 |
| 7Networks_LH_SomMot_7 | 0.022402 | 0.7139 | -0.11424 | 0.35455 |
| 7Networks_LH_SomMot_8 | -0.015646 | 0.79978 | 0.16104 | 0.20094 |
| 7Networks_LH_SomMot_9 | 0.021912 | 0.71896 | 0.044365 | 0.69357 |
| 7Networks_LH_SomMot_10 | 0.040408 | 0.5338 | 0.0018957 | 0.98346 |
| 7Networks_LH_SomMot_11 | -0.031879 | 0.5857 | 0.17057 | 0.21643 |
| 7Networks_LH_SomMot_12 | 0.02527 | 0.68136 | -0.046722 | 0.67003 |
| 7Networks_LH_SomMot_13 | -0.046903 | 0.4495 | 0.14179 | 0.23077 |
| 7Networks_LH_SomMot_14 | 0.024183 | 0.68735 | -0.055121 | 0.60424 |
| 7Networks_LH_SomMot_15 | -0.034015 | 0.58054 | 0.051024 | 0.60102 |
| 7Networks_LH_SomMot_16 | 0.059599 | 0.35429 | 0.0059328 | 0.95628 |
| 7Networks_LH_DorsAttn_Post_1 | 0.012847 | 0.83681 | 0.0024396 | 0.98349 |
| 7Networks_LH_DorsAttn_Post_2 | -0.082169 | 0.15444 | 0.21064 | 0.12309 |
| 7Networks_LH_DorsAttn_Post_3 | -0.13785 | 0.026164 | 0.18714 | 0.089026 |
| 7Networks_LH_DorsAttn_Post_4 | 0.089721 | 0.14692 | -0.079131 | 0.48982 |
| 7Networks_LH_DorsAttn_Post_5 | -0.020779 | 0.72154 | 0.14512 | 0.29471 |
| 7Networks_LH_DorsAttn_Post_6 | 0.048387 | 0.42126 | -0.067583 | 0.62783 |
| 7Networks_LH_DorsAttn_Post_7 | -0.062047 | 0.33429 | 0.061672 | 0.53802 |
| 7Networks_LH_DorsAttn_Post_8 | -0.016782 | 0.78155 | 0.18563 | 0.115 |
| 7Networks_LH_DorsAttn_Post_9 | -0.018915 | 0.76367 | -0.012598 | 0.89756 |
| 7Networks_LH_DorsAttn_Post_10 | 0.031742 | 0.62404 | -0.0054248 | 0.95587 |
| 7Networks_LH_DorsAttn_FEF_1 | 0.0012074 | 0.98385 | -0.03539 | 0.77955 |
| 7Networks_LH_DorsAttn_FEF_2 | 0.051657 | 0.40508 | -0.23482 | 0.035912 |
| 7Networks_LH_DorsAttn_PrCv_1 | 0.071308 | 0.22766 | 0.057345 | 0.70572 |
| 7Networks_LH_SalVentAttn_ParOper_1 | -0.048551 | 0.40677 | 0.44971 | 0.011622 |
| 7Networks_LH_SalVentAttn_ParOper_2 | 0.011054 | 0.85074 | 0.07777 | 0.51737 |
| 7Networks_LH_SalVentAttn_ParOper_3 | 0.035949 | 0.5466 | -0.22066 | 0.1244 |
| 7Networks_LH_SalVentAttn_FrOper_1 | 0.027079 | 0.65782 | -0.13815 | 0.20704 |
| 7Networks_LH_SalVentAttn_FrOper_2 | 0.0060741 | 0.92349 | -0.068191 | 0.51249 |
| 7Networks_LH_SalVentAttn_FrOper_3 | 0.082737 | 0.18921 | -0.15307 | 0.13103 |
| 7Networks_LH_SalVentAttn_FrOper_4 | -0.05578 | 0.36937 | 0.22878 | 0.087082 |
| 7Networks_LH_SalVentAttn_PFCl_1 | 0.036465 | 0.55057 | -0.080429 | 0.40039 |
| 7Networks_LH_SalVentAttn_Med_1 | 0.06556 | 0.29499 | 0.039304 | 0.73709 |
| 7Networks_LH_SalVentAttn_Med_2 | -0.029269 | 0.63817 | 0.20471 | 0.076651 |
| 7Networks_LH_SalVentAttn_Med_3 | 0.18241 | 0.0023486 | -0.18904 | 0.15321 |
| 7Networks_LH_Limbic_OFC_1 | 0.016641 | 0.7784 | -0.057306 | 0.62549 |
| 7Networks_LH_Limbic_OFC_2 | 0.020569 | 0.73956 | -0.066454 | 0.54852 |
| 7Networks_LH_Limbic_TempPole_1 | -0.020167 | 0.7478 | -0.049579 | 0.67028 |
| 7Networks_LH_Limbic_TempPole_2 | -0.068709 | 0.2488 | 0.12783 | 0.21985 |
| 7Networks_LH_Limbic_TempPole_3 | 0.035139 | 0.54352 | -0.24495 | 0.051972 |
| 7Networks_LH_Limbic_TempPole_4 | -0.08702 | 0.17095 | 0.001956 | 0.98381 |
| 7Networks_LH_Cont_Par_1 | -0.049678 | 0.40022 | 0.20071 | 0.15089 |
| 7Networks_LH_Cont_Par_2 | 0.007694 | 0.8956 | 0.22045 | 0.071016 |
| 7Networks_LH_Cont_Par_3 | 0.07245 | 0.203 | -0.19127 | 0.51629 |
| 7Networks_LH_Cont_Temp_1 | -0.049154 | 0.40271 | 0.27491 | 0.063131 |
| 7Networks_LH_Cont_PFCl_1 | -0.046553 | 0.41537 | 0.09774 | 0.42352 |
| 7Networks_LH_Cont_PFCl_2 | -0.010549 | 0.86113 | -0.016505 | 0.89088 |
| 7Networks_LH_Cont_PFCl_3 | 0.042447 | 0.49671 | -0.17574 | 0.092612 |
| 7Networks_LH_Cont_PFCl_4 | -0.028096 | 0.63689 | -0.043983 | 0.72709 |
| 7Networks_LH_Cont_PFCl_5 | 0.0063653 | 0.9169 | -0.095556 | 0.34152 |
| 7Networks_LH_Cont_PFCl_6 | 0.014141 | 0.81043 | -0.19224 | 0.1834 |
| 7Networks_LH_Cont_pCun_1 | -0.066858 | 0.26904 | 0.092518 | 0.38309 |
| 7Networks_LH_Cont_Cing_1 | -0.11147 | 0.091664 | 0.077725 | 0.39249 |
| 7Networks_LH_Cont_Cing_2 | 0.086283 | 0.16925 | -0.13526 | 0.22553 |
| 7Networks_LH_Default_Temp_1 | -0.04993 | 0.41833 | 0.086754 | 0.49008 |
| 7Networks_LH_Default_Temp_2 | -0.015926 | 0.78895 | 0.023261 | 0.84843 |
| 7Networks_LH_Default_Temp_3 | -0.04147 | 0.49615 | 0.023729 | 0.83154 |
| 7Networks_LH_Default_Temp_4 | 0.012872 | 0.82815 | 0.013016 | 0.93042 |
| 7Networks_LH_Default_Temp_5 | -0.12892 | 0.030379 | 0.19183 | 0.18859 |
| 7Networks_LH_Default_Temp_6 | 0.058995 | 0.3182 | 0.028938 | 0.81058 |
| 7Networks_LH_Default_Temp_7 | -0.034855 | 0.57043 | 0.038947 | 0.74722 |
| 7Networks_LH_Default_Temp_8 | -0.055349 | 0.34985 | 0.1208 | 0.35757 |
| 7Networks_LH_Default_Temp_9 | -0.11427 | 0.047385 | 0.4982 | 5.21e-05 |
| 7Networks_LH_Default_PFC_1 | 0.015253 | 0.80163 | 0.023044 | 0.83034 |
| 7Networks_LH_Default_PFC_2 | 0.14183 | 0.019705 | -0.07732 | 0.54361 |
| 7Networks_LH_Default_PFC_3 | -0.029937 | 0.60228 | -0.077713 | 0.63792 |
| 7Networks_LH_Default_PFC_4 | 0.018597 | 0.76949 | -0.063143 | 0.48475 |
| 7Networks_LH_Default_PFC_5 | 0.0052145 | 0.93273 | -0.072579 | 0.54359 |
| 7Networks_LH_Default_PFC_6 | -0.038198 | 0.52674 | 0.035423 | 0.75324 |
| 7Networks_LH_Default_PFC_7 | 0.008005 | 0.9011 | -0.081125 | 0.38503 |
| 7Networks_LH_Default_PFC_8 | -0.04726 | 0.45704 | 0.01941 | 0.85888 |
| 7Networks_LH_Default_PFC_9 | 0.0006621 | 0.99141 | -0.084682 | 0.36825 |
| 7Networks_LH_Default_PFC_10 | 0.034605 | 0.56842 | -0.033061 | 0.7549 |
| 7Networks_LH_Default_PFC_11 | -0.0053709 | 0.92953 | -0.080094 | 0.41108 |
| 7Networks_LH_Default_PFC_12 | 0.023434 | 0.70878 | -0.1089 | 0.25206 |
| 7Networks_LH_Default_PFC_13 | 0.030111 | 0.63253 | -0.068264 | 0.47474 |
| 7Networks_LH_Default_PCC_1 | -0.067327 | 0.29068 | 0.14357 | 0.17746 |
| 7Networks_LH_Default_PCC_2 | 0.020287 | 0.73103 | 0.083937 | 0.48916 |
| 7Networks_LH_Default_PCC_3 | -0.057912 | 0.3506 | 0.055444 | 0.60229 |
| 7Networks_LH_Default_PCC_4 | -0.067292 | 0.28639 | 0.084173 | 0.43057 |
| 7Networks_LH_Default_PHC_1 | 0.023201 | 0.7205 | -0.022727 | 0.81902 |
| 7Networks_RH_Vis_1 | 0.025792 | 0.67275 | 0.11076 | 0.30486 |
| 7Networks_RH_Vis_2 | -0.037932 | 0.54234 | 0.017212 | 0.87986 |
| 7Networks_RH_Vis_3 | -0.069689 | 0.26834 | 0.02386 | 0.82949 |
| 7Networks_RH_Vis_4 | -0.078664 | 0.22617 | 0.065232 | 0.43832 |
| 7Networks_RH_Vis_5 | -0.032746 | 0.58188 | 0.14489 | 0.23971 |
| 7Networks_RH_Vis_6 | 0.060811 | 0.33681 | -0.089479 | 0.35169 |
| 7Networks_RH_Vis_7 | 0.040591 | 0.52061 | -0.10804 | 0.23124 |
| 7Networks_RH_Vis_8 | -0.077806 | 0.22347 | 0.081951 | 0.41943 |
| 7Networks_RH_Vis_9 | -0.12498 | 0.058197 | 0.046815 | 0.58067 |
| 7Networks_RH_Vis_10 | 0.012055 | 0.85353 | 0.027264 | 0.7814 |
| 7Networks_RH_Vis_11 | -0.030673 | 0.59609 | 0.029942 | 0.8172 |
| 7Networks_RH_Vis_12 | -0.063738 | 0.32387 | 0.10698 | 0.24239 |
| 7Networks_RH_Vis_13 | 0.0042855 | 0.94602 | 0.050114 | 0.59646 |
| 7Networks_RH_Vis_14 | -0.10908 | 0.083476 | 0.075251 | 0.45205 |
| 7Networks_RH_Vis_15 | 0.01894 | 0.75207 | -0.003272 | 0.97901 |
| 7Networks_RH_SomMot_1 | 0.0060077 | 0.92545 | -0.037959 | 0.7088 |
| 7Networks_RH_SomMot_2 | 0.0003322 | 0.99578 | -0.017854 | 0.87804 |
| 7Networks_RH_SomMot_3 | 0.073656 | 0.23798 | -0.12889 | 0.25554 |
| 7Networks_RH_SomMot_4 | -0.10396 | 0.091408 | 0.16114 | 0.15342 |
| 7Networks_RH_SomMot_5 | -0.099081 | 0.094896 | 0.30489 | 0.039043 |
| 7Networks_RH_SomMot_6 | 0.03068 | 0.61525 | -0.1115 | 0.36072 |
| 7Networks_RH_SomMot_7 | -0.02715 | 0.6621 | 0.11586 | 0.23764 |
| 7Networks_RH_SomMot_8 | 0.03211 | 0.57914 | 0.014912 | 0.91834 |
| 7Networks_RH_SomMot_9 | 0.14682 | 0.019982 | -0.15484 | 0.19432 |
| 7Networks_RH_SomMot_10 | 0.12265 | 0.040929 | -0.22477 | 0.15868 |
| 7Networks_RH_SomMot_11 | 0.11928 | 0.044933 | -0.13389 | 0.2486 |
| 7Networks_RH_SomMot_12 | 0.033194 | 0.60135 | -0.042358 | 0.66231 |
| 7Networks_RH_SomMot_13 | -0.033354 | 0.57265 | 0.151 | 0.19275 |
| 7Networks_RH_SomMot_14 | -0.013894 | 0.82124 | -0.10084 | 0.33701 |
| 7Networks_RH_SomMot_15 | -0.033134 | 0.59847 | 0.13934 | 0.22787 |
| 7Networks_RH_SomMot_16 | 0.049156 | 0.42372 | -0.16076 | 0.20616 |
| 7Networks_RH_SomMot_17 | -0.013447 | 0.83426 | -0.043259 | 0.6478 |
| 7Networks_RH_SomMot_18 | -0.011777 | 0.84464 | 0.029374 | 0.7772 |
| 7Networks_RH_SomMot_19 | -0.073228 | 0.24941 | 0.078787 | 0.43681 |
| 7Networks_RH_DorsAttn_Post_1 | -0.040541 | 0.4945 | 0.07579 | 0.56922 |
| 7Networks_RH_DorsAttn_Post_2 | 0.010981 | 0.85371 | 0.031156 | 0.81417 |
| 7Networks_RH_DorsAttn_Post_3 | 0.018623 | 0.75483 | -0.0139 | 0.90487 |
| 7Networks_RH_DorsAttn_Post_4 | 0.10217 | 0.085061 | -0.10224 | 0.46516 |
| 7Networks_RH_DorsAttn_Post_5 | 0.07109 | 0.23011 | 0.17888 | 0.23056 |
| 7Networks_RH_DorsAttn_Post_6 | -0.037489 | 0.54228 | 0.0077089 | 0.94115 |
| 7Networks_RH_DorsAttn_Post_7 | 0.094227 | 0.1044 | -0.06988 | 0.67998 |
| 7Networks_RH_DorsAttn_Post_8 | -0.055313 | 0.36767 | 0.10858 | 0.32297 |
| 7Networks_RH_DorsAttn_Post_9 | -0.11172 | 0.06457 | 0.097959 | 0.3821 |
| 7Networks_RH_DorsAttn_Post_10 | 0.0065193 | 0.91602 | 0.066388 | 0.51352 |
| 7Networks_RH_DorsAttn_FEF_1 | 0.057465 | 0.35332 | -0.016993 | 0.88894 |
| 7Networks_RH_DorsAttn_FEF_2 | 0.079956 | 0.17606 | -0.16492 | 0.2391 |
| 7Networks_RH_DorsAttn_PrCv_1 | 0.034624 | 0.57171 | 0.033622 | 0.7768 |
| 7Networks_RH_SalVentAttn_TempOccPar_1 | -0.055551 | 0.3496 | 0.13085 | 0.33651 |
| 7Networks_RH_SalVentAttn_TempOccPar_2 | -0.034108 | 0.55298 | 0.21391 | 0.15619 |
| 7Networks_RH_SalVentAttn_TempOccPar_3 | -0.014745 | 0.80389 | 0.066698 | 0.60599 |
| 7Networks_RH_SalVentAttn_PrC_1 | 0.034203 | 0.5573 | -0.14436 | 0.38077 |
| 7Networks_RH_SalVentAttn_FrOper_1 | -0.10899 | 0.070204 | 0.024027 | 0.85036 |
| 7Networks_RH_SalVentAttn_FrOper_2 | 0.11162 | 0.090163 | -0.039414 | 0.67997 |
| 7Networks_RH_SalVentAttn_FrOper_3 | -0.10961 | 0.063444 | 0.22582 | 0.048594 |
| 7Networks_RH_SalVentAttn_FrOper_4 | -0.038408 | 0.52142 | 0.016602 | 0.89945 |
| 7Networks_RH_SalVentAttn_Med_1 | 0.008653 | 0.88464 | -0.012837 | 0.91935 |
| 7Networks_RH_SalVentAttn_Med_2 | 0.030873 | 0.61957 | 0.028197 | 0.78219 |
| 7Networks_RH_SalVentAttn_Med_3 | 0.1234 | 0.048592 | -0.1865 | 0.1132 |
| 7Networks_RH_Limbic_OFC_1 | 0.055683 | 0.38469 | -0.059048 | 0.5804 |
| 7Networks_RH_Limbic_OFC_2 | 0.097447 | 0.097481 | -0.052069 | 0.6227 |
| 7Networks_RH_Limbic_OFC_3 | 0.025041 | 0.68725 | -0.11927 | 0.27572 |
| 7Networks_RH_Limbic_TempPole_1 | -0.014221 | 0.80784 | -0.13906 | 0.35545 |
| 7Networks_RH_Limbic_TempPole_2 | -0.065724 | 0.28728 | 0.16808 | 0.13421 |
| 7Networks_RH_Limbic_TempPole_3 | 0.020005 | 0.75813 | 0.064125 | 0.49351 |
| 7Networks_RH_Cont_Par_1 | -0.0085431 | 0.88421 | 0.040877 | 0.80632 |
| 7Networks_RH_Cont_Par_2 | 0.056387 | 0.33171 | -0.064617 | 0.65577 |
| 7Networks_RH_Cont_Par_3 | 0.050984 | 0.37838 | -0.025144 | 0.88575 |
| 7Networks_RH_Cont_Temp_1 | -0.019349 | 0.74772 | 0.30613 | 0.033051 |
| 7Networks_RH_Cont_PFCv_1 | 0.10628 | 0.072742 | -0.12425 | 0.3811 |
| 7Networks_RH_Cont_PFCl_1 | 0.031837 | 0.5907 | 0.038827 | 0.75487 |
| 7Networks_RH_Cont_PFCl_2 | 0.051712 | 0.39068 | -0.14617 | 0.21597 |
| 7Networks_RH_Cont_PFCl_3 | -0.015155 | 0.79744 | 0.01252 | 0.91023 |
| 7Networks_RH_Cont_PFCl_4 | 0.10506 | 0.087014 | -0.11848 | 0.24684 |
| 7Networks_RH_Cont_PFCl_5 | 0.001159 | 0.98441 | -0.10959 | 0.32553 |
| 7Networks_RH_Cont_PFCl_6 | -0.042857 | 0.46401 | -0.052324 | 0.64359 |
| 7Networks_RH_Cont_PFCl_7 | 0.061375 | 0.32877 | -0.000357 | 0.99748 |
| 7Networks_RH_Cont_pCun_1 | -0.026653 | 0.65047 | 0.11823 | 0.35632 |
| 7Networks_RH_Cont_PFCmp_1 | -0.022101 | 0.72204 | -0.052293 | 0.57996 |
| 7Networks_RH_Cont_PFCmp_2 | 0.054398 | 0.38748 | -0.079633 | 0.44224 |
| 7Networks_RH_Cont_PFCmp_3 | 0.13632 | 0.022917 | -0.35277 | 0.0055624 |
| 7Networks_RH_Cont_PFCmp_4 | 0.015009 | 0.81673 | -0.08206 | 0.41685 |
| 7Networks_RH_Default_Par_1 | 0.096409 | 0.11484 | -0.1736 | 0.20686 |
| 7Networks_RH_Default_Par_2 | 0.051848 | 0.37915 | 0.035998 | 0.80278 |
| 7Networks_RH_Default_Par_3 | 0.07584 | 0.19682 | -0.10138 | 0.48369 |
| 7Networks_RH_Default_Temp_1 | -0.072035 | 0.23246 | -0.13063 | 0.2536 |
| 7Networks_RH_Default_Temp_2 | -0.06481 | 0.28543 | 0.25578 | 0.027171 |
| 7Networks_RH_Default_Temp_3 | -0.010969 | 0.86504 | 0.15502 | 0.13457 |
| 7Networks_RH_Default_Temp_4 | -0.090786 | 0.12372 | 0.14349 | 0.23308 |
| 7Networks_RH_Default_Temp_5 | -0.029523 | 0.62965 | 0.093208 | 0.48505 |
| 7Networks_RH_Default_PFCv_1 | 0.028163 | 0.64781 | 0.0085197 | 0.93844 |
| 7Networks_RH_Default_PFCm_1 | -0.028774 | 0.62448 | 0.093378 | 0.43438 |
| 7Networks_RH_Default_PFCm_2 | 0.11179 | 0.056766 | -0.088394 | 0.49661 |
| 7Networks_RH_Default_PFCm_3 | -0.079264 | 0.21623 | 0.084014 | 0.46581 |
| 7Networks_RH_Default_PFCm_4 | -0.061389 | 0.35221 | -0.040624 | 0.64718 |
| 7Networks_RH_Default_PFCm_5 | 0.10794 | 0.083631 | -0.2236 | 0.016305 |
| 7Networks_RH_Default_PFCm_6 | 0.12873 | 0.030521 | -0.31017 | 0.0075385 |
| 7Networks_RH_Default_PFCm_7 | 0.062026 | 0.27803 | -0.081251 | 0.58736 |
| 7Networks_RH_Default_PCC_1 | -0.10668 | 0.093358 | 0.108 | 0.33288 |
| 7Networks_RH_Default_PCC_2 | -0.074807 | 0.24147 | 0.066631 | 0.52251 |
| 7Networks_RH_Default_PCC_3 | 0.065721 | 0.25947 | 0.035262 | 0.79011 |

**Supplementary Table 8. Genetic and environmental correlation between Extraversion and cortical thickness in HCP.** Genetic correlation between Extraversion and local cortical thickness calculated using solar 8.4.0. Regions are named according to the Schaefer 200 atlas, based on 7-networks. Here we report environmental correlation (ρ_e_) and genetic correlation (ρ_g_) and the associated p-values.

| ROI | ρ_e_ | p | ρ_g_ | p |
| --- | --- | --- | --- | --- |
| 7Networks_LH_Vis_1 | 0.053537 | 0.39554 | -0.080022 | 0.49455 |
| 7Networks_LH_Vis_2 | -0.085001 | 0.17545 | 0.1109 | 0.40806 |
| 7Networks_LH_Vis_3 | 0.039429 | 0.52086 | -0.29094 | 0.12616 |
| 7Networks_LH_Vis_4 | 0.069618 | 0.26821 | -0.090178 | 0.38445 |
| 7Networks_LH_Vis_5 | 0.040657 | 0.52541 | -0.15432 | 0.19358 |
| 7Networks_LH_Vis_6 | 0.060361 | 0.35379 | 0.024472 | 0.8298 |
| 7Networks_LH_Vis_7 | -0.0089871 | 0.89153 | -0.034108 | 0.74849 |
| 7Networks_LH_Vis_8 | -0.050117 | 0.38866 | -0.055774 | 0.70196 |
| 7Networks_LH_Vis_9 | -0.02416 | 0.70959 | -0.16524 | 0.12112 |
| 7Networks_LH_Vis_10 | 0.016238 | 0.80753 | -0.095617 | 0.35591 |
| 7Networks_LH_Vis_11 | -0.038029 | 0.5355 | -0.14332 | 0.26521 |
| 7Networks_LH_Vis_12 | 0.029155 | 0.66211 | -0.18079 | 0.11137 |
| 7Networks_LH_Vis_13 | -0.092174 | 0.18095 | -0.11126 | 0.28247 |
| 7Networks_LH_Vis_14 | -0.079717 | 0.22427 | -0.18043 | 0.11719 |
| 7Networks_LH_SomMot_1 | 0.091518 | 0.16996 | -0.162 | 0.13853 |
| 7Networks_LH_SomMot_2 | -0.0033659 | 0.95625 | -0.077091 | 0.59107 |
| 7Networks_LH_SomMot_3 | 0.0008662 | 0.98972 | -0.10388 | 0.32722 |
| 7Networks_LH_SomMot_4 | 0.062144 | 0.3342 | 0.095516 | 0.42968 |
| 7Networks_LH_SomMot_5 | 0.080971 | 0.17764 | -0.028336 | 0.85044 |
| 7Networks_LH_SomMot_6 | 0.018197 | 0.78616 | 0.0355 | 0.7311 |
| 7Networks_LH_SomMot_7 | 0.0044337 | 0.94375 | -0.026163 | 0.85543 |
| 7Networks_LH_SomMot_8 | 0.050155 | 0.4305 | -0.16214 | 0.26882 |
| 7Networks_LH_SomMot_9 | 0.039121 | 0.5333 | -0.037334 | 0.77424 |
| 7Networks_LH_SomMot_10 | -0.014293 | 0.8321 | -0.071083 | 0.50275 |
| 7Networks_LH_SomMot_11 | -0.0019291 | 0.9743 | 0.10793 | 0.49893 |
| 7Networks_LH_SomMot_12 | 0.024481 | 0.69978 | 0.050893 | 0.68949 |
| 7Networks_LH_SomMot_13 | 0.037204 | 0.56296 | -0.11595 | 0.39473 |
| 7Networks_LH_SomMot_14 | 0.013938 | 0.82307 | 0.0006109 | 0.99609 |
| 7Networks_LH_SomMot_15 | -0.0079652 | 0.90076 | 0.14473 | 0.20061 |
| 7Networks_LH_SomMot_16 | -0.076053 | 0.25921 | 0.032401 | 0.79724 |
| 7Networks_LH_DorsAttn_Post_1 | -0.16395 | 0.010125 | 0.18221 | 0.19733 |
| 7Networks_LH_DorsAttn_Post_2 | -0.025592 | 0.66519 | 0.085932 | 0.5839 |
| 7Networks_LH_DorsAttn_Post_3 | 0.021458 | 0.73394 | -0.11798 | 0.36056 |
| 7Networks_LH_DorsAttn_Post_4 | -0.072268 | 0.25862 | 0.068141 | 0.6048 |
| 7Networks_LH_DorsAttn_Post_5 | 0.033327 | 0.5771 | -0.16289 | 0.30803 |
| 7Networks_LH_DorsAttn_Post_6 | 0.043485 | 0.48169 | -0.26291 | 0.10776 |
| 7Networks_LH_DorsAttn_Post_7 | 0.081021 | 0.22564 | -0.061497 | 0.59763 |
| 7Networks_LH_DorsAttn_Post_8 | 0.068724 | 0.2678 | -0.079172 | 0.56127 |
| 7Networks_LH_DorsAttn_Post_9 | -0.041357 | 0.52659 | 0.04599 | 0.68303 |
| 7Networks_LH_DorsAttn_Post_10 | 0.012499 | 0.85286 | 0.094899 | 0.40641 |
| 7Networks_LH_DorsAttn_FEF_1 | 0.041959 | 0.49151 | -0.045694 | 0.7518 |
| 7Networks_LH_DorsAttn_FEF_2 | -0.026133 | 0.68853 | 0.27256 | 0.032747 |
| 7Networks_LH_DorsAttn_PrCv_1 | 0.025218 | 0.67801 | -0.12603 | 0.48072 |
| 7Networks_LH_SalVentAttn_ParOper_1 | 0.079801 | 0.18621 | -0.39486 | 0.050257 |
| 7Networks_LH_SalVentAttn_ParOper_2 | 0.09847 | 0.10104 | 0.062847 | 0.65047 |
| 7Networks_LH_SalVentAttn_ParOper_3 | -0.027282 | 0.65669 | 0.25781 | 0.11783 |
| 7Networks_LH_SalVentAttn_FrOper_1 | 0.021535 | 0.7316 | 0.081292 | 0.52399 |
| 7Networks_LH_SalVentAttn_FrOper_2 | 0.0040228 | 0.95092 | -0.02999 | 0.80516 |
| 7Networks_LH_SalVentAttn_FrOper_3 | -0.065504 | 0.31622 | 0.18495 | 0.11493 |
| 7Networks_LH_SalVentAttn_FrOper_4 | 0.078521 | 0.21202 | -0.23768 | 0.14091 |
| 7Networks_LH_SalVentAttn_PFCl_1 | 0.031273 | 0.62011 | 0.028991 | 0.79279 |
| 7Networks_LH_SalVentAttn_Med_1 | -0.031516 | 0.62802 | -0.0049621 | 0.97093 |
| 7Networks_LH_SalVentAttn_Med_2 | -0.0509 | 0.42794 | -0.051951 | 0.70107 |
| 7Networks_LH_SalVentAttn_Med_3 | -0.16597 | 0.0075355 | 0.31922 | 0.037709 |
| 7Networks_LH_Limbic_OFC_1 | -0.047383 | 0.43451 | 0.11043 | 0.41854 |
| 7Networks_LH_Limbic_OFC_2 | -0.026395 | 0.67966 | 0.14835 | 0.24876 |
| 7Networks_LH_Limbic_TempPole_1 | 0.0058285 | 0.92838 | -0.0025622 | 0.98482 |
| 7Networks_LH_Limbic_TempPole_2 | -0.0051759 | 0.93266 | 0.03622 | 0.76234 |
| 7Networks_LH_Limbic_TempPole_3 | -0.023271 | 0.69305 | 0.11885 | 0.42108 |
| 7Networks_LH_Limbic_TempPole_4 | 0.11511 | 0.079438 | -0.23433 | 0.040042 |
| 7Networks_LH_Cont_Par_1 | -0.042268 | 0.48702 | 0.095689 | 0.55504 |
| 7Networks_LH_Cont_Par_2 | 0.00833 | 0.88991 | -0.098702 | 0.48822 |
| 7Networks_LH_Cont_Par_3 | 0.0064707 | 0.91106 | -0.17782 | 0.6151 |
| 7Networks_LH_Cont_Temp_1 | -0.11517 | 0.056294 | 0.15167 | 0.3721 |
| 7Networks_LH_Cont_PFCl_1 | -0.10445 | 0.07608 | 0.14082 | 0.31262 |
| 7Networks_LH_Cont_PFCl_2 | 0.0024864 | 0.96805 | 0.12924 | 0.35719 |
| 7Networks_LH_Cont_PFCl_3 | 0.022492 | 0.72658 | -0.035246 | 0.77393 |
| 7Networks_LH_Cont_PFCl_4 | 0.11022 | 0.068968 | 0.0081348 | 0.95538 |
| 7Networks_LH_Cont_PFCl_5 | 0.078468 | 0.2138 | 0.044752 | 0.69926 |
| 7Networks_LH_Cont_PFCl_6 | -0.028699 | 0.63527 | 0.046735 | 0.77889 |
| 7Networks_LH_Cont_pCun_1 | 0.017133 | 0.78424 | -0.028656 | 0.81372 |
| 7Networks_LH_Cont_Cing_1 | 0.062287 | 0.36574 | 0.049674 | 0.64097 |
| 7Networks_LH_Cont_Cing_2 | -0.018677 | 0.77418 | 0.029035 | 0.82281 |
| 7Networks_LH_Default_Temp_1 | -0.019248 | 0.76154 | 0.08072 | 0.5824 |
| 7Networks_LH_Default_Temp_2 | -0.18277 | 0.0025379 | 0.30708 | 0.028699 |
| 7Networks_LH_Default_Temp_3 | -0.037831 | 0.54635 | 0.059492 | 0.64693 |
| 7Networks_LH_Default_Temp_4 | -0.02172 | 0.7212 | 0.11325 | 0.51445 |
| 7Networks_LH_Default_Temp_5 | 0.12704 | 0.044484 | -0.2608 | 0.10646 |
| 7Networks_LH_Default_Temp_6 | -0.094622 | 0.11834 | -0.096021 | 0.49354 |
| 7Networks_LH_Default_Temp_7 | -0.021291 | 0.73531 | -0.15165 | 0.27841 |
| 7Networks_LH_Default_Temp_8 | 0.02711 | 0.6559 | -0.25684 | 0.089084 |
| 7Networks_LH_Default_Temp_9 | -0.01211 | 0.83802 | -0.11566 | 0.41095 |
| 7Networks_LH_Default_PFC_1 | 0.008559 | 0.89191 | 0.063287 | 0.61523 |
| 7Networks_LH_Default_PFC_2 | 0.0097872 | 0.87605 | -0.091627 | 0.53662 |
| 7Networks_LH_Default_PFC_3 | -0.054574 | 0.34975 | 0.54331 | 0.0062386 |
| 7Networks_LH_Default_PFC_4 | 0.0734 | 0.26474 | 0.029364 | 0.7767 |
| 7Networks_LH_Default_PFC_5 | 0.036195 | 0.57149 | 0.10258 | 0.4587 |
| 7Networks_LH_Default_PFC_6 | 0.046967 | 0.45348 | -0.081911 | 0.52946 |
| 7Networks_LH_Default_PFC_7 | 0.018851 | 0.77837 | 0.088869 | 0.40692 |
| 7Networks_LH_Default_PFC_8 | 0.041454 | 0.52853 | -0.23808 | 0.060608 |
| 7Networks_LH_Default_PFC_9 | 0.077158 | 0.23269 | 0.16521 | 0.12984 |
| 7Networks_LH_Default_PFC_10 | -0.0014469 | 0.98126 | 0.2267 | 0.064771 |
| 7Networks_LH_Default_PFC_11 | 0.013029 | 0.83558 | 0.19112 | 0.087605 |
| 7Networks_LH_Default_PFC_12 | 0.073851 | 0.25637 | 0.039332 | 0.71993 |
| 7Networks_LH_Default_PFC_13 | 0.011871 | 0.85637 | 0.2071 | 0.060798 |
| 7Networks_LH_Default_PCC_1 | 0.041692 | 0.52181 | -0.027633 | 0.82436 |
| 7Networks_LH_Default_PCC_2 | 0.011513 | 0.84931 | -0.034073 | 0.80824 |
| 7Networks_LH_Default_PCC_3 | -0.015493 | 0.81005 | 0.0045453 | 0.97047 |
| 7Networks_LH_Default_PCC_4 | 0.022176 | 0.73478 | -0.07873 | 0.5274 |
| 7Networks_LH_Default_PHC_1 | -0.012888 | 0.84913 | 0.072734 | 0.53178 |
| 7Networks_RH_Vis_1 | -0.10692 | 0.088373 | 0.069403 | 0.58062 |
| 7Networks_RH_Vis_2 | 0.079509 | 0.21906 | -0.19484 | 0.13632 |
| 7Networks_RH_Vis_3 | 0.046323 | 0.47771 | -0.022524 | 0.86002 |
| 7Networks_RH_Vis_4 | 0.20965 | 0.0018512 | -0.22295 | 0.021199 |
| 7Networks_RH_Vis_5 | -0.10183 | 0.097222 | 0.11306 | 0.42317 |
| 7Networks_RH_Vis_6 | 0.14308 | 0.028281 | -0.18751 | 0.089969 |
| 7Networks_RH_Vis_7 | 0.035772 | 0.58779 | 0.019244 | 0.85419 |
| 7Networks_RH_Vis_8 | 0.01956 | 0.76714 | -0.14968 | 0.20488 |
| 7Networks_RH_Vis_9 | 0.11032 | 0.10562 | -0.13142 | 0.18503 |
| 7Networks_RH_Vis_10 | 0.10643 | 0.11375 | -0.27396 | 0.016507 |
| 7Networks_RH_Vis_11 | -0.054489 | 0.35761 | -0.15089 | 0.31436 |
| 7Networks_RH_Vis_12 | -0.018922 | 0.77813 | -0.16057 | 0.13046 |
| 7Networks_RH_Vis_13 | 0.10284 | 0.1147 | -0.19221 | 0.084952 |
| 7Networks_RH_Vis_14 | 0.055131 | 0.39844 | -0.065248 | 0.5747 |
| 7Networks_RH_Vis_15 | 0.0061621 | 0.9205 | -0.2335 | 0.10633 |
| 7Networks_RH_SomMot_1 | 0.032782 | 0.61995 | -0.0341 | 0.77207 |
| 7Networks_RH_SomMot_2 | -0.03881 | 0.55165 | 0.065865 | 0.62779 |
| 7Networks_RH_SomMot_3 | -0.028605 | 0.65841 | -0.096233 | 0.46683 |
| 7Networks_RH_SomMot_4 | 0.13351 | 0.03729 | -0.32774 | 0.010006 |
| 7Networks_RH_SomMot_5 | 0.02142 | 0.72438 | -0.10228 | 0.5575 |
| 7Networks_RH_SomMot_6 | -0.059529 | 0.34211 | 0.39681 | 0.0043669 |
| 7Networks_RH_SomMot_7 | -0.027023 | 0.67459 | 0.089866 | 0.42844 |
| 7Networks_RH_SomMot_8 | -0.012817 | 0.82893 | -0.15698 | 0.3524 |
| 7Networks_RH_SomMot_9 | -0.061856 | 0.34738 | 0.099585 | 0.46975 |
| 7Networks_RH_SomMot_10 | -0.083616 | 0.17576 | 0.33604 | 0.07144 |
| 7Networks_RH_SomMot_11 | -0.054939 | 0.36693 | -0.060138 | 0.65642 |
| 7Networks_RH_SomMot_12 | 0.029496 | 0.65251 | -0.095682 | 0.39314 |
| 7Networks_RH_SomMot_13 | 0.039699 | 0.51591 | -0.10734 | 0.42019 |
| 7Networks_RH_SomMot_14 | -0.060215 | 0.34624 | 0.20887 | 0.082407 |
| 7Networks_RH_SomMot_15 | 0.074078 | 0.25471 | -0.21551 | 0.1102 |
| 7Networks_RH_SomMot_16 | 0.044258 | 0.48576 | -0.017018 | 0.9083 |
| 7Networks_RH_SomMot_17 | 0.016732 | 0.80182 | 0.0034331 | 0.9743 |
| 7Networks_RH_SomMot_18 | -0.0002032 | 0.99724 | 0.0729 | 0.54356 |
| 7Networks_RH_SomMot_19 | 0.037853 | 0.56816 | -0.044358 | 0.7071 |
| 7Networks_RH_DorsAttn_Post_1 | 0.12956 | 0.033488 | -0.39271 | 0.0090691 |
| 7Networks_RH_DorsAttn_Post_2 | -0.042592 | 0.4863 | 0.047714 | 0.75593 |
| 7Networks_RH_DorsAttn_Post_3 | -0.055149 | 0.36853 | 0.1499 | 0.26152 |
| 7Networks_RH_DorsAttn_Post_4 | -0.066956 | 0.27209 | 0.035153 | 0.82871 |
| 7Networks_RH_DorsAttn_Post_5 | -0.034534 | 0.57084 | 0.002895 | 0.9866 |
| 7Networks_RH_DorsAttn_Post_6 | -0.048955 | 0.44155 | 0.016534 | 0.89107 |
| 7Networks_RH_DorsAttn_Post_7 | -0.052304 | 0.37659 | -0.13033 | 0.50992 |
| 7Networks_RH_DorsAttn_Post_8 | -0.021661 | 0.73243 | -0.04801 | 0.70717 |
| 7Networks_RH_DorsAttn_Post_9 | -0.012034 | 0.84687 | -0.0079447 | 0.95142 |
| 7Networks_RH_DorsAttn_Post_10 | 0.063493 | 0.32141 | -0.054187 | 0.64792 |
| 7Networks_RH_DorsAttn_FEF_1 | 0.036238 | 0.57044 | -0.082884 | 0.55697 |
| 7Networks_RH_DorsAttn_FEF_2 | -0.10139 | 0.094386 | 0.23664 | 0.13632 |
| 7Networks_RH_DorsAttn_PrCv_1 | 0.043708 | 0.48577 | 0.015835 | 0.90703 |
| 7Networks_RH_SalVentAttn_TempOccPar_1 | 0.025134 | 0.68427 | -0.27981 | 0.07122 |
| 7Networks_RH_SalVentAttn_TempOccPar_2 | 0.025227 | 0.66675 | -0.046896 | 0.79158 |
| 7Networks_RH_SalVentAttn_TempOccPar_3 | -0.0060195 | 0.92069 | -0.046135 | 0.75756 |
| 7Networks_RH_SalVentAttn_PrC_1 | -0.039251 | 0.51251 | 0.12158 | 0.5175 |
| 7Networks_RH_SalVentAttn_FrOper_1 | 0.066307 | 0.28731 | -0.16304 | 0.26723 |
| 7Networks_RH_SalVentAttn_FrOper_2 | -0.0089948 | 0.8961 | -0.048914 | 0.66139 |
| 7Networks_RH_SalVentAttn_FrOper_3 | -0.02695 | 0.66061 | -0.093119 | 0.47909 |
| 7Networks_RH_SalVentAttn_FrOper_4 | -0.023284 | 0.70533 | 0.073645 | 0.62407 |
| 7Networks_RH_SalVentAttn_Med_1 | 0.0077895 | 0.89902 | -0.087767 | 0.5504 |
| 7Networks_RH_SalVentAttn_Med_2 | -0.02831 | 0.65829 | -0.1114 | 0.3448 |
| 7Networks_RH_SalVentAttn_Med_3 | -0.055515 | 0.39558 | 0.18657 | 0.171 |
| 7Networks_RH_Limbic_OFC_1 | -0.020512 | 0.75904 | -0.0068245 | 0.9561 |
| 7Networks_RH_Limbic_OFC_2 | -0.027698 | 0.65098 | -0.0096158 | 0.93808 |
| 7Networks_RH_Limbic_OFC_3 | -0.00191 | 0.97631 | -0.0046917 | 0.96927 |
| 7Networks_RH_Limbic_TempPole_1 | 0.0028129 | 0.96242 | 0.0094998 | 0.95699 |
| 7Networks_RH_Limbic_TempPole_2 | 0.04069 | 0.52812 | -0.21731 | 0.094393 |
| 7Networks_RH_Limbic_TempPole_3 | -0.017347 | 0.79823 | 0.020666 | 0.84912 |
| 7Networks_RH_Cont_Par_1 | 0.042869 | 0.47291 | -0.030203 | 0.87495 |
| 7Networks_RH_Cont_Par_2 | -0.049235 | 0.40761 | 0.047455 | 0.77758 |
| 7Networks_RH_Cont_Par_3 | -0.0673 | 0.25583 | -0.082241 | 0.68871 |
| 7Networks_RH_Cont_Temp_1 | -0.0271 | 0.66604 | 0.06713 | 0.68218 |
| 7Networks_RH_Cont_PFCv_1 | -0.033149 | 0.58753 | -0.11875 | 0.47414 |
| 7Networks_RH_Cont_PFCl_1 | -0.049598 | 0.41437 | -0.063746 | 0.65747 |
| 7Networks_RH_Cont_PFCl_2 | 0.0083144 | 0.89339 | -0.081534 | 0.55489 |
| 7Networks_RH_Cont_PFCl_3 | -0.011735 | 0.84692 | 0.060028 | 0.63876 |
| 7Networks_RH_Cont_PFCl_4 | -0.041926 | 0.51093 | 0.089136 | 0.44296 |
| 7Networks_RH_Cont_PFCl_5 | 0.010977 | 0.85801 | 0.078803 | 0.54055 |
| 7Networks_RH_Cont_PFCl_6 | 0.029267 | 0.62772 | 0.26061 | 0.044661 |
| 7Networks_RH_Cont_PFCl_7 | -0.054037 | 0.40326 | 0.1808 | 0.17861 |
| 7Networks_RH_Cont_pCun_1 | 0.026904 | 0.65622 | -0.093957 | 0.51925 |
| 7Networks_RH_Cont_PFCmp_1 | -0.082324 | 0.20068 | 0.18823 | 0.080778 |
| 7Networks_RH_Cont_PFCmp_2 | -0.12242 | 0.059834 | 0.20125 | 0.093539 |
| 7Networks_RH_Cont_PFCmp_3 | -0.0023353 | 0.97013 | 0.075503 | 0.60523 |
| 7Networks_RH_Cont_PFCmp_4 | 0.037722 | 0.57667 | 0.15783 | 0.17972 |
| 7Networks_RH_Default_Par_1 | -0.092129 | 0.14154 | 0.20127 | 0.20897 |
| 7Networks_RH_Default_Par_2 | -0.13136 | 0.029462 | 0.19083 | 0.25272 |
| 7Networks_RH_Default_Par_3 | -0.096041 | 0.10903 | 0.17631 | 0.29545 |
| 7Networks_RH_Default_Temp_1 | 0.10527 | 0.090832 | -0.0009406 | 0.99403 |
| 7Networks_RH_Default_Temp_2 | -0.032442 | 0.60537 | 0.046196 | 0.73227 |
| 7Networks_RH_Default_Temp_3 | -0.04314 | 0.52315 | -0.0025058 | 0.98349 |
| 7Networks_RH_Default_Temp_4 | 0.046821 | 0.44162 | 0.027101 | 0.84237 |
| 7Networks_RH_Default_Temp_5 | -0.01359 | 0.829 | -0.094507 | 0.54411 |
| 7Networks_RH_Default_PFCv_1 | -0.020133 | 0.75225 | 0.14711 | 0.25633 |
| 7Networks_RH_Default_PFCm_1 | 0.12364 | 0.040129 | -0.2789 | 0.043711 |
| 7Networks_RH_Default_PFCm_2 | 0.0011902 | 0.98433 | -0.047617 | 0.75028 |
| 7Networks_RH_Default_PFCm_3 | -0.052223 | 0.43133 | 0.11435 | 0.40067 |
| 7Networks_RH_Default_PFCm_4 | 0.028907 | 0.6746 | 0.1454 | 0.15898 |
| 7Networks_RH_Default_PFCm_5 | 0.040152 | 0.53835 | 0.20964 | 0.050636 |
| 7Networks_RH_Default_PFCm_6 | -0.060876 | 0.32567 | 0.23515 | 0.074308 |
| 7Networks_RH_Default_PFCm_7 | -0.0021966 | 0.97005 | 0.062016 | 0.71978 |
| 7Networks_RH_Default_PCC_1 | 0.065014 | 0.32566 | -0.12205 | 0.34071 |
| 7Networks_RH_Default_PCC_2 | 0.032446 | 0.62287 | -0.14395 | 0.23764 |
| 7Networks_RH_Default_PCC_3 | -0.046719 | 0.43497 | -0.10252 | 0.50449 |

**Supplementary Table 9. Genetic and environmental correlation between Neuroticism and cortical thickness in HCP.** Genetic correlation between Neuroticism and local cortical thickness calculated using solar 8.4.0. Regions are named according to the Schaefer 200 atlas, based on 7-networks. Here we report environmental correlation (ρ_e_) and genetic correlation (ρ_g_) and the associated p-values.

| ROI | ρ_e_ | p | ρ_g_ | p |
| --- | --- | --- | --- | --- |
| 7Networks_LH_Vis_1 | -0.047877 | 0.45749 | 0.25034 | 0.0042261 |
| 7Networks_LH_Vis_2 | -0.0027296 | 0.96605 | 0.010559 | 0.91487 |
| 7Networks_LH_Vis_3 | 0.0020641 | 0.97383 | -0.13515 | 0.34983 |
| 7Networks_LH_Vis_4 | 0.065051 | 0.30471 | 0.026561 | 0.73032 |
| 7Networks_LH_Vis_5 | 0.031692 | 0.62767 | -0.055558 | 0.5298 |
| 7Networks_LH_Vis_6 | -0.016376 | 0.80384 | 0.068957 | 0.40936 |
| 7Networks_LH_Vis_7 | -0.0011101 | 0.98665 | 0.074864 | 0.34494 |
| 7Networks_LH_Vis_8 | -0.14611 | 0.014575 | 0.20597 | 0.057727 |
| 7Networks_LH_Vis_9 | 0.11043 | 0.089814 | 0.0078776 | 0.92191 |
| 7Networks_LH_Vis_10 | -0.084906 | 0.20532 | 0.085363 | 0.26812 |
| 7Networks_LH_Vis_11 | 0.19401 | 0.0019177 | -0.17036 | 0.07541 |
| 7Networks_LH_Vis_12 | -0.027136 | 0.68839 | 0.035967 | 0.66643 |
| 7Networks_LH_Vis_13 | -0.082967 | 0.23039 | 0.024316 | 0.75156 |
| 7Networks_LH_Vis_14 | 0.085281 | 0.20056 | -0.098157 | 0.24969 |
| 7Networks_LH_SomMot_1 | -0.040441 | 0.54663 | 0.14549 | 0.074439 |
| 7Networks_LH_SomMot_2 | -0.031094 | 0.6214 | 0.1756 | 0.099401 |
| 7Networks_LH_SomMot_3 | 0.045761 | 0.49958 | 0.061823 | 0.43304 |
| 7Networks_LH_SomMot_4 | 0.094192 | 0.14969 | 0.040456 | 0.65443 |
| 7Networks_LH_SomMot_5 | -0.036404 | 0.55775 | -0.081721 | 0.46516 |
| 7Networks_LH_SomMot_6 | 0.1341 | 0.046001 | -0.045297 | 0.55884 |
| 7Networks_LH_SomMot_7 | -0.038099 | 0.56128 | 0.038746 | 0.71698 |
| 7Networks_LH_SomMot_8 | 0.099988 | 0.12581 | -0.25277 | 0.01901 |
| 7Networks_LH_SomMot_9 | -0.018037 | 0.78009 | -0.10423 | 0.2824 |
| 7Networks_LH_SomMot_10 | 0.064797 | 0.33952 | 0.040622 | 0.6086 |
| 7Networks_LH_SomMot_11 | -0.0013146 | 0.98293 | -0.19634 | 0.098135 |
| 7Networks_LH_SomMot_12 | -0.0023356 | 0.96948 | -0.013002 | 0.89045 |
| 7Networks_LH_SomMot_13 | -0.13066 | 0.047272 | 0.061289 | 0.5401 |
| 7Networks_LH_SomMot_14 | 0.012453 | 0.84395 | -0.050036 | 0.58395 |
| 7Networks_LH_SomMot_15 | 0.13834 | 0.037146 | -0.087852 | 0.28975 |
| 7Networks_LH_SomMot_16 | 0.12312 | 0.071644 | -0.043525 | 0.65425 |
| 7Networks_LH_DorsAttn_Post_1 | 0.054777 | 0.41234 | -0.0047965 | 0.96245 |
| 7Networks_LH_DorsAttn_Post_2 | -0.074136 | 0.22368 | -0.026498 | 0.82176 |
| 7Networks_LH_DorsAttn_Post_3 | 0.0059476 | 0.92658 | -0.12272 | 0.20424 |
| 7Networks_LH_DorsAttn_Post_4 | 0.040624 | 0.53783 | -0.043703 | 0.65485 |
| 7Networks_LH_DorsAttn_Post_5 | 0.0055079 | 0.92948 | -0.12125 | 0.30778 |
| 7Networks_LH_DorsAttn_Post_6 | 0.047818 | 0.45868 | -0.31287 | 0.0081973 |
| 7Networks_LH_DorsAttn_Post_7 | 0.048414 | 0.48084 | -0.15208 | 0.071998 |
| 7Networks_LH_DorsAttn_Post_8 | -0.0083791 | 0.89699 | -0.14415 | 0.15702 |
| 7Networks_LH_DorsAttn_Post_9 | 0.088791 | 0.18619 | -0.10055 | 0.2293 |
| 7Networks_LH_DorsAttn_Post_10 | 0.077377 | 0.25989 | -0.12791 | 0.12901 |
| 7Networks_LH_DorsAttn_FEF_1 | 0.02409 | 0.69992 | -0.11174 | 0.29934 |
| 7Networks_LH_DorsAttn_FEF_2 | -0.0054613 | 0.93412 | -0.11127 | 0.24493 |
| 7Networks_LH_DorsAttn_PrCv_1 | -0.14019 | 0.025654 | 0.27029 | 0.034949 |
| 7Networks_LH_SalVentAttn_ParOper_1 | -0.011825 | 0.84995 | -0.1362 | 0.37587 |
| 7Networks_LH_SalVentAttn_ParOper_2 | 0.1058 | 0.08768 | -0.15232 | 0.1434 |
| 7Networks_LH_SalVentAttn_ParOper_3 | 0.046311 | 0.4699 | -0.1865 | 0.13163 |
| 7Networks_LH_SalVentAttn_FrOper_1 | 0.031429 | 0.62575 | -0.014733 | 0.87668 |
| 7Networks_LH_SalVentAttn_FrOper_2 | -0.077199 | 0.24107 | 0.15156 | 0.092961 |
| 7Networks_LH_SalVentAttn_FrOper_3 | -0.0031189 | 0.96208 | -0.0062703 | 0.94248 |
| 7Networks_LH_SalVentAttn_FrOper_4 | -0.12037 | 0.06786 | 0.15277 | 0.18585 |
| 7Networks_LH_SalVentAttn_PFCl_1 | -0.013474 | 0.83232 | -0.19769 | 0.016261 |
| 7Networks_LH_SalVentAttn_Med_1 | 0.058263 | 0.37685 | -0.16551 | 0.099074 |
| 7Networks_LH_SalVentAttn_Med_2 | 0.023709 | 0.71767 | -0.10352 | 0.30122 |
| 7Networks_LH_SalVentAttn_Med_3 | -0.016261 | 0.7995 | -0.019578 | 0.86445 |
| 7Networks_LH_Limbic_OFC_1 | -0.064079 | 0.3039 | -0.010361 | 0.91867 |
| 7Networks_LH_Limbic_OFC_2 | -0.046152 | 0.48162 | 0.073162 | 0.44974 |
| 7Networks_LH_Limbic_TempPole_1 | -0.12799 | 0.053521 | 0.1865 | 0.060594 |
| 7Networks_LH_Limbic_TempPole_2 | -0.11102 | 0.073703 | 0.11613 | 0.19772 |
| 7Networks_LH_Limbic_TempPole_3 | -0.093617 | 0.12441 | 0.22367 | 0.040437 |
| 7Networks_LH_Limbic_TempPole_4 | 0.048096 | 0.47474 | 0.10383 | 0.22468 |
| 7Networks_LH_Cont_Par_1 | 0.012376 | 0.84411 | -0.24586 | 0.041558 |
| 7Networks_LH_Cont_Par_2 | 0.055685 | 0.37754 | -0.16076 | 0.1255 |
| 7Networks_LH_Cont_Par_3 | 0.023072 | 0.7035 | -0.61527 | 0.023377 |
| 7Networks_LH_Cont_Temp_1 | -0.051739 | 0.40668 | 0.015634 | 0.90259 |
| 7Networks_LH_Cont_PFCl_1 | -0.028233 | 0.63992 | -0.12102 | 0.25479 |
| 7Networks_LH_Cont_PFCl_2 | -0.056376 | 0.37569 | 0.0026803 | 0.97926 |
| 7Networks_LH_Cont_PFCl_3 | -0.025933 | 0.6915 | -0.17792 | 0.047717 |
| 7Networks_LH_Cont_PFCl_4 | -0.12514 | 0.046221 | -0.14873 | 0.17119 |
| 7Networks_LH_Cont_PFCl_5 | -0.060778 | 0.34531 | -0.020322 | 0.81417 |
| 7Networks_LH_Cont_PFCl_6 | -0.10145 | 0.10372 | 0.02227 | 0.85701 |
| 7Networks_LH_Cont_pCun_1 | -0.13945 | 0.028334 | 0.064878 | 0.47417 |
| 7Networks_LH_Cont_Cing_1 | -0.16194 | 0.017895 | 0.13782 | 0.078656 |
| 7Networks_LH_Cont_Cing_2 | -0.031663 | 0.63316 | 0.093499 | 0.32624 |
| 7Networks_LH_Default_Temp_1 | -0.1366 | 0.037612 | 0.28867 | 0.0067798 |
| 7Networks_LH_Default_Temp_2 | -0.18056 | 0.0037357 | 0.032467 | 0.75837 |
| 7Networks_LH_Default_Temp_3 | -0.025449 | 0.69124 | 0.087542 | 0.36563 |
| 7Networks_LH_Default_Temp_4 | -0.094748 | 0.12986 | 0.2418 | 0.068916 |
| 7Networks_LH_Default_Temp_5 | -0.10286 | 0.10565 | 0.2069 | 0.10886 |
| 7Networks_LH_Default_Temp_6 | -0.082272 | 0.18578 | 0.2374 | 0.024113 |
| 7Networks_LH_Default_Temp_7 | 0.036942 | 0.5685 | -0.19257 | 0.063367 |
| 7Networks_LH_Default_Temp_8 | -0.042844 | 0.49552 | -0.0049762 | 0.96433 |
| 7Networks_LH_Default_Temp_9 | -0.026336 | 0.66691 | -0.093316 | 0.37622 |
| 7Networks_LH_Default_PFC_1 | -0.0037976 | 0.95269 | 0.044949 | 0.62807 |
| 7Networks_LH_Default_PFC_2 | 0.066145 | 0.3062 | -0.042077 | 0.70106 |
| 7Networks_LH_Default_PFC_3 | -0.087375 | 0.15028 | 0.18103 | 0.19311 |
| 7Networks_LH_Default_PFC_4 | -0.027099 | 0.68338 | -0.088624 | 0.25236 |
| 7Networks_LH_Default_PFC_5 | -0.1218 | 0.06425 | 0.046441 | 0.64919 |
| 7Networks_LH_Default_PFC_6 | -0.065028 | 0.30734 | -0.037655 | 0.69885 |
| 7Networks_LH_Default_PFC_7 | -0.01271 | 0.85047 | -0.12836 | 0.10752 |
| 7Networks_LH_Default_PFC_8 | -0.065448 | 0.33233 | 0.037162 | 0.69332 |
| 7Networks_LH_Default_PFC_9 | -0.050075 | 0.44229 | -0.12706 | 0.11676 |
| 7Networks_LH_Default_PFC_10 | 0.043267 | 0.49506 | -0.14947 | 0.096427 |
| 7Networks_LH_Default_PFC_11 | 0.01802 | 0.77713 | -0.19591 | 0.019358 |
| 7Networks_LH_Default_PFC_12 | -0.049657 | 0.45166 | -0.078663 | 0.33485 |
| 7Networks_LH_Default_PFC_13 | -0.014897 | 0.82266 | -0.033393 | 0.68652 |
| 7Networks_LH_Default_PCC_1 | -0.20284 | 0.0021478 | 0.31793 | 0.0004296 |
| 7Networks_LH_Default_PCC_2 | 0.031194 | 0.61785 | 0.054649 | 0.60259 |
| 7Networks_LH_Default_PCC_3 | 0.060689 | 0.35253 | -0.13911 | 0.12258 |
| 7Networks_LH_Default_PCC_4 | -0.024948 | 0.70905 | -0.047753 | 0.6054 |
| 7Networks_LH_Default_PHC_1 | -0.08906 | 0.19414 | 0.11631 | 0.17103 |
| 7Networks_RH_Vis_1 | 0.011955 | 0.85296 | 0.039556 | 0.67133 |
| 7Networks_RH_Vis_2 | -0.044233 | 0.50293 | 0.053047 | 0.59047 |
| 7Networks_RH_Vis_3 | -0.034403 | 0.60608 | 0.17045 | 0.075226 |
| 7Networks_RH_Vis_4 | 0.014586 | 0.83018 | 0.058569 | 0.4162 |
| 7Networks_RH_Vis_5 | 2.23e-05 | 1 | 0.16618 | 0.11398 |
| 7Networks_RH_Vis_6 | -0.15573 | 0.018292 | 0.21051 | 0.014021 |
| 7Networks_RH_Vis_7 | 0.011149 | 0.8669 | 0.031504 | 0.68596 |
| 7Networks_RH_Vis_8 | -0.014096 | 0.83416 | 0.088537 | 0.31086 |
| 7Networks_RH_Vis_9 | -0.043623 | 0.52785 | 0.0017798 | 0.98085 |
| 7Networks_RH_Vis_10 | 0.015104 | 0.82725 | 0.017292 | 0.8381 |
| 7Networks_RH_Vis_11 | 0.013021 | 0.83155 | 0.046146 | 0.67972 |
| 7Networks_RH_Vis_12 | 0.059262 | 0.38287 | 0.003335 | 0.96442 |
| 7Networks_RH_Vis_13 | 0.041056 | 0.53805 | 0.090736 | 0.26667 |
| 7Networks_RH_Vis_14 | 0.034435 | 0.60881 | -0.05475 | 0.52642 |
| 7Networks_RH_Vis_15 | 0.10597 | 0.098354 | -0.092332 | 0.38799 |
| 7Networks_RH_SomMot_1 | 0.087682 | 0.1907 | 0.062613 | 0.4715 |
| 7Networks_RH_SomMot_2 | -0.012096 | 0.85655 | 0.06734 | 0.50151 |
| 7Networks_RH_SomMot_3 | 0.0020774 | 0.97408 | 0.021084 | 0.8304 |
| 7Networks_RH_SomMot_4 | -0.057146 | 0.37929 | 0.13923 | 0.15527 |
| 7Networks_RH_SomMot_5 | 0.008822 | 0.88914 | 0.004562 | 0.97189 |
| 7Networks_RH_SomMot_6 | 0.075046 | 0.2463 | -0.11337 | 0.28198 |
| 7Networks_RH_SomMot_7 | 0.090575 | 0.16481 | -0.041182 | 0.62654 |
| 7Networks_RH_SomMot_8 | 0.069932 | 0.25704 | -0.052423 | 0.67574 |
| 7Networks_RH_SomMot_9 | -0.037802 | 0.57812 | -0.062285 | 0.54451 |
| 7Networks_RH_SomMot_10 | 0.01087 | 0.86578 | 0.031528 | 0.82012 |
| 7Networks_RH_SomMot_11 | 0.07053 | 0.26825 | -0.093008 | 0.35519 |
| 7Networks_RH_SomMot_12 | 0.12959 | 0.049368 | 0.0179 | 0.82962 |
| 7Networks_RH_SomMot_13 | 0.11337 | 0.071887 | -0.20319 | 0.038311 |
| 7Networks_RH_SomMot_14 | -0.0188 | 0.77175 | -0.022283 | 0.80584 |
| 7Networks_RH_SomMot_15 | 0.01978 | 0.77068 | -0.012566 | 0.89989 |
| 7Networks_RH_SomMot_16 | 0.022899 | 0.72677 | -0.05458 | 0.61832 |
| 7Networks_RH_SomMot_17 | -0.0043214 | 0.94892 | 0.051149 | 0.52805 |
| 7Networks_RH_SomMot_18 | 0.069537 | 0.27114 | 0.039778 | 0.65626 |
| 7Networks_RH_SomMot_19 | 0.14314 | 0.032471 | -0.0065742 | 0.94095 |
| 7Networks_RH_DorsAttn_Post_1 | 0.041513 | 0.51182 | 0.16284 | 0.15244 |
| 7Networks_RH_DorsAttn_Post_2 | 0.0081352 | 0.89745 | 0.17148 | 0.13439 |
| 7Networks_RH_DorsAttn_Post_3 | 0.03494 | 0.57841 | -0.10873 | 0.27805 |
| 7Networks_RH_DorsAttn_Post_4 | -0.0082814 | 0.89591 | -0.12842 | 0.28487 |
| 7Networks_RH_DorsAttn_Post_5 | 0.012089 | 0.84967 | -0.044443 | 0.73021 |
| 7Networks_RH_DorsAttn_Post_6 | 0.030115 | 0.64281 | -0.11829 | 0.18505 |
| 7Networks_RH_DorsAttn_Post_7 | 0.0056954 | 0.92648 | -0.13212 | 0.36486 |
| 7Networks_RH_DorsAttn_Post_8 | -0.044658 | 0.49063 | 0.015317 | 0.87166 |
| 7Networks_RH_DorsAttn_Post_9 | 0.0091581 | 0.88671 | -0.026618 | 0.78467 |
| 7Networks_RH_DorsAttn_Post_10 | -0.055311 | 0.39592 | 0.016689 | 0.84876 |
| 7Networks_RH_DorsAttn_FEF_1 | 0.072076 | 0.27177 | -0.15759 | 0.13453 |
| 7Networks_RH_DorsAttn_FEF_2 | 0.08876 | 0.15261 | -0.26742 | 0.0261 |
| 7Networks_RH_DorsAttn_PrCv_1 | -0.026636 | 0.67973 | 0.028124 | 0.78175 |
| 7Networks_RH_SalVentAttn_TempOccPar_1 | 0.025323 | 0.69037 | 0.17683 | 0.13029 |
| 7Networks_RH_SalVentAttn_TempOccPar_2 | 0.090708 | 0.13254 | -0.10111 | 0.44799 |
| 7Networks_RH_SalVentAttn_TempOccPar_3 | 0.005516 | 0.92941 | 0.032502 | 0.77013 |
| 7Networks_RH_SalVentAttn_PrC_1 | -0.015161 | 0.80795 | 0.070134 | 0.61581 |
| 7Networks_RH_SalVentAttn_FrOper_1 | -0.03008 | 0.64129 | 0.25572 | 0.017876 |
| 7Networks_RH_SalVentAttn_FrOper_2 | 0.14804 | 0.032053 | -0.0005382 | 0.99459 |
| 7Networks_RH_SalVentAttn_FrOper_3 | -0.10011 | 0.10956 | 0.14946 | 0.13266 |
| 7Networks_RH_SalVentAttn_FrOper_4 | -0.091993 | 0.14994 | 0.25337 | 0.021244 |
| 7Networks_RH_SalVentAttn_Med_1 | 0.083248 | 0.18554 | -0.14304 | 0.19007 |
| 7Networks_RH_SalVentAttn_Med_2 | -0.0069754 | 0.91502 | 0.01306 | 0.88142 |
| 7Networks_RH_SalVentAttn_Med_3 | 0.14057 | 0.034752 | -0.10546 | 0.29909 |
| 7Networks_RH_Limbic_OFC_1 | 0.0028532 | 0.96657 | 0.026906 | 0.76803 |
| 7Networks_RH_Limbic_OFC_2 | -0.079244 | 0.20085 | 0.13042 | 0.15977 |
| 7Networks_RH_Limbic_OFC_3 | 0.021672 | 0.74012 | -0.1467 | 0.11436 |
| 7Networks_RH_Limbic_TempPole_1 | -0.0043151 | 0.94411 | 0.36748 | 0.0045149 |
| 7Networks_RH_Limbic_TempPole_2 | -0.048192 | 0.46232 | 0.15112 | 0.11722 |
| 7Networks_RH_Limbic_TempPole_3 | -0.018937 | 0.78204 | 0.011975 | 0.88168 |
| 7Networks_RH_Cont_Par_1 | 0.034226 | 0.58469 | -0.040601 | 0.77741 |
| 7Networks_RH_Cont_Par_2 | 0.046798 | 0.44736 | -0.11663 | 0.34767 |
| 7Networks_RH_Cont_Par_3 | -0.014755 | 0.80977 | -0.065073 | 0.66701 |
| 7Networks_RH_Cont_Temp_1 | 0.029293 | 0.65215 | -0.013151 | 0.91389 |
| 7Networks_RH_Cont_PFCv_1 | -0.043185 | 0.49524 | 0.019203 | 0.87688 |
| 7Networks_RH_Cont_PFCl_1 | -0.010404 | 0.86738 | -0.015973 | 0.88071 |
| 7Networks_RH_Cont_PFCl_2 | -0.025513 | 0.68598 | -0.11958 | 0.23628 |
| 7Networks_RH_Cont_PFCl_3 | 0.0057649 | 0.92596 | -0.14242 | 0.13926 |
| 7Networks_RH_Cont_PFCl_4 | -0.099027 | 0.13556 | 0.080294 | 0.35062 |
| 7Networks_RH_Cont_PFCl_5 | -0.052868 | 0.39574 | -0.15745 | 0.10193 |
| 7Networks_RH_Cont_PFCl_6 | -0.0054907 | 0.92893 | -0.16166 | 0.098969 |
| 7Networks_RH_Cont_PFCl_7 | 0.010729 | 0.87195 | 0.0042544 | 0.96512 |
| 7Networks_RH_Cont_pCun_1 | 0.038379 | 0.53765 | 0.014394 | 0.89477 |
| 7Networks_RH_Cont_PFCmp_1 | -0.10307 | 0.11252 | 0.048371 | 0.55242 |
| 7Networks_RH_Cont_PFCmp_2 | -0.052259 | 0.43139 | 0.11205 | 0.2053 |
| 7Networks_RH_Cont_PFCmp_3 | 0.028641 | 0.65387 | -0.1681 | 0.12154 |
| 7Networks_RH_Cont_PFCmp_4 | 0.085144 | 0.21313 | -0.11733 | 0.17645 |
| 7Networks_RH_Default_Par_1 | 0.033991 | 0.59721 | 0.022103 | 0.85441 |
| 7Networks_RH_Default_Par_2 | -0.023787 | 0.70465 | 0.12054 | 0.33341 |
| 7Networks_RH_Default_Par_3 | 0.06102 | 0.33099 | -0.10825 | 0.38287 |
| 7Networks_RH_Default_Temp_1 | -0.065993 | 0.30308 | 0.19809 | 0.043939 |
| 7Networks_RH_Default_Temp_2 | -0.067774 | 0.29184 | 0.11654 | 0.24423 |
| 7Networks_RH_Default_Temp_3 | -0.075433 | 0.27279 | 0.072447 | 0.4165 |
| 7Networks_RH_Default_Temp_4 | -0.01913 | 0.76009 | 0.02613 | 0.79735 |
| 7Networks_RH_Default_Temp_5 | 0.0039176 | 0.9521 | 0.24243 | 0.034091 |
| 7Networks_RH_Default_PFCv_1 | -0.043205 | 0.51109 | 0.13676 | 0.14697 |
| 7Networks_RH_Default_PFCm_1 | -0.086902 | 0.16127 | 0.067947 | 0.50731 |
| 7Networks_RH_Default_PFCm_2 | 0.0024132 | 0.96919 | -0.041556 | 0.70862 |
| 7Networks_RH_Default_PFCm_3 | 0.03672 | 0.59045 | 0.080296 | 0.42046 |
| 7Networks_RH_Default_PFCm_4 | -0.0015703 | 0.98174 | -0.14564 | 0.055887 |
| 7Networks_RH_Default_PFCm_5 | 0.13419 | 0.040708 | -0.23767 | 0.0028657 |
| 7Networks_RH_Default_PFCm_6 | 0.086476 | 0.17274 | -0.25749 | 0.0086362 |
| 7Networks_RH_Default_PFCm_7 | 0.03059 | 0.61364 | -0.16213 | 0.211 |
| 7Networks_RH_Default_PCC_1 | 0.0012425 | 0.98533 | 0.18947 | 0.045664 |
| 7Networks_RH_Default_PCC_2 | 0.076534 | 0.25585 | 0.056522 | 0.53083 |
| 7Networks_RH_Default_PCC_3 | 0.0075094 | 0.90336 | -0.068331 | 0.54846 |

**Supplementary Table 10. Genetic correlation between Openness and cortical thickness.** Genetic correlation between Openness and local cortical thickness calculated using solar 8.4.0. Regions are named according to the Schaefer 200 atlas, based on 7-networks. Here we report environmental correlation (ρ_e_) and genetic correlation (ρ_g_) and the associated p-values.

| ROIS | h2 | p |
| --- | --- | --- |
| 7Networks_LH_Vis_1 | 0.6006 | 2,40E-23 |
| 7Networks_LH_Vis_2 | 0.52732 | 4,50E-16 |
| 7Networks_LH_Vis_3 | 0.36554 | 4,98E-08 |
| 7Networks_LH_Vis_4 | 0.4093 | 7,94E-09 |
| 7Networks_LH_Vis_5 | 0.31529 | 1,13E-04 |
| 7Networks_LH_Vis_6 | 0.62576 | 8,30E-25 |
| 7Networks_LH_Vis_7 | 0.61774 | 2,61E-26 |
| 7Networks_LH_Vis_8 | 0.36974 | 1,04E-07 |
| 7Networks_LH_Vis_9 | 0.44971 | 2,92E-14 |
| 7Networks_LH_Vis_10 | 0.81315 | 4,67E-58 |
| 7Networks_LH_Vis_11 | 0.45706 | 1,13E-14 |
| 7Networks_LH_Vis_12 | 0.57605 | 9,43E-23 |
| 7Networks_LH_Vis_13 | 0.64077 | 1,16E-28 |
| 7Networks_LH_Vis_14 | 0.46439 | 8,69E-13 |
| 7Networks_LH_SomMot_1 | 0.47555 | 1,65E-14 |
| 7Networks_LH_SomMot_2 | 0.50429 | 7,37E-19 |
| 7Networks_LH_SomMot_3 | 0.41359 | 4,58E-11 |
| 7Networks_LH_SomMot_4 | 0.32472 | 1,11E-05 |
| 7Networks_LH_SomMot_5 | 0.41953 | 7,96E-10 |
| 7Networks_LH_SomMot_6 | 0.47561 | 7,16E-17 |
| 7Networks_LH_SomMot_7 | 0.26654 | 3,00E-07 |
| 7Networks_LH_SomMot_8 | 0.46299 | 6,07E-15 |
| 7Networks_LH_SomMot_9 | 0.2167 | 1.31e-05 |
| 7Networks_LH_SomMot_10 | 0.35846 | 1,45E-08 |
| 7Networks_LH_SomMot_11 | 0.39431 | 4,63E-08 |
| 7Networks_LH_SomMot_12 | 0.46338 | 1,54E-16 |
| 7Networks_LH_SomMot_13 | 0.407 | 3,00E-09 |
| 7Networks_LH_SomMot_14 | 0.29839 | 5,33E-05 |
| 7Networks_LH_SomMot_15 | 0.40605 | 1,37E-10 |
| 7Networks_LH_SomMot_16 | 0.39266 | 2,01E-11 |
| 7Networks_LH_DorsAttn_Post_1 | 0.35377 | 2,78E-07 |
| 7Networks_LH_DorsAttn_Post_2 | 0.39186 | 4,21E-09 |
| 7Networks_LH_DorsAttn_Post_3 | 0.4686 | 7,76E-12 |
| 7Networks_LH_DorsAttn_Post_4 | 0.36877 | 2,43E-07 |
| 7Networks_LH_DorsAttn_Post_5 | 0.2174 | 2.28e-05 |
| 7Networks_LH_DorsAttn_Post_6 | 0.19368 | 7.17e-05 |
| 7Networks_LH_DorsAttn_Post_7 | 0.34452 | 4,19E-07 |
| 7Networks_LH_DorsAttn_Post_8 | 0.31345 | 1,57E-05 |
| 7Networks_LH_DorsAttn_Post_9 | 0.42219 | 7,51E-12 |
| 7Networks_LH_DorsAttn_Post_10 | 0.35749 | 1,65E-07 |
| 7Networks_LH_DorsAttn_FEF_1 | 0.2743 | 2,76E-04 |
| 7Networks_LH_DorsAttn_FEF_2 | 0.24563 | 9,00E-07 |
| 7Networks_LH_DorsAttn_PrCv_1 | 0.31883 | 7,84E-05 |
| 7Networks_LH_SalVentAttn_ParOper_1 | 0.3763 | 1,23E-07 |
| 7Networks_LH_SalVentAttn_ParOper_2 | 0.42416 | 6,25E-12 |
| 7Networks_LH_SalVentAttn_ParOper_3 | 0.31805 | 4,62E-06 |
| 7Networks_LH_SalVentAttn_FrOper_1 | 0.6105 | 3,33E-25 |
| 7Networks_LH_SalVentAttn_FrOper_2 | 0.56057 | 2,97E-25 |
| 7Networks_LH_SalVentAttn_FrOper_3 | 0.50991 | 1,50E-18 |
| 7Networks_LH_SalVentAttn_FrOper_4 | 0.39745 | 4,07E-08 |
| 7Networks_LH_SalVentAttn_PFCl_1 | 0.28579 | 1.3e-06 |
| 7Networks_LH_SalVentAttn_Med_1 | 0.54355 | 1,99E-19 |
| 7Networks_LH_SalVentAttn_Med_2 | 0.45812 | 1,95E-13 |
| 7Networks_LH_SalVentAttn_Med_3 | 0.33891 | 4,94E-05 |
| 7Networks_LH_Limbic_OFC_1 | 0.58907 | 2,26E-28 |
| 7Networks_LH_Limbic_OFC_2 | 0.57848 | 2,20E-23 |
| 7Networks_LH_Limbic_TempPole_1 | 0.54323 | 9,76E-20 |
| 7Networks_LH_Limbic_TempPole_2 | 0.43688 | 1,21E-13 |
| 7Networks_LH_Limbic_TempPole_3 | 0.40956 | 1,46E-10 |
| 7Networks_LH_Limbic_TempPole_4 | 0.60667 | 1,13E-25 |
| 7Networks_LH_Cont_Par_1 | 0.23828 | 2.32e-05 |
| 7Networks_LH_Cont_Par_2 | 0.21504 | 4.17e-05 |
| 7Networks_LH_Cont_Par_3 | 0.19393 | 0.0001607 |
| 7Networks_LH_Cont_Temp_1 | 0.32912 | 6,44E-06 |
| 7Networks_LH_Cont_PFCl_1 | 0.47049 | 9,15E-14 |
| 7Networks_LH_Cont_PFCl_2 | 0.45638 | 2,22E-12 |
| 7Networks_LH_Cont_PFCl_3 | 0.3855 | 3,67E-07 |
| 7Networks_LH_Cont_PFCl_4 | 0.40703 | 1,68E-09 |
| 7Networks_LH_Cont_PFCl_5 | 0.33369 | 2,40E-06 |
| 7Networks_LH_Cont_PFCl_6 | 0.22579 | 1.19e-05 |
| 7Networks_LH_Cont_pCun_1 | 0.49605 | 1,18E-14 |
| 7Networks_LH_Cont_Cing_1 | 0.50214 | 2,09E-15 |
| 7Networks_LH_Cont_Cing_2 | 0.32673 | 1,49E-06 |
| 7Networks_LH_Default_Temp_1 | 0.6418 | 2,20E-29 |
| 7Networks_LH_Default_Temp_2 | 0.33919 | 2,82E-06 |
| 7Networks_LH_Default_Temp_3 | 0.38831 | 1,46E-10 |
| 7Networks_LH_Default_Temp_4 | 0.47738 | 6,29E-15 |
| 7Networks_LH_Default_Temp_5 | 0.33322 | 2,13E-06 |
| 7Networks_LH_Default_Temp_6 | 0.3563 | 2,23E-06 |
| 7Networks_LH_Default_Temp_7 | 0.28457 | 6,00E-07 |
| 7Networks_LH_Default_Temp_8 | 0.14798 | 0.002916 |
| 7Networks_LH_Default_Temp_9 | 0.33789 | 3,25E-06 |
| 7Networks_LH_Default_PFC_1 | 0.43855 | 4,18E-15 |
| 7Networks_LH_Default_PFC_2 | 0.58646 | 2,73E-28 |
| 7Networks_LH_Default_PFC_3 | 0.43535 | 6,11E-13 |
| 7Networks_LH_Default_PFC_4 | 0.49262 | 5,88E-15 |
| 7Networks_LH_Default_PFC_5 | 0.38363 | 4,31E-10 |
| 7Networks_LH_Default_PFC_6 | 0.58969 | 5,04E-25 |
| 7Networks_LH_Default_PFC_7 | 0.43194 | 3,00E-13 |
| 7Networks_LH_Default_PFC_8 | 0.43254 | 3,22E-11 |
| 7Networks_LH_Default_PFC_9 | 0.38884 | 2,94E-07 |
| 7Networks_LH_Default_PFC_10 | 0.45365 | 1,02E-12 |
| 7Networks_LH_Default_PFC_11 | 0.34174 | 8,11E-07 |
| 7Networks_LH_Default_PFC_12 | 0.30495 | 2,34E-04 |
| 7Networks_LH_Default_PFC_13 | 0.31747 | 2,00E-07 |
| 7Networks_LH_Default_PCC_1 | 0.70108 | 9,20E-40 |
| 7Networks_LH_Default_PCC_2 | 0.47296 | 1,06E-13 |
| 7Networks_LH_Default_PCC_3 | 0.36687 | 1,07E-07 |
| 7Networks_LH_Default_PCC_4 | 0.40502 | 6,27E-09 |
| 7Networks_LH_Default_PHC_1 | 0.55641 | 1,22E-24 |
| 7Networks_RH_Vis_1 | 0.53593 | 9,37E-21 |
| 7Networks_RH_Vis_2 | 0.50592 | 6,08E-16 |
| 7Networks_RH_Vis_3 | 0.42698 | 2,27E-10 |
| 7Networks_RH_Vis_4 | 0.50718 | 4,49E-18 |
| 7Networks_RH_Vis_5 | 0.2593 | 1.7e-06 |
| 7Networks_RH_Vis_6 | 0.66068 | 2,98E-25 |
| 7Networks_RH_Vis_7 | 0.61861 | 1,35E-27 |
| 7Networks_RH_Vis_8 | 0.40532 | 9,87E-09 |
| 7Networks_RH_Vis_9 | 0.76834 | 2,40E-45 |
| 7Networks_RH_Vis_10 | 0.78665 | 5,68E-49 |
| 7Networks_RH_Vis_11 | 0.27566 | 3,00E-07 |
| 7Networks_RH_Vis_12 | 0.53112 | 2,68E-19 |
| 7Networks_RH_Vis_13 | 0.51609 | 6,69E-17 |
| 7Networks_RH_Vis_14 | 0.41268 | 1,11E-09 |
| 7Networks_RH_Vis_15 | 0.3054 | 1,63E-04 |
| 7Networks_RH_SomMot_1 | 0.60607 | 1,86E-27 |
| 7Networks_RH_SomMot_2 | 0.57569 | 8,19E-22 |
| 7Networks_RH_SomMot_3 | 0.49611 | 1,10E-14 |
| 7Networks_RH_SomMot_4 | 0.4721 | 3,77E-15 |
| 7Networks_RH_SomMot_5 | 0.28577 | 1,00E-07 |
| 7Networks_RH_SomMot_6 | 0.48915 | 5,34E-14 |
| 7Networks_RH_SomMot_7 | 0.41646 | 1,50E-10 |
| 7Networks_RH_SomMot_8 | 0.33997 | 6,13E-06 |
| 7Networks_RH_SomMot_9 | 0.20591 | 6.29e-05 |
| 7Networks_RH_SomMot_10 | 0.30525 | 3,57E-05 |
| 7Networks_RH_SomMot_11 | 0.53654 | 1.12e-22 |
| 7Networks_RH_SomMot_12 | 0.36916 | 8,27E-09 |
| 7Networks_RH_SomMot_13 | 0.34898 | 3,83E-06 |
| 7Networks_RH_SomMot_14 | 0.44949 | 9,15E-14 |
| 7Networks_RH_SomMot_15 | 0.20415 | 7.03e-05 |
| 7Networks_RH_SomMot_16 | 0.30143 | 1,00E-07 |
| 7Networks_RH_SomMot_17 | 0.16801 | 0.0012705 |
| 7Networks_RH_SomMot_18 | 0.46965 | 1,48E-14 |
| 7Networks_RH_SomMot_19 | 0.43103 | 7,22E-13 |
| 7Networks_RH_DorsAttn_Post_1 | 0.48661 | 2,16E-16 |
| 7Networks_RH_DorsAttn_Post_2 | 0.32684 | 1,14E-05 |
| 7Networks_RH_DorsAttn_Post_3 | 0.39992 | 5,21E-11 |
| 7Networks_RH_DorsAttn_Post_4 | 0.23662 | 4.5e-06 |
| 7Networks_RH_DorsAttn_Post_5 | 0.28345 | 1,00E-07 |
| 7Networks_RH_DorsAttn_Post_6 | 0.25298 | 2.2e-06 |
| 7Networks_RH_DorsAttn_Post_7 | 0.23287 | 2.49e-05 |
| 7Networks_RH_DorsAttn_Post_8 | 0.23369 | 1.2e-05 |
| 7Networks_RH_DorsAttn_Post_9 | 0.41653 | 3,52E-11 |
| 7Networks_RH_DorsAttn_Post_10 | 0.36988 | 3,62E-08 |
| 7Networks_RH_DorsAttn_FEF_1 | 0.20897 | 8.73e-05 |
| 7Networks_RH_DorsAttn_FEF_2 | 0.54616 | 1,55E-19 |
| 7Networks_RH_DorsAttn_PrCv_1 | 0.32573 | 9,19E-06 |
| 7Networks_RH_SalVentAttn_TempOccPar_1 | 0.32273 | 4,43E-05 |
| 7Networks_RH_SalVentAttn_TempOccPar_2 | 0.21492 | 2.66e-05 |
| 7Networks_RH_SalVentAttn_TempOccPar_3 | 0.40254 | 3,61E-09 |
| 7Networks_RH_SalVentAttn_PrC_1 | 0.28167 | 2,00E-07 |
| 7Networks_RH_SalVentAttn_FrOper_1 | 0.56851 | 2,02E-23 |
| 7Networks_RH_SalVentAttn_FrOper_2 | 0.5442 | 2,20E-20 |
| 7Networks_RH_SalVentAttn_FrOper_3 | 0.41808 | 1,16E-11 |
| 7Networks_RH_SalVentAttn_FrOper_4 | 0.46186 | 1,11E-13 |
| 7Networks_RH_SalVentAttn_Med_1 | 0.57864 | 8,50E-22 |
| 7Networks_RH_SalVentAttn_Med_2 | 0.51585 | 8,63E-19 |
| 7Networks_RH_SalVentAttn_Med_3 | 0.48762 | 4,35E-15 |
| 7Networks_RH_Limbic_OFC_1 | 0.54485 | 5,67E-20 |
| 7Networks_RH_Limbic_OFC_2 | 0.48934 | 1,93E-18 |
| 7Networks_RH_Limbic_OFC_3 | 0.44581 | 1,93E-09 |
| 7Networks_RH_Limbic_TempPole_1 | 0.48102 | 2,78E-17 |
| 7Networks_RH_Limbic_TempPole_2 | 0.45362 | 1,79E-14 |
| 7Networks_RH_Limbic_TempPole_3 | 0.45799 | 5,51E-14 |
| 7Networks_RH_Cont_Par_1 | 0.27408 | 2,00E-07 |
| 7Networks_RH_Cont_Par_2 | 0.21241 | 0.0001276 |
| 7Networks_RH_Cont_Par_3 | 0.20691 | 4.23e-05 |
| 7Networks_RH_Cont_Temp_1 | 0.36999 | 1,11E-06 |
| 7Networks_RH_Cont_PFCv_1 | 0.30523 | 3,00E-05 |
| 7Networks_RH_Cont_PFCl_1 | 0.52348 | 5,21E-21 |
| 7Networks_RH_Cont_PFCl_2 | 0.33427 | 6,41E-05 |
| 7Networks_RH_Cont_PFCl_3 | 0.41058 | 7,47E-10 |
| 7Networks_RH_Cont_PFCl_4 | 0.43111 | 2,70E-12 |
| 7Networks_RH_Cont_PFCl_5 | 0.31826 | 7,35E-05 |
| 7Networks_RH_Cont_PFCl_6 | 0.28379 | 1,00E-07 |
| 7Networks_RH_Cont_PFCl_7 | 0.3732 | 1,82E-09 |
| 7Networks_RH_Cont_pCun_1 | 0.43375 | 4,36E-10 |
| 7Networks_RH_Cont_PFCmp_1 | 0.47566 | 1,03E-14 |
| 7Networks_RH_Cont_PFCmp_2 | 0.39914 | 2,61E-09 |
| 7Networks_RH_Cont_PFCmp_3 | 0.56241 | 2,22E-21 |
| 7Networks_RH_Cont_PFCmp_4 | 0.44582 | 1,25E-11 |
| 7Networks_RH_Default_Par_1 | 0.32937 | 3,47E-05 |
| 7Networks_RH_Default_Par_2 | 0.19904 | 0.0001579 |
| 7Networks_RH_Default_Par_3 | 0.17497 | 0.0008717 |
| 7Networks_RH_Default_Temp_1 | 0.47657 | 7,43E-17 |
| 7Networks_RH_Default_Temp_2 | 0.41053 | 3,27E-11 |
| 7Networks_RH_Default_Temp_3 | 0.39808 | 8,79E-12 |
| 7Networks_RH_Default_Temp_4 | 0.35559 | 2,80E-07 |
| 7Networks_RH_Default_Temp_5 | 0.4521 | 5,39E-13 |
| 7Networks_RH_Default_PFCv_1 | 0.35218 | 2,55E-08 |
| 7Networks_RH_Default_PFCm_1 | 0.59353 | 5,75E-22 |
| 7Networks_RH_Default_PFCm_2 | 0.5114 | 1,37E-14 |
| 7Networks_RH_Default_PFCm_3 | 0.18881 | 0.0009007 |
| 7Networks_RH_Default_PFCm_4 | 0.5631 | 8,06E-20 |
| 7Networks_RH_Default_PFCm_5 | 0.3668 | 4,03E-09 |
| 7Networks_RH_Default_PFCm_6 | 0.33739 | 3,57E-06 |
| 7Networks_RH_Default_PFCm_7 | 0.29149 | 1,00E-07 |
| 7Networks_RH_Default_PCC_1 | 0.66787 | 1,20E-32 |
| 7Networks_RH_Default_PCC_2 | 0.40918 | 5,43E-09 |
| 7Networks_RH_Default_PCC_3 | 0.32893 | 1,29E-04 |

**Supplementary Table 11. Heritability of surface area.** Heritability of surface area calculated using solar 8.4.0. regions are named according to the Schaefer 200 atlas, based on 7-networks. Here we report heritability values, standard errors (SE), and p-values. All regions were significantly heritable at FDRq<0.05. Nomenclature of the regions is based on the official parcel names of the Schaefer 200 – 7 networks parcel solution.

| ROIS | ρ_e_ | p | ρ_g_ | p |
| --- | --- | --- | --- | --- |
| 7Networks_LH_Vis_1 | -0.030928 | 0.66109 | 0.045374 | 0.67995 |
| 7Networks_LH_Vis_2 | -0.12482 | 0.0694 | 0.12922 | 0.28403 |
| 7Networks_LH_Vis_3 | -0.098226 | 0.11137 | 0.31493 | 0.025786 |
| 7Networks_LH_Vis_4 | -0.12779 | 0.048835 | 0.31229 | 0.020927 |
| 7Networks_LH_Vis_5 | -0.076592 | 0.21456 | 0.072776 | 0.63048 |
| 7Networks_LH_Vis_6 | 0.0066246 | 0.92689 | -0.12719 | 0.2438 |
| 7Networks_LH_Vis_7 | 0.0067707 | 0.92277 | 0.022167 | 0.83463 |
| 7Networks_LH_Vis_8 | -0.0073002 | 0.90735 | 0.068315 | 0.62281 |
| 7Networks_LH_Vis_9 | -0.047945 | 0.45002 | -0.0003375 | 0.99701 |
| 7Networks_LH_Vis_10 | 0.065372 | 0.40082 | -0.0091931 | 0.91354 |
| 7Networks_LH_Vis_11 | -0.05668 | 0.38075 | 0.10387 | 0.40019 |
| 7Networks_LH_Vis_12 | 0.089871 | 0.18435 | -0.036313 | 0.74212 |
| 7Networks_LH_Vis_13 | -0.0062731 | 0.92959 | 0.070556 | 0.50013 |
| 7Networks_LH_Vis_14 | -0.095079 | 0.14673 | 0.12743 | 0.32014 |
| 7Networks_LH_SomMot_1 | 0.097818 | 0.12941 | -0.14353 | 0.23337 |
| 7Networks_LH_SomMot_2 | -0.022765 | 0.72219 | -0.0098379 | 0.93118 |
| 7Networks_LH_SomMot_3 | -0.031675 | 0.6128 | 0.085213 | 0.50449 |
| 7Networks_LH_SomMot_4 | -0.1121 | 0.06246 | 0.32454 | 0.026767 |
| 7Networks_LH_SomMot_5 | -0.058738 | 0.36222 | 0.085849 | 0.51703 |
| 7Networks_LH_SomMot_6 | 0.020959 | 0.73913 | -0.044739 | 0.70052 |
| 7Networks_LH_SomMot_7 | -0.09445 | 0.1121 | 0.31656 | 0.044486 |
| 7Networks_LH_SomMot_8 | 0.022157 | 0.72474 | 0.015357 | 0.89717 |
| 7Networks_LH_SomMot_9 | -0.046123 | 0.42295 | 0.062421 | 0.71688 |
| 7Networks_LH_SomMot_10 | -0.0090494 | 0.88056 | -0.037057 | 0.78259 |
| 7Networks_LH_SomMot_11 | 0.036415 | 0.57373 | 0.072707 | 0.59889 |
| 7Networks_LH_SomMot_12 | -0.063156 | 0.3046 | 0.28286 | 0.018938 |
| 7Networks_LH_SomMot_13 | 0.0047618 | 0.94066 | -0.021909 | 0.86992 |
| 7Networks_LH_SomMot_14 | 0.040007 | 0.49988 | 0.0082763 | 0.95526 |
| 7Networks_LH_SomMot_15 | -0.021376 | 0.73097 | 0.10726 | 0.40747 |
| 7Networks_LH_SomMot_16 | 0.012034 | 0.84439 | 0.10102 | 0.44114 |
| 7Networks_LH_DorsAttn_Post_1 | -0.072299 | 0.2418 | 0.14957 | 0.28909 |
| 7Networks_LH_DorsAttn_Post_2 | 0.058433 | 0.35297 | -0.030711 | 0.81811 |
| 7Networks_LH_DorsAttn_Post_3 | -0.034041 | 0.61151 | 0.013965 | 0.9134 |
| 7Networks_LH_DorsAttn_Post_4 | -0.051902 | 0.40934 | 0.0024049 | 0.986 |
| 7Networks_LH_DorsAttn_Post_5 | -0.032912 | 0.56926 | 0.14521 | 0.4031 |
| 7Networks_LH_DorsAttn_Post_6 | 0.08139 | 0.14618 | -0.26876 | 0.13345 |
| 7Networks_LH_DorsAttn_Post_7 | -0.040875 | 0.49833 | -0.036083 | 0.79484 |
| 7Networks_LH_DorsAttn_Post_8 | 0.088346 | 0.13716 | -0.19384 | 0.18092 |
| 7Networks_LH_DorsAttn_Post_9 | -0.0052027 | 0.93388 | 0.034154 | 0.78781 |
| 7Networks_LH_DorsAttn_Post_10 | -0.045717 | 0.45789 | 0.13746 | 0.3157 |
| 7Networks_LH_DorsAttn_FEF_1 | 0.0016306 | 0.97735 | 0.19598 | 0.19503 |
| 7Networks_LH_DorsAttn_FEF_2 | 0.00187 | 0.97427 | 0.15158 | 0.35198 |
| 7Networks_LH_DorsAttn_PrCv_1 | -0.032533 | 0.59715 | 0.17597 | 0.24503 |
| 7Networks_LH_SalVentAttn_ParOper_1 | -0.11553 | 0.064708 | 0.18599 | 0.1889 |
| 7Networks_LH_SalVentAttn_ParOper_2 | -0.056488 | 0.36508 | 0.0034846 | 0.97783 |
| 7Networks_LH_SalVentAttn_ParOper_3 | -0.074301 | 0.22359 | 0.18795 | 0.21325 |
| 7Networks_LH_SalVentAttn_FrOper_1 | 0.06732 | 0.33476 | -0.073175 | 0.49307 |
| 7Networks_LH_SalVentAttn_FrOper_2 | -0.082541 | 0.20264 | 0.1651 | 0.11756 |
| 7Networks_LH_SalVentAttn_FrOper_3 | -0.10253 | 0.11092 | 0.1694 | 0.14686 |
| 7Networks_LH_SalVentAttn_FrOper_4 | 0.0026363 | 0.96725 | 0.16846 | 0.22165 |
| 7Networks_LH_SalVentAttn_PFCl_1 | -0.13694 | 0.029757 | 0.19647 | 0.25538 |
| 7Networks_LH_SalVentAttn_Med_1 | -0.023697 | 0.72308 | 0.053112 | 0.63984 |
| 7Networks_LH_SalVentAttn_Med_2 | 0.026756 | 0.67844 | 0.0098467 | 0.93676 |
| 7Networks_LH_SalVentAttn_Med_3 | -0.032985 | 0.60415 | 0.043144 | 0.77416 |
| 7Networks_LH_Limbic_OFC_1 | -0.12498 | 0.058766 | 0.1311 | 0.2055 |
| 7Networks_LH_Limbic_OFC_2 | 0.0023688 | 0.97232 | -0.10061 | 0.35693 |
| 7Networks_LH_Limbic_TempPole_1 | 0.016413 | 0.80701 | 0.048984 | 0.66741 |
| 7Networks_LH_Limbic_TempPole_2 | -0.037151 | 0.55025 | 0.07606 | 0.53619 |
| 7Networks_LH_Limbic_TempPole_3 | -0.024961 | 0.69099 | 0.14303 | 0.26588 |
| 7Networks_LH_Limbic_TempPole_4 | -0.012266 | 0.85875 | -0.075688 | 0.4763 |
| 7Networks_LH_Cont_Par_1 | -0.070941 | 0.24278 | 0.16567 | 0.34868 |
| 7Networks_LH_Cont_Par_2 | -0.030006 | 0.60985 | 0.12635 | 0.47282 |
| 7Networks_LH_Cont_Par_3 | 0.030373 | 0.59934 | -0.057288 | 0.75446 |
| 7Networks_LH_Cont_Temp_1 | -0.047103 | 0.43741 | 0.14873 | 0.30905 |
| 7Networks_LH_Cont_PFCl_1 | -0.090404 | 0.16535 | 0.10546 | 0.40451 |
| 7Networks_LH_Cont_PFCl_2 | -0.04415 | 0.49872 | -0.01099 | 0.93057 |
| 7Networks_LH_Cont_PFCl_3 | -0.14414 | 0.028397 | 0.31706 | 0.023582 |
| 7Networks_LH_Cont_PFCl_4 | -0.0008191 | 0.98822 | -0.041523 | 0.75456 |
| 7Networks_LH_Cont_PFCl_5 | 0.068352 | 0.26388 | 0.021857 | 0.87856 |
| 7Networks_LH_Cont_PFCl_6 | -0.010339 | 0.85957 | 0.074746 | 0.66207 |
| 7Networks_LH_Cont_pCun_1 | 0.043914 | 0.51147 | 0.031915 | 0.79404 |
| 7Networks_LH_Cont_Cing_1 | -0.091354 | 0.16913 | 0.24613 | 0.04664 |
| 7Networks_LH_Cont_Cing_2 | -0.038107 | 0.53505 | 0.10667 | 0.47458 |
| 7Networks_LH_Default_Temp_1 | -0.10397 | 0.14203 | 0.18337 | 0.07783 |
| 7Networks_LH_Default_Temp_2 | -0.085602 | 0.16104 | 0.25846 | 0.070163 |
| 7Networks_LH_Default_Temp_3 | -0.021488 | 0.72507 | 0.022147 | 0.86518 |
| 7Networks_LH_Default_Temp_4 | 0.040685 | 0.53201 | 0.020557 | 0.8655 |
| 7Networks_LH_Default_Temp_5 | -0.055403 | 0.3618 | 0.19556 | 0.17395 |
| 7Networks_LH_Default_Temp_6 | 0.028704 | 0.65064 | 0.11609 | 0.42081 |
| 7Networks_LH_Default_Temp_7 | 0.0075942 | 0.9019 | -0.017337 | 0.91361 |
| 7Networks_LH_Default_Temp_8 | -0.083582 | 0.14143 | 0.39502 | 0.068803 |
| 7Networks_LH_Default_Temp_9 | 0.018389 | 0.76358 | 0.0888 | 0.53259 |
| 7Networks_LH_Default_PFC_1 | -0.065343 | 0.28231 | 0.17325 | 0.13857 |
| 7Networks_LH_Default_PFC_2 | -0.11262 | 0.088432 | 0.12634 | 0.22447 |
| 7Networks_LH_Default_PFC_3 | 0.027648 | 0.66026 | 0.069425 | 0.57734 |
| 7Networks_LH_Default_PFC_4 | -0.10762 | 0.10473 | 0.23102 | 0.053147 |
| 7Networks_LH_Default_PFC_5 | 0.002457 | 0.96787 | 0.0096717 | 0.94237 |
| 7Networks_LH_Default_PFC_6 | -0.029716 | 0.66323 | 0.01629 | 0.88014 |
| 7Networks_LH_Default_PFC_7 | -0.070274 | 0.28862 | 0.12644 | 0.34613 |
| 7Networks_LH_Default_PFC_8 | -0.061964 | 0.33989 | 0.030805 | 0.81733 |
| 7Networks_LH_Default_PFC_9 | -0.065824 | 0.31407 | 0.13213 | 0.3409 |
| 7Networks_LH_Default_PFC_10 | 0.036967 | 0.5677 | -0.06933 | 0.57686 |
| 7Networks_LH_Default_PFC_11 | 0.0085065 | 0.88899 | 0.060839 | 0.66674 |
| 7Networks_LH_Default_PFC_12 | 0.016272 | 0.79086 | 0.14511 | 0.34055 |
| 7Networks_LH_Default_PFC_13 | -0.020319 | 0.75334 | 0.071933 | 0.65328 |
| 7Networks_LH_Default_PCC_1 | 0.04991 | 0.49057 | -0.13955 | 0.14508 |
| 7Networks_LH_Default_PCC_2 | -0.024821 | 0.70755 | 0.017311 | 0.89058 |
| 7Networks_LH_Default_PCC_3 | -0.012112 | 0.84561 | 0.16889 | 0.22811 |
| 7Networks_LH_Default_PCC_4 | -0.13558 | 0.033214 | 0.18588 | 0.16454 |
| 7Networks_LH_Default_PHC_1 | 0.046179 | 0.47795 | -0.087322 | 0.41379 |
| 7Networks_RH_Vis_1 | 0.040235 | 0.54185 | -0.12334 | 0.27473 |
| 7Networks_RH_Vis_2 | 0.041745 | 0.52716 | -0.041821 | 0.72518 |
| 7Networks_RH_Vis_3 | -0.002628 | 0.96756 | -0.12209 | 0.34946 |
| 7Networks_RH_Vis_4 | -0.046581 | 0.47213 | 0.23795 | 0.042172 |
| 7Networks_RH_Vis_5 | -0.12598 | 0.034775 | 0.37756 | 0.024389 |
| 7Networks_RH_Vis_6 | -0.13874 | 0.06149 | 0.099819 | 0.34787 |
| 7Networks_RH_Vis_7 | -0.034892 | 0.61679 | 0.025954 | 0.80741 |
| 7Networks_RH_Vis_8 | -0.072142 | 0.26712 | 0.20744 | 0.12763 |
| 7Networks_RH_Vis_9 | 0.021569 | 0.77823 | 0.0052264 | 0.95443 |
| 7Networks_RH_Vis_10 | 0.14608 | 0.064013 | -0.10879 | 0.21648 |
| 7Networks_RH_Vis_11 | 0.01178 | 0.84603 | 0.052068 | 0.74398 |
| 7Networks_RH_Vis_12 | -0.049735 | 0.45229 | 0.15216 | 0.18057 |
| 7Networks_RH_Vis_13 | 0.091747 | 0.16973 | -0.20882 | 0.073044 |
| 7Networks_RH_Vis_14 | 0.02168 | 0.73619 | 0.016691 | 0.90032 |
| 7Networks_RH_Vis_15 | -0.0067054 | 0.91201 | 0.017717 | 0.90619 |
| 7Networks_RH_SomMot_1 | -0.091908 | 0.17846 | -0.0098895 | 0.92554 |
| 7Networks_RH_SomMot_2 | 0.045117 | 0.51071 | -0.068733 | 0.53559 |
| 7Networks_RH_SomMot_3 | -0.072843 | 0.27337 | 0.054595 | 0.65593 |
| 7Networks_RH_SomMot_4 | -0.084327 | 0.18708 | 0.058893 | 0.62538 |
| 7Networks_RH_SomMot_5 | -0.024053 | 0.6882 | 0.18588 | 0.23556 |
| 7Networks_RH_SomMot_6 | -0.20083 | 0.002301 | 0.30822 | 0.016553 |
| 7Networks_RH_SomMot_7 | 0.084442 | 0.17731 | -0.13238 | 0.30225 |
| 7Networks_RH_SomMot_8 | -0.043386 | 0.48119 | 0.093471 | 0.51459 |
| 7Networks_RH_SomMot_9 | -0.13336 | 0.022746 | 0.13711 | 0.4427 |
| 7Networks_RH_SomMot_10 | 0.066209 | 0.26494 | -0.098192 | 0.49949 |
| 7Networks_RH_SomMot_11 | -0.074831 | 0.26473 | 0.037308 | 0.74947 |
| 7Networks_RH_SomMot_12 | -0.015196 | 0.80283 | -0.020399 | 0.87882 |
| 7Networks_RH_SomMot_13 | -0.060649 | 0.33567 | 0.18421 | 0.20573 |
| 7Networks_RH_SomMot_14 | 0.022087 | 0.7277 | 0.052609 | 0.66891 |
| 7Networks_RH_SomMot_15 | -0.045058 | 0.44182 | 0.23318 | 0.19172 |
| 7Networks_RH_SomMot_16 | 0.091614 | 0.13051 | -0.2979 | 0.055549 |
| 7Networks_RH_SomMot_17 | -0.062778 | 0.28416 | 0.35241 | 0.078817 |
| 7Networks_RH_SomMot_18 | -0.078129 | 0.22223 | 0.065291 | 0.59388 |
| 7Networks_RH_SomMot_19 | 0.074308 | 0.2339 | -0.072194 | 0.55743 |
| 7Networks_RH_DorsAttn_Post_1 | -0.080852 | 0.21092 | 0.077662 | 0.52114 |
| 7Networks_RH_DorsAttn_Post_2 | -0.06608 | 0.28131 | 0.24494 | 0.10105 |
| 7Networks_RH_DorsAttn_Post_3 | -0.16929 | 0.0057493 | 0.12721 | 0.31875 |
| 7Networks_RH_DorsAttn_Post_4 | -0.095527 | 0.10102 | 0.30482 | 0.067098 |
| 7Networks_RH_DorsAttn_Post_5 | -0.17429 | 0.003885 | 0.30028 | 0.050363 |
| 7Networks_RH_DorsAttn_Post_6 | -0.0011879 | 0.98416 | 0.062362 | 0.70637 |
| 7Networks_RH_DorsAttn_Post_7 | -0.013578 | 0.8238 | 0.02446 | 0.8906 |
| 7Networks_RH_DorsAttn_Post_8 | 0.048446 | 0.41731 | -0.14592 | 0.39019 |
| 7Networks_RH_DorsAttn_Post_9 | 0.061171 | 0.3374 | -0.072013 | 0.58607 |
| 7Networks_RH_DorsAttn_Post_10 | 0.0084911 | 0.89118 | 0.083211 | 0.54255 |
| 7Networks_RH_DorsAttn_FEF_1 | 0.096089 | 0.10861 | 0.051858 | 0.77678 |
| 7Networks_RH_DorsAttn_FEF_2 | 0.084734 | 0.21035 | 0.056407 | 0.62135 |
| 7Networks_RH_DorsAttn_PrCv_1 | -0.065579 | 0.27767 | 0.18928 | 0.19392 |
| 7Networks_RH_SalVentAttn_TempOccPar_1 | 0.016347 | 0.79103 | -0.074103 | 0.61511 |
| 7Networks_RH_SalVentAttn_TempOccPar_2 | -0.01636 | 0.77814 | -0.045144 | 0.79552 |
| 7Networks_RH_SalVentAttn_TempOccPar_3 | -0.098408 | 0.12014 | 0.18501 | 0.15927 |
| 7Networks_RH_SalVentAttn_PrC_1 | 0.026757 | 0.65621 | 0.063111 | 0.68626 |
| 7Networks_RH_SalVentAttn_FrOper_1 | -0.036725 | 0.58455 | 0.018876 | 0.86238 |
| 7Networks_RH_SalVentAttn_FrOper_2 | -0.026999 | 0.68417 | 0.0066662 | 0.95236 |
| 7Networks_RH_SalVentAttn_FrOper_3 | -0.025237 | 0.68587 | 0.11228 | 0.37473 |
| 7Networks_RH_SalVentAttn_FrOper_4 | -0.054139 | 0.39982 | 0.12478 | 0.31913 |
| 7Networks_RH_SalVentAttn_Med_1 | -0.12017 | 0.080634 | 0.14662 | 0.18603 |
| 7Networks_RH_SalVentAttn_Med_2 | -0.12126 | 0.062327 | 0.11724 | 0.31321 |
| 7Networks_RH_SalVentAttn_Med_3 | -0.049048 | 0.45344 | 0.063466 | 0.60229 |
| 7Networks_RH_Limbic_OFC_1 | -0.083704 | 0.20917 | -0.0020453 | 0.98555 |
| 7Networks_RH_Limbic_OFC_2 | -0.060144 | 0.33894 | 0.022201 | 0.84616 |
| 7Networks_RH_Limbic_OFC_3 | -0.11962 | 0.072785 | 0.21127 | 0.11386 |
| 7Networks_RH_Limbic_TempPole_1 | -0.11084 | 0.078633 | 0.20265 | 0.078907 |
| 7Networks_RH_Limbic_TempPole_2 | 0.050272 | 0.42577 | -0.013407 | 0.91138 |
| 7Networks_RH_Limbic_TempPole_3 | 0.034674 | 0.58774 | 0.024629 | 0.841 |
| 7Networks_RH_Cont_Par_1 | -0.035164 | 0.55744 | 0.067976 | 0.66905 |
| 7Networks_RH_Cont_Par_2 | -0.0094666 | 0.87579 | 0.14714 | 0.42627 |
| 7Networks_RH_Cont_Par_3 | -0.039083 | 0.49747 | 0.11793 | 0.50859 |
| 7Networks_RH_Cont_Temp_1 | -0.073407 | 0.25841 | 0.2018 | 0.1594 |
| 7Networks_RH_Cont_PFCv_1 | -0.079062 | 0.18234 | 0.051086 | 0.7236 |
| 7Networks_RH_Cont_PFCl_1 | -0.043209 | 0.50207 | 0.040032 | 0.71825 |
| 7Networks_RH_Cont_PFCl_2 | 0.010181 | 0.87236 | 0.14392 | 0.33737 |
| 7Networks_RH_Cont_PFCl_3 | -0.05224 | 0.41346 | 0.12405 | 0.34406 |
| 7Networks_RH_Cont_PFCl_4 | -0.12206 | 0.055291 | 0.25385 | 0.053692 |
| 7Networks_RH_Cont_PFCl_5 | -0.01695 | 0.78449 | 0.074164 | 0.62544 |
| 7Networks_RH_Cont_PFCl_6 | -0.033826 | 0.58122 | -0.065231 | 0.68134 |
| 7Networks_RH_Cont_PFCl_7 | -0.075001 | 0.21011 | 0.22148 | 0.098988 |
| 7Networks_RH_Cont_pCun_1 | 0.004197 | 0.94914 | -0.10979 | 0.40647 |
| 7Networks_RH_Cont_PFCmp_1 | -0.07733 | 0.23353 | 0.21531 | 0.086422 |
| 7Networks_RH_Cont_PFCmp_2 | 0.05281 | 0.40481 | -0.0523 | 0.69586 |
| 7Networks_RH_Cont_PFCmp_3 | -0.1496 | 0.026679 | 0.17353 | 0.12117 |
| 7Networks_RH_Cont_PFCmp_4 | -0.007828 | 0.904 | 0.058451 | 0.64831 |
| 7Networks_RH_Default_Par_1 | -0.055779 | 0.37073 | 0.16982 | 0.24348 |
| 7Networks_RH_Default_Par_2 | -0.018605 | 0.75362 | -0.09482 | 0.60902 |
| 7Networks_RH_Default_Par_3 | 0.018702 | 0.75061 | -0.021872 | 0.9125 |
| 7Networks_RH_Default_Temp_1 | -0.029248 | 0.64191 | 0.13072 | 0.26521 |
| 7Networks_RH_Default_Temp_2 | -0.0051599 | 0.93368 | 0.18329 | 0.15309 |
| 7Networks_RH_Default_Temp_3 | -0.044008 | 0.46463 | 0.035169 | 0.77961 |
| 7Networks_RH_Default_Temp_4 | 0.0078569 | 0.89872 | -0.13916 | 0.31406 |
| 7Networks_RH_Default_Temp_5 | -0.070867 | 0.27122 | 0.033243 | 0.79347 |
| 7Networks_RH_Default_PFCv_1 | -0.068462 | 0.25639 | 0.13214 | 0.32681 |
| 7Networks_RH_Default_PFCm_1 | -0.063893 | 0.36399 | 0.0006798 | 0.99508 |
| 7Networks_RH_Default_PFCm_2 | -0.07802 | 0.25478 | 0.06592 | 0.59143 |
| 7Networks_RH_Default_PFCm_3 | -0.069811 | 0.25886 | 0.046511 | 0.81644 |
| 7Networks_RH_Default_PFCm_4 | -0.077408 | 0.25861 | 0.2693 | 0.023979 |
| 7Networks_RH_Default_PFCm_5 | -0.17254 | 0.0046775 | 0.4151 | 0.0022735 |
| 7Networks_RH_Default_PFCm_6 | -0.075253 | 0.21481 | 0.23068 | 0.1161 |
| 7Networks_RH_Default_PFCm_7 | 0.039168 | 0.51906 | 0.1769 | 0.25504 |
| 7Networks_RH_Default_PCC_1 | 0.04563 | 0.52428 | -0.21217 | 0.033669 |
| 7Networks_RH_Default_PCC_2 | -0.04711 | 0.46751 | 0.097667 | 0.47057 |
| 7Networks_RH_Default_PCC_3 | -0.014351 | 0.82052 | -0.068083 | 0.65429 |

**Supplementary Table 12. Genetic and environmental correlation between Agreeableness and surface area in HCP.** Genetic correlation between Agreeableness and local surface area calculated using solar 8.4.0. Regions are named according to the Schaefer 200 atlas, based on 7-networks. Here we report environmental correlation (ρ_e_) and genetic correlation (ρ_g_) and the associated p-values.

| ROIS | ρ_e_ | p | ρ_g_ | p |
| --- | --- | --- | --- | --- |
| 7Networks_LH_Vis_1 | 0.10734 | 0.12855 | -0.13813 | 0.12368 |
| 7Networks_LH_Vis_2 | 0.063607 | 0.36168 | -0.16456 | 0.097395 |
| 7Networks_LH_Vis_3 | 0.041662 | 0.50974 | -0.11786 | 0.30135 |
| 7Networks_LH_Vis_4 | 0.072385 | 0.27667 | -0.18161 | 0.10417 |
| 7Networks_LH_Vis_5 | -0.057788 | 0.36091 | 0.044127 | 0.72087 |
| 7Networks_LH_Vis_6 | -0.14802 | 0.040604 | 0.084543 | 0.34007 |
| 7Networks_LH_Vis_7 | 0.088704 | 0.20525 | -0.0086421 | 0.92088 |
| 7Networks_LH_Vis_8 | -0.074775 | 0.24239 | -0.031264 | 0.78344 |
| 7Networks_LH_Vis_9 | -0.016008 | 0.80485 | -0.027353 | 0.7874 |
| 7Networks_LH_Vis_10 | 0.013029 | 0.86514 | 0.025284 | 0.71457 |
| 7Networks_LH_Vis_11 | -0.038469 | 0.55839 | -0.042168 | 0.67981 |
| 7Networks_LH_Vis_12 | 0.0028014 | 0.96742 | 0.068917 | 0.44132 |
| 7Networks_LH_Vis_13 | 0.033172 | 0.64183 | -0.066166 | 0.44144 |
| 7Networks_LH_Vis_14 | 0.086267 | 0.19681 | -0.076946 | 0.45423 |
| 7Networks_LH_SomMot_1 | -0.017957 | 0.78501 | -0.088227 | 0.37403 |
| 7Networks_LH_SomMot_2 | -0.017094 | 0.79269 | 0.012751 | 0.89216 |
| 7Networks_LH_SomMot_3 | 0.070282 | 0.2711 | -0.025091 | 0.81093 |
| 7Networks_LH_SomMot_4 | -0.091565 | 0.13687 | 0.044672 | 0.7046 |
| 7Networks_LH_SomMot_5 | 0.11253 | 0.090251 | -0.15869 | 0.1351 |
| 7Networks_LH_SomMot_6 | 0.058073 | 0.36004 | -0.14457 | 0.13642 |
| 7Networks_LH_SomMot_7 | 0.0086869 | 0.8848 | -0.1415 | 0.26912 |
| 7Networks_LH_SomMot_8 | 0.058632 | 0.35578 | -0.039986 | 0.6802 |
| 7Networks_LH_SomMot_9 | 0.0001566 | 0.99701 | 0.057111 | 0.6858 |
| 7Networks_LH_SomMot_10 | 0.032468 | 0.59442 | -0.16819 | 0.12502 |
| 7Networks_LH_SomMot_11 | 0.066866 | 0.30696 | -0.084038 | 0.45936 |
| 7Networks_LH_SomMot_12 | -0.029805 | 0.63543 | 0.0019107 | 0.98377 |
| 7Networks_LH_SomMot_13 | 0.012534 | 0.84732 | -0.074658 | 0.49441 |
| 7Networks_LH_SomMot_14 | 0.043516 | 0.47213 | -0.10159 | 0.4003 |
| 7Networks_LH_SomMot_15 | -0.0062225 | 0.92155 | 0.07089 | 0.50258 |
| 7Networks_LH_SomMot_16 | 0.0082208 | 0.89537 | -0.053009 | 0.61792 |
| 7Networks_LH_DorsAttn_Post_1 | 0.11174 | 0.074099 | -0.29007 | 0.013333 |
| 7Networks_LH_DorsAttn_Post_2 | -0.061078 | 0.34067 | -0.099012 | 0.36908 |
| 7Networks_LH_DorsAttn_Post_3 | 0.20426 | 0.002299 | -0.26321 | 0.011616 |
| 7Networks_LH_DorsAttn_Post_4 | -0.017446 | 0.78594 | 0.099665 | 0.3882 |
| 7Networks_LH_DorsAttn_Post_5 | 0.030155 | 0.61142 | 0.0021163 | 0.98806 |
| 7Networks_LH_DorsAttn_Post_6 | -0.019172 | 0.73966 | 0.13645 | 0.35018 |
| 7Networks_LH_DorsAttn_Post_7 | -0.093433 | 0.12718 | 0.23212 | 0.044142 |
| 7Networks_LH_DorsAttn_Post_8 | -0.011896 | 0.84541 | 0.085199 | 0.47373 |
| 7Networks_LH_DorsAttn_Post_9 | 0.13807 | 0.029734 | -0.22241 | 0.034129 |
| 7Networks_LH_DorsAttn_Post_10 | 0.084402 | 0.1756 | -0.20075 | 0.079075 |
| 7Networks_LH_DorsAttn_FEF_1 | -0.035654 | 0.54579 | 0.086709 | 0.48636 |
| 7Networks_LH_DorsAttn_FEF_2 | 0.024882 | 0.67382 | -0.028503 | 0.82835 |
| 7Networks_LH_DorsAttn_PrCv_1 | 0.0035977 | 0.95437 | -0.036905 | 0.76213 |
| 7Networks_LH_SalVentAttn_ParOper_1 | 0.071814 | 0.27158 | -0.14859 | 0.18658 |
| 7Networks_LH_SalVentAttn_ParOper_2 | 0.079167 | 0.2141 | -0.033511 | 0.7435 |
| 7Networks_LH_SalVentAttn_ParOper_3 | 0.013891 | 0.82462 | -0.029126 | 0.81091 |
| 7Networks_LH_SalVentAttn_FrOper_1 | 0.09765 | 0.17208 | -0.1953 | 0.023527 |
| 7Networks_LH_SalVentAttn_FrOper_2 | 0.0014631 | 0.98209 | 0.007139 | 0.93486 |
| 7Networks_LH_SalVentAttn_FrOper_3 | -0.068021 | 0.29949 | 0.026769 | 0.77385 |
| 7Networks_LH_SalVentAttn_FrOper_4 | 0.0035866 | 0.95645 | -0.0079319 | 0.94357 |
| 7Networks_LH_SalVentAttn_PFCl_1 | -0.11949 | 0.064757 | 0.018053 | 0.8932 |
| 7Networks_LH_SalVentAttn_Med_1 | -0.024665 | 0.71406 | 0.019348 | 0.8334 |
| 7Networks_LH_SalVentAttn_Med_2 | 0.011114 | 0.86547 | 0.08836 | 0.3857 |
| 7Networks_LH_SalVentAttn_Med_3 | -0.015097 | 0.81606 | -0.081874 | 0.50255 |
| 7Networks_LH_Limbic_OFC_1 | -0.017751 | 0.78995 | -0.050252 | 0.55391 |
| 7Networks_LH_Limbic_OFC_2 | 0.18932 | 0.0058875 | -0.19541 | 0.027387 |
| 7Networks_LH_Limbic_TempPole_1 | 0.097989 | 0.14509 | -0.20497 | 0.030325 |
| 7Networks_LH_Limbic_TempPole_2 | 0.074895 | 0.23528 | -0.18682 | 0.063945 |
| 7Networks_LH_Limbic_TempPole_3 | -0.053656 | 0.40186 | 0.069135 | 0.51158 |
| 7Networks_LH_Limbic_TempPole_4 | 0.05535 | 0.4238 | -0.13903 | 0.10972 |
| 7Networks_LH_Cont_Par_1 | 0.008729 | 0.88921 | -0.072242 | 0.61615 |
| 7Networks_LH_Cont_Par_2 | -0.047747 | 0.42861 | 0.14609 | 0.30945 |
| 7Networks_LH_Cont_Par_3 | -0.004409 | 0.94062 | -0.011064 | 0.94129 |
| 7Networks_LH_Cont_Temp_1 | -0.083342 | 0.17873 | 0.014923 | 0.899 |
| 7Networks_LH_Cont_PFCl_1 | -0.1039 | 0.11608 | 0.042604 | 0.67411 |
| 7Networks_LH_Cont_PFCl_2 | -0.017436 | 0.79264 | 0.032391 | 0.75479 |
| 7Networks_LH_Cont_PFCl_3 | -0.062233 | 0.35189 | 0.024705 | 0.83061 |
| 7Networks_LH_Cont_PFCl_4 | 0.01229 | 0.85187 | -0.012793 | 0.90749 |
| 7Networks_LH_Cont_PFCl_5 | 0.063455 | 0.30657 | -0.056869 | 0.62639 |
| 7Networks_LH_Cont_PFCl_6 | -0.060547 | 0.31277 | 0.032771 | 0.81504 |
| 7Networks_LH_Cont_pCun_1 | 0.0048466 | 0.9431 | 0.023586 | 0.81339 |
| 7Networks_LH_Cont_Cing_1 | -0.030235 | 0.65656 | -0.04769 | 0.6306 |
| 7Networks_LH_Cont_Cing_2 | 0.035651 | 0.5708 | -0.042436 | 0.72246 |
| 7Networks_LH_Default_Temp_1 | 0.092659 | 0.19391 | -0.1279 | 0.13555 |
| 7Networks_LH_Default_Temp_2 | 0.004595 | 0.94153 | -0.10917 | 0.3546 |
| 7Networks_LH_Default_Temp_3 | -0.018208 | 0.7721 | 0.043273 | 0.68642 |
| 7Networks_LH_Default_Temp_4 | -0.023711 | 0.72058 | -0.057521 | 0.56554 |
| 7Networks_LH_Default_Temp_5 | 0.095485 | 0.12231 | -0.14375 | 0.2243 |
| 7Networks_LH_Default_Temp_6 | 0.031445 | 0.62822 | -0.11976 | 0.30948 |
| 7Networks_LH_Default_Temp_7 | 0.022495 | 0.72121 | -0.11305 | 0.38409 |
| 7Networks_LH_Default_Temp_8 | 0.058394 | 0.32111 | -0.25059 | 0.14946 |
| 7Networks_LH_Default_Temp_9 | 0.032981 | 0.59647 | -0.13412 | 0.24915 |
| 7Networks_LH_Default_PFC_1 | -0.02378 | 0.69774 | 0.043754 | 0.65131 |
| 7Networks_LH_Default_PFC_2 | 0.04749 | 0.47728 | -0.065618 | 0.43886 |
| 7Networks_LH_Default_PFC_3 | 0.057791 | 0.36744 | 0.071083 | 0.48869 |
| 7Networks_LH_Default_PFC_4 | -0.045784 | 0.50029 | -0.0077291 | 0.93817 |
| 7Networks_LH_Default_PFC_5 | -0.0062264 | 0.92132 | 0.053354 | 0.62672 |
| 7Networks_LH_Default_PFC_6 | 0.014113 | 0.83833 | -0.082463 | 0.34927 |
| 7Networks_LH_Default_PFC_7 | 0.011132 | 0.86912 | -0.049814 | 0.65069 |
| 7Networks_LH_Default_PFC_8 | -0.02357 | 0.71975 | -0.092476 | 0.3803 |
| 7Networks_LH_Default_PFC_9 | -0.0081536 | 0.90203 | 0.035083 | 0.75891 |
| 7Networks_LH_Default_PFC_10 | 0.013141 | 0.84168 | -0.059935 | 0.55688 |
| 7Networks_LH_Default_PFC_11 | -0.016062 | 0.79576 | -0.013816 | 0.90462 |
| 7Networks_LH_Default_PFC_12 | 0.045178 | 0.4712 | 0.0088629 | 0.94354 |
| 7Networks_LH_Default_PFC_13 | 0.028915 | 0.66058 | -0.0022993 | 0.98551 |
| 7Networks_LH_Default_PCC_1 | 0.10237 | 0.15799 | -0.022006 | 0.77895 |
| 7Networks_LH_Default_PCC_2 | -0.017781 | 0.79004 | -0.034372 | 0.7348 |
| 7Networks_LH_Default_PCC_3 | -0.0027765 | 0.96521 | 0.022648 | 0.84251 |
| 7Networks_LH_Default_PCC_4 | -0.0089398 | 0.89109 | -0.072829 | 0.50738 |
| 7Networks_LH_Default_PHC_1 | 0.16815 | 0.010472 | -0.25039 | 0.0037036 |
| 7Networks_RH_Vis_1 | 0.031181 | 0.641 | -0.10455 | 0.25998 |
| 7Networks_RH_Vis_2 | 0.06328 | 0.34218 | -0.16978 | 0.077809 |
| 7Networks_RH_Vis_3 | 0.045826 | 0.48578 | -0.13057 | 0.22251 |
| 7Networks_RH_Vis_4 | -0.0067432 | 0.91828 | -0.061881 | 0.51314 |
| 7Networks_RH_Vis_5 | -0.028082 | 0.65014 | -0.057934 | 0.66751 |
| 7Networks_RH_Vis_6 | 0.014374 | 0.84622 | 0.0125 | 0.88529 |
| 7Networks_RH_Vis_7 | -0.15288 | 0.028932 | 0.11277 | 0.18787 |
| 7Networks_RH_Vis_8 | -0.057867 | 0.38035 | -0.025397 | 0.81982 |
| 7Networks_RH_Vis_9 | 0.031235 | 0.68045 | 0.028952 | 0.6972 |
| 7Networks_RH_Vis_10 | -0.10526 | 0.16869 | 0.11823 | 0.10687 |
| 7Networks_RH_Vis_11 | -0.0069755 | 0.91029 | 0.085652 | 0.51258 |
| 7Networks_RH_Vis_12 | 0.035928 | 0.59085 | -0.033226 | 0.72043 |
| 7Networks_RH_Vis_13 | -0.01877 | 0.78155 | -0.022626 | 0.81487 |
| 7Networks_RH_Vis_14 | 0.1022 | 0.12038 | -0.090728 | 0.40136 |
| 7Networks_RH_Vis_15 | 0.10484 | 0.089399 | -0.04821 | 0.6947 |
| 7Networks_RH_SomMot_1 | -0.041485 | 0.54644 | 0.041369 | 0.63227 |
| 7Networks_RH_SomMot_2 | -0.038198 | 0.58159 | 0.083708 | 0.36405 |
| 7Networks_RH_SomMot_3 | 0.014909 | 0.82583 | 0.038058 | 0.7028 |
| 7Networks_RH_SomMot_4 | -0.078016 | 0.22716 | 0.14228 | 0.15704 |
| 7Networks_RH_SomMot_5 | 0.0032889 | 0.95742 | 0.064585 | 0.61067 |
| 7Networks_RH_SomMot_6 | -0.049669 | 0.46866 | 0.13193 | 0.20177 |
| 7Networks_RH_SomMot_7 | -0.027664 | 0.66582 | -0.056025 | 0.59597 |
| 7Networks_RH_SomMot_8 | -0.083805 | 0.18454 | 0.030015 | 0.79694 |
| 7Networks_RH_SomMot_9 | -0.055239 | 0.3552 | 0.15189 | 0.30739 |
| 7Networks_RH_SomMot_10 | 0.019844 | 0.74448 | -0.18111 | 0.1283 |
| 7Networks_RH_SomMot_11 | 0.051185 | 0.44713 | -0.11939 | 0.20128 |
| 7Networks_RH_SomMot_12 | -0.057938 | 0.35055 | 0.0409 | 0.71187 |
| 7Networks_RH_SomMot_13 | 0.068871 | 0.28441 | -0.13127 | 0.27954 |
| 7Networks_RH_SomMot_14 | 0.012105 | 0.8514 | -0.11227 | 0.26429 |
| 7Networks_RH_SomMot_15 | -0.04407 | 0.45956 | 0.083005 | 0.57183 |
| 7Networks_RH_SomMot_16 | -0.035838 | 0.56809 | 0.039386 | 0.75381 |
| 7Networks_RH_SomMot_17 | 0.039974 | 0.50394 | -0.10958 | 0.50604 |
| 7Networks_RH_SomMot_18 | -0.016044 | 0.80473 | -0.025691 | 0.79445 |
| 7Networks_RH_SomMot_19 | 0.015527 | 0.8063 | -0.055803 | 0.58145 |
| 7Networks_RH_DorsAttn_Post_1 | 0.075505 | 0.24869 | -0.10044 | 0.3051 |
| 7Networks_RH_DorsAttn_Post_2 | -0.023709 | 0.70655 | -0.13365 | 0.27184 |
| 7Networks_RH_DorsAttn_Post_3 | 0.0075621 | 0.90358 | 0.098093 | 0.35173 |
| 7Networks_RH_DorsAttn_Post_4 | -0.0080338 | 0.89225 | -0.08985 | 0.50777 |
| 7Networks_RH_DorsAttn_Post_5 | 0.068128 | 0.2674 | -0.10539 | 0.40286 |
| 7Networks_RH_DorsAttn_Post_6 | -0.035719 | 0.56121 | 0.0066304 | 0.96098 |
| 7Networks_RH_DorsAttn_Post_7 | -0.010375 | 0.86903 | 0.0075825 | 0.95844 |
| 7Networks_RH_DorsAttn_Post_8 | 0.040775 | 0.50288 | -0.11758 | 0.4016 |
| 7Networks_RH_DorsAttn_Post_9 | 0.11749 | 0.071095 | -0.14584 | 0.17077 |
| 7Networks_RH_DorsAttn_Post_10 | 0.056069 | 0.37418 | -0.13053 | 0.24755 |
| 7Networks_RH_DorsAttn_FEF_1 | -0.012354 | 0.84088 | -0.091378 | 0.54099 |
| 7Networks_RH_DorsAttn_FEF_2 | -0.047144 | 0.49048 | 0.076638 | 0.41348 |
| 7Networks_RH_DorsAttn_PrCv_1 | -0.010068 | 0.87032 | -0.021831 | 0.85298 |
| 7Networks_RH_SalVentAttn_TempOccPar_1 | 0.09417 | 0.13107 | -0.21978 | 0.06921 |
| 7Networks_RH_SalVentAttn_TempOccPar_2 | 0.059292 | 0.32646 | -0.17021 | 0.22894 |
| 7Networks_RH_SalVentAttn_TempOccPar_3 | 0.016346 | 0.7996 | 0.056614 | 0.60214 |
| 7Networks_RH_SalVentAttn_PrC_1 | -0.029995 | 0.6231 | -0.071747 | 0.57159 |
| 7Networks_RH_SalVentAttn_FrOper_1 | -0.022115 | 0.74488 | -0.06062 | 0.49814 |
| 7Networks_RH_SalVentAttn_FrOper_2 | 0.034346 | 0.60702 | -0.098767 | 0.27785 |
| 7Networks_RH_SalVentAttn_FrOper_3 | -0.0008578 | 0.98912 | 0.0042609 | 0.96736 |
| 7Networks_RH_SalVentAttn_FrOper_4 | -0.0088675 | 0.89243 | -0.080989 | 0.4206 |
| 7Networks_RH_SalVentAttn_Med_1 | -0.015596 | 0.82146 | -0.056874 | 0.52861 |
| 7Networks_RH_SalVentAttn_Med_2 | -0.041699 | 0.52841 | -0.011594 | 0.90178 |
| 7Networks_RH_SalVentAttn_Med_3 | -0.031783 | 0.63095 | 0.022127 | 0.8223 |
| 7Networks_RH_Limbic_OFC_1 | -0.0083306 | 0.90174 | -0.058824 | 0.52349 |
| 7Networks_RH_Limbic_OFC_2 | 0.03839 | 0.54743 | -0.13334 | 0.15281 |
| 7Networks_RH_Limbic_OFC_3 | -0.10251 | 0.13164 | 0.080403 | 0.45533 |
| 7Networks_RH_Limbic_TempPole_1 | -0.021313 | 0.7395 | -0.0072543 | 0.93934 |
| 7Networks_RH_Limbic_TempPole_2 | -0.015557 | 0.80758 | -0.023384 | 0.81358 |
| 7Networks_RH_Limbic_TempPole_3 | 0.041459 | 0.5248 | -0.087159 | 0.39457 |
| 7Networks_RH_Cont_Par_1 | 0.0681 | 0.26616 | -0.18694 | 0.15143 |
| 7Networks_RH_Cont_Par_2 | -0.029663 | 0.63366 | -0.11199 | 0.46378 |
| 7Networks_RH_Cont_Par_3 | 0.047716 | 0.41949 | -0.058868 | 0.68381 |
| 7Networks_RH_Cont_Temp_1 | 0.14606 | 0.023159 | -0.37118 | 0.0026228 |
| 7Networks_RH_Cont_PFCv_1 | 0.015077 | 0.80172 | -0.15095 | 0.20495 |
| 7Networks_RH_Cont_PFCl_1 | 0.043381 | 0.50544 | -0.086328 | 0.34179 |
| 7Networks_RH_Cont_PFCl_2 | -0.10091 | 0.11833 | 0.077895 | 0.52314 |
| 7Networks_RH_Cont_PFCl_3 | -0.069669 | 0.27927 | 0.10508 | 0.33009 |
| 7Networks_RH_Cont_PFCl_4 | -0.048676 | 0.45772 | -0.025079 | 0.81353 |
| 7Networks_RH_Cont_PFCl_5 | 0.030024 | 0.6351 | -0.016899 | 0.89113 |
| 7Networks_RH_Cont_PFCl_6 | -0.00223 | 0.97093 | 0.0012626 | 0.99141 |
| 7Networks_RH_Cont_PFCl_7 | -0.085125 | 0.1616 | 0.0428 | 0.69113 |
| 7Networks_RH_Cont_pCun_1 | 0.052369 | 0.43504 | -0.14649 | 0.17287 |
| 7Networks_RH_Cont_PFCmp_1 | -0.028947 | 0.66319 | 0.02347 | 0.8161 |
| 7Networks_RH_Cont_PFCmp_2 | -0.012673 | 0.84535 | 0.085949 | 0.43436 |
| 7Networks_RH_Cont_PFCmp_3 | -0.0016197 | 0.97954 | -0.048473 | 0.59229 |
| 7Networks_RH_Cont_PFCmp_4 | 0.090298 | 0.17349 | -0.17784 | 0.082089 |
| 7Networks_RH_Default_Par_1 | -0.067312 | 0.28497 | 0.0093088 | 0.9376 |
| 7Networks_RH_Default_Par_2 | 0.10468 | 0.076702 | -0.5114 | 0.0006666 |
| 7Networks_RH_Default_Par_3 | 0.1177 | 0.046936 | -0.56102 | 0.0009986 |
| 7Networks_RH_Default_Temp_1 | 0.14165 | 0.029395 | -0.12018 | 0.20268 |
| 7Networks_RH_Default_Temp_2 | -0.03956 | 0.53082 | -0.014094 | 0.89269 |
| 7Networks_RH_Default_Temp_3 | 0.023167 | 0.7056 | -0.05196 | 0.61375 |
| 7Networks_RH_Default_Temp_4 | -0.05961 | 0.34187 | -0.041998 | 0.71289 |
| 7Networks_RH_Default_Temp_5 | 0.010676 | 0.87061 | -0.088027 | 0.38781 |
| 7Networks_RH_Default_PFCv_1 | -0.057084 | 0.35173 | 0.058087 | 0.60862 |
| 7Networks_RH_Default_PFCm_1 | -0.017603 | 0.80447 | -0.051424 | 0.57324 |
| 7Networks_RH_Default_PFCm_2 | -0.046314 | 0.50379 | -0.0050646 | 0.9596 |
| 7Networks_RH_Default_PFCm_3 | -0.02174 | 0.73254 | 0.09338 | 0.56732 |
| 7Networks_RH_Default_PFCm_4 | 0.039269 | 0.57399 | -0.082726 | 0.37699 |
| 7Networks_RH_Default_PFCm_5 | -0.061473 | 0.3366 | 0.074862 | 0.5032 |
| 7Networks_RH_Default_PFCm_6 | -0.089304 | 0.14861 | 0.071238 | 0.5379 |
| 7Networks_RH_Default_PFCm_7 | 0.00068 | 0.99126 | -0.030805 | 0.80798 |
| 7Networks_RH_Default_PCC_1 | -0.0039496 | 0.95631 | -0.060424 | 0.46419 |
| 7Networks_RH_Default_PCC_2 | -0.0049658 | 0.94015 | -0.0083853 | 0.93964 |
| 7Networks_RH_Default_PCC_3 | -0.025645 | 0.69174 | 0.035839 | 0.77161 |

**Supplementary Table 13. Genetic and environmental correlation between Conscientiousness and surface area in HCP.** Genetic correlation between Conscientiousness and local surface area calculated using solar 8.4.0. Regions are named according to the Schaefer 200 atlas, based on 7-networks. Here we report environmental correlation (ρ_e_) and genetic correlation (ρ_g_) and the associated p-values.

| ROIS | ρ_e_ | p | ρ_g_ | p |
| --- | --- | --- | --- | --- |
| 7Networks_LH_Vis_1 | 0.1093 | 0.10561 | -0.016092 | 0.8522 |
| 7Networks_LH_Vis_2 | 0.0084063 | 0.9002 | -0.035603 | 0.70597 |
| 7Networks_LH_Vis_3 | 0.085706 | 0.16193 | -0.10192 | 0.34625 |
| 7Networks_LH_Vis_4 | -0.040758 | 0.52746 | 0.084891 | 0.42456 |
| 7Networks_LH_Vis_5 | 0.020265 | 0.74133 | 0.062845 | 0.59621 |
| 7Networks_LH_Vis_6 | 0.0009481 | 0.98906 | -0.054269 | 0.52433 |
| 7Networks_LH_Vis_7 | 0.12933 | 0.053191 | -0.11008 | 0.19154 |
| 7Networks_LH_Vis_8 | -0.016752 | 0.78721 | -0.037657 | 0.73012 |
| 7Networks_LH_Vis_9 | 0.016367 | 0.79329 | -0.026143 | 0.78705 |
| 7Networks_LH_Vis_10 | 0.098877 | 0.17508 | -0.08068 | 0.22494 |
| 7Networks_LH_Vis_11 | -0.061261 | 0.33291 | 0.002959 | 0.97583 |
| 7Networks_LH_Vis_12 | 0.15779 | 0.016339 | -0.14408 | 0.091392 |
| 7Networks_LH_Vis_13 | 0.12739 | 0.060533 | -0.087209 | 0.28706 |
| 7Networks_LH_Vis_14 | -0.02148 | 0.73998 | 0.044487 | 0.65184 |
| 7Networks_LH_SomMot_1 | -0.018522 | 0.77061 | 0.031272 | 0.74273 |
| 7Networks_LH_SomMot_2 | 0.012855 | 0.83726 | 0.081416 | 0.36395 |
| 7Networks_LH_SomMot_3 | 0.012215 | 0.84289 | 0.023222 | 0.81804 |
| 7Networks_LH_SomMot_4 | 0.068049 | 0.25858 | -0.13514 | 0.22858 |
| 7Networks_LH_SomMot_5 | -0.033977 | 0.59439 | 0.056204 | 0.58641 |
| 7Networks_LH_SomMot_6 | 0.084184 | 0.17167 | -0.12874 | 0.17311 |
| 7Networks_LH_SomMot_7 | 0.0001215 | 0.99774 | -0.055763 | 0.65586 |
| 7Networks_LH_SomMot_8 | -0.0036207 | 0.95313 | 0.068816 | 0.46164 |
| 7Networks_LH_SomMot_9 | -0.046878 | 0.41721 | 0.066015 | 0.62785 |
| 7Networks_LH_SomMot_10 | -0.063161 | 0.28825 | 0.0006613 | 0.99471 |
| 7Networks_LH_SomMot_11 | 0.061221 | 0.33325 | -0.0004272 | 0.99643 |
| 7Networks_LH_SomMot_12 | -0.090813 | 0.13554 | 0.10973 | 0.23529 |
| 7Networks_LH_SomMot_13 | 0.0026687 | 0.96622 | 0.012946 | 0.90193 |
| 7Networks_LH_SomMot_14 | 0.040597 | 0.49033 | -0.074672 | 0.52257 |
| 7Networks_LH_SomMot_15 | -0.033433 | 0.58927 | 0.13466 | 0.18264 |
| 7Networks_LH_SomMot_16 | -0.043127 | 0.47575 | 0.11645 | 0.25322 |
| 7Networks_LH_DorsAttn_Post_1 | -0.085428 | 0.16338 | 0.049967 | 0.65117 |
| 7Networks_LH_DorsAttn_Post_2 | -0.024441 | 0.69447 | 0.053607 | 0.61406 |
| 7Networks_LH_DorsAttn_Post_3 | 0.0018704 | 0.97732 | 0.018872 | 0.85075 |
| 7Networks_LH_DorsAttn_Post_4 | -0.02626 | 0.67334 | 0.016366 | 0.88181 |
| 7Networks_LH_DorsAttn_Post_5 | 0.025468 | 0.65996 | -0.033149 | 0.80769 |
| 7Networks_LH_DorsAttn_Post_6 | 0.070941 | 0.21043 | -0.18215 | 0.19477 |
| 7Networks_LH_DorsAttn_Post_7 | 0.086395 | 0.14899 | 0.014865 | 0.89186 |
| 7Networks_LH_DorsAttn_Post_8 | 0.09592 | 0.10529 | 0.078399 | 0.49349 |
| 7Networks_LH_DorsAttn_Post_9 | 0.035143 | 0.57117 | 0.012094 | 0.90404 |
| 7Networks_LH_DorsAttn_Post_10 | -0.022343 | 0.7132 | 0.059688 | 0.58419 |
| 7Networks_LH_DorsAttn_FEF_1 | -0.027559 | 0.63451 | 0.035892 | 0.76373 |
| 7Networks_LH_DorsAttn_FEF_2 | 0.034026 | 0.55494 | 0.039533 | 0.75545 |
| 7Networks_LH_DorsAttn_PrCv_1 | 0.016941 | 0.7818 | 0.061479 | 0.60065 |
| 7Networks_LH_SalVentAttn_ParOper_1 | -0.0090056 | 0.88498 | 0.1105 | 0.30967 |
| 7Networks_LH_SalVentAttn_ParOper_2 | 0.027733 | 0.65119 | 0.013772 | 0.88914 |
| 7Networks_LH_SalVentAttn_ParOper_3 | -0.10104 | 0.094961 | 0.32362 | 0.0054801 |
| 7Networks_LH_SalVentAttn_FrOper_1 | 0.074427 | 0.26698 | -0.0044661 | 0.95753 |
| 7Networks_LH_SalVentAttn_FrOper_2 | -0.054688 | 0.38618 | 0.13908 | 0.094826 |
| 7Networks_LH_SalVentAttn_FrOper_3 | -0.0299 | 0.63582 | 0.039253 | 0.66179 |
| 7Networks_LH_SalVentAttn_FrOper_4 | 0.014705 | 0.81642 | -0.0064225 | 0.95205 |
| 7Networks_LH_SalVentAttn_PFCl_1 | -0.094054 | 0.13492 | 0.16063 | 0.21071 |
| 7Networks_LH_SalVentAttn_Med_1 | 0.035776 | 0.58127 | 0.14966 | 0.090079 |
| 7Networks_LH_SalVentAttn_Med_2 | -0.035726 | 0.57355 | 0.079034 | 0.41695 |
| 7Networks_LH_SalVentAttn_Med_3 | -0.049529 | 0.43319 | 0.10699 | 0.35882 |
| 7Networks_LH_Limbic_OFC_1 | -0.14128 | 0.026736 | 0.14084 | 0.087946 |
| 7Networks_LH_Limbic_OFC_2 | 0.10277 | 0.11915 | -0.016583 | 0.84691 |
| 7Networks_LH_Limbic_TempPole_1 | 0.085961 | 0.18888 | -0.070707 | 0.4274 |
| 7Networks_LH_Limbic_TempPole_2 | 0.016533 | 0.78761 | -0.07948 | 0.41325 |
| 7Networks_LH_Limbic_TempPole_3 | -0.0091323 | 0.88236 | 0.031584 | 0.75565 |
| 7Networks_LH_Limbic_TempPole_4 | 0.017948 | 0.78749 | -0.037756 | 0.6504 |
| 7Networks_LH_Cont_Par_1 | -0.024384 | 0.6881 | 0.14288 | 0.29865 |
| 7Networks_LH_Cont_Par_2 | 0.041364 | 0.4805 | 0.016763 | 0.90413 |
| 7Networks_LH_Cont_Par_3 | 0.051297 | 0.37629 | -0.048217 | 0.73853 |
| 7Networks_LH_Cont_Temp_1 | -0.010207 | 0.8659 | -0.0049548 | 0.9652 |
| 7Networks_LH_Cont_PFCl_1 | -0.05468 | 0.39411 | 0.051898 | 0.59872 |
| 7Networks_LH_Cont_PFCl_2 | 0.0065631 | 0.91828 | 0.0026191 | 0.97871 |
| 7Networks_LH_Cont_PFCl_3 | -0.055455 | 0.3889 | 0.11684 | 0.28739 |
| 7Networks_LH_Cont_PFCl_4 | -0.021341 | 0.73401 | 0.065126 | 0.53158 |
| 7Networks_LH_Cont_PFCl_5 | 0.053859 | 0.37675 | 0.0031175 | 0.97798 |
| 7Networks_LH_Cont_PFCl_6 | -0.03988 | 0.4948 | 0.092247 | 0.49422 |
| 7Networks_LH_Cont_pCun_1 | 0.084743 | 0.19428 | -0.041693 | 0.66754 |
| 7Networks_LH_Cont_Cing_1 | -0.034874 | 0.59232 | 0.060474 | 0.52208 |
| 7Networks_LH_Cont_Cing_2 | 0.0025995 | 0.96599 | -0.035003 | 0.76189 |
| 7Networks_LH_Default_Temp_1 | 0.03383 | 0.62011 | -0.070568 | 0.38887 |
| 7Networks_LH_Default_Temp_2 | -0.028979 | 0.63365 | -0.012172 | 0.91369 |
| 7Networks_LH_Default_Temp_3 | -0.0025398 | 0.96599 | 0.036051 | 0.72578 |
| 7Networks_LH_Default_Temp_4 | 0.030484 | 0.63342 | -0.037139 | 0.69727 |
| 7Networks_LH_Default_Temp_5 | 0.004817 | 0.93628 | 0.07998 | 0.47628 |
| 7Networks_LH_Default_Temp_6 | 0.019222 | 0.75951 | -0.010894 | 0.92326 |
| 7Networks_LH_Default_Temp_7 | 0.023036 | 0.70625 | 0.069793 | 0.57653 |
| 7Networks_LH_Default_Temp_8 | -0.024583 | 0.67026 | 0.087127 | 0.5989 |
| 7Networks_LH_Default_Temp_9 | 0.037584 | 0.53417 | -0.051857 | 0.64358 |
| 7Networks_LH_Default_PFC_1 | -0.050764 | 0.39218 | 0.16336 | 0.08222 |
| 7Networks_LH_Default_PFC_2 | -0.065891 | 0.3063 | 0.064534 | 0.42944 |
| 7Networks_LH_Default_PFC_3 | -0.0050335 | 0.93516 | 0.16672 | 0.092129 |
| 7Networks_LH_Default_PFC_4 | 0.035853 | 0.57789 | 0.013447 | 0.88701 |
| 7Networks_LH_Default_PFC_5 | -0.012885 | 0.83263 | 0.10294 | 0.32721 |
| 7Networks_LH_Default_PFC_6 | 0.054812 | 0.40726 | -0.069595 | 0.41061 |
| 7Networks_LH_Default_PFC_7 | 0.0023226 | 0.97153 | 0.028714 | 0.78343 |
| 7Networks_LH_Default_PFC_8 | 0.074281 | 0.2443 | -0.027321 | 0.78614 |
| 7Networks_LH_Default_PFC_9 | -0.047938 | 0.45296 | 0.16874 | 0.12008 |
| 7Networks_LH_Default_PFC_10 | -0.0077895 | 0.90178 | 0.072627 | 0.45779 |
| 7Networks_LH_Default_PFC_11 | -0.024897 | 0.68007 | 0.0018847 | 0.98623 |
| 7Networks_LH_Default_PFC_12 | 0.033716 | 0.5796 | 0.078857 | 0.51015 |
| 7Networks_LH_Default_PFC_13 | 0.086057 | 0.17575 | 0.019414 | 0.8748 |
| 7Networks_LH_Default_PCC_1 | 0.20657 | 0.0028714 | -0.15835 | 0.034174 |
| 7Networks_LH_Default_PCC_2 | -0.012907 | 0.84084 | 0.090499 | 0.35167 |
| 7Networks_LH_Default_PCC_3 | 0.0069257 | 0.9103 | 0.081167 | 0.45496 |
| 7Networks_LH_Default_PCC_4 | -0.077284 | 0.21701 | 0.19198 | 0.069655 |
| 7Networks_LH_Default_PHC_1 | 0.11265 | 0.074566 | -0.091898 | 0.27645 |
| 7Networks_RH_Vis_1 | 0.021503 | 0.7387 | -0.034578 | 0.69658 |
| 7Networks_RH_Vis_2 | 0.034294 | 0.59423 | 0.0007791 | 0.99304 |
| 7Networks_RH_Vis_3 | 0.090029 | 0.15622 | -0.020337 | 0.84218 |
| 7Networks_RH_Vis_4 | -0.057066 | 0.36786 | 0.089874 | 0.32521 |
| 7Networks_RH_Vis_5 | -0.014919 | 0.80382 | 0.11595 | 0.36878 |
| 7Networks_RH_Vis_6 | 0.0020902 | 0.97567 | 0.011134 | 0.89351 |
| 7Networks_RH_Vis_7 | -0.035799 | 0.59447 | 0.01481 | 0.85862 |
| 7Networks_RH_Vis_8 | 0.038699 | 0.5436 | -0.0030115 | 0.97724 |
| 7Networks_RH_Vis_9 | 0.048164 | 0.50472 | -0.000327 | 0.99626 |
| 7Networks_RH_Vis_10 | 0.040761 | 0.57803 | -0.021148 | 0.76351 |
| 7Networks_RH_Vis_11 | -0.021678 | 0.71969 | 0.033256 | 0.79161 |
| 7Networks_RH_Vis_12 | -0.035286 | 0.58406 | 0.031961 | 0.72088 |
| 7Networks_RH_Vis_13 | 0.084374 | 0.19413 | -0.037074 | 0.68882 |
| 7Networks_RH_Vis_14 | 0.11719 | 0.063741 | -0.061283 | 0.55707 |
| 7Networks_RH_Vis_15 | 0.098019 | 0.10533 | -0.16052 | 0.17226 |
| 7Networks_RH_SomMot_1 | -0.055554 | 0.40048 | 0.054038 | 0.51454 |
| 7Networks_RH_SomMot_2 | 0.082732 | 0.21226 | 0.042343 | 0.62663 |
| 7Networks_RH_SomMot_3 | 0.0083719 | 0.89775 | -0.051252 | 0.58992 |
| 7Networks_RH_SomMot_4 | -0.013503 | 0.8296 | 0.062385 | 0.50901 |
| 7Networks_RH_SomMot_5 | 0.069692 | 0.24378 | -0.039754 | 0.74504 |
| 7Networks_RH_SomMot_6 | 0.079356 | 0.22862 | -0.053262 | 0.5887 |
| 7Networks_RH_SomMot_7 | 0.11756 | 0.056148 | -0.13715 | 0.18888 |
| 7Networks_RH_SomMot_8 | 0.011715 | 0.84731 | -0.010189 | 0.92772 |
| 7Networks_RH_SomMot_9 | 0.021465 | 0.71388 | -0.072498 | 0.61267 |
| 7Networks_RH_SomMot_10 | 0.089915 | 0.1288 | -0.22346 | 0.050831 |
| 7Networks_RH_SomMot_11 | -0.051193 | 0.43189 | 0.17091 | 0.056316 |
| 7Networks_RH_SomMot_12 | -0.018689 | 0.7572 | -0.073138 | 0.49126 |
| 7Networks_RH_SomMot_13 | 0.016669 | 0.79065 | 0.081268 | 0.47622 |
| 7Networks_RH_SomMot_14 | 0.038095 | 0.54167 | -0.042431 | 0.66181 |
| 7Networks_RH_SomMot_15 | 0.0078332 | 0.89395 | 0.14256 | 0.31333 |
| 7Networks_RH_SomMot_16 | 0.050967 | 0.40262 | 0.095988 | 0.42639 |
| 7Networks_RH_SomMot_17 | 0.051619 | 0.37769 | 0.036699 | 0.81647 |
| 7Networks_RH_SomMot_18 | -0.035473 | 0.57208 | 0.035146 | 0.71157 |
| 7Networks_RH_SomMot_19 | 0.053942 | 0.37973 | -0.057541 | 0.55398 |
| 7Networks_RH_DorsAttn_Post_1 | -0.15368 | 0.014489 | 0.10014 | 0.28697 |
| 7Networks_RH_DorsAttn_Post_2 | -0.034664 | 0.5705 | 0.021029 | 0.8563 |
| 7Networks_RH_DorsAttn_Post_3 | 0.092827 | 0.12488 | -0.067071 | 0.50917 |
| 7Networks_RH_DorsAttn_Post_4 | -0.015706 | 0.78665 | -0.086937 | 0.50477 |
| 7Networks_RH_DorsAttn_Post_5 | -0.033223 | 0.57828 | 0.027414 | 0.8219 |
| 7Networks_RH_DorsAttn_Post_6 | 0.09767 | 0.10287 | -0.068638 | 0.59899 |
| 7Networks_RH_DorsAttn_Post_7 | -0.061476 | 0.31276 | 0.12273 | 0.37554 |
| 7Networks_RH_DorsAttn_Post_8 | 0.053852 | 0.36318 | 0.069709 | 0.60477 |
| 7Networks_RH_DorsAttn_Post_9 | 0.013939 | 0.8245 | 0.080826 | 0.43085 |
| 7Networks_RH_DorsAttn_Post_10 | -0.038805 | 0.52845 | 0.13308 | 0.21539 |
| 7Networks_RH_DorsAttn_FEF_1 | -0.018251 | 0.7587 | 0.054043 | 0.70727 |
| 7Networks_RH_DorsAttn_FEF_2 | 0.0015014 | 0.97975 | 0.14158 | 0.11779 |
| 7Networks_RH_DorsAttn_PrCv_1 | -0.13625 | 0.022374 | 0.18673 | 0.10342 |
| 7Networks_RH_SalVentAttn_TempOccPar_1 | 0.046398 | 0.44667 | -0.051738 | 0.65477 |
| 7Networks_RH_SalVentAttn_TempOccPar_2 | 0.13942 | 0.016628 | -0.12666 | 0.35512 |
| 7Networks_RH_SalVentAttn_TempOccPar_3 | 0.041416 | 0.50679 | -0.010553 | 0.91935 |
| 7Networks_RH_SalVentAttn_PrC_1 | 0.099871 | 0.093784 | -0.070977 | 0.55837 |
| 7Networks_RH_SalVentAttn_FrOper_1 | 0.065144 | 0.31756 | -0.067495 | 0.43026 |
| 7Networks_RH_SalVentAttn_FrOper_2 | -0.054206 | 0.39871 | 0.042588 | 0.62649 |
| 7Networks_RH_SalVentAttn_FrOper_3 | -0.023895 | 0.69598 | 0.16598 | 0.09601 |
| 7Networks_RH_SalVentAttn_FrOper_4 | 0.068727 | 0.27661 | 0.01579 | 0.86996 |
| 7Networks_RH_SalVentAttn_Med_1 | -0.027186 | 0.68188 | 0.058578 | 0.49903 |
| 7Networks_RH_SalVentAttn_Med_2 | 0.007384 | 0.90788 | -0.068603 | 0.44666 |
| 7Networks_RH_SalVentAttn_Med_3 | 0.021754 | 0.73246 | 0.15116 | 0.1078 |
| 7Networks_RH_Limbic_OFC_1 | 0.036105 | 0.57877 | -0.016421 | 0.85236 |
| 7Networks_RH_Limbic_OFC_2 | -0.10589 | 0.084664 | 0.075035 | 0.40532 |
| 7Networks_RH_Limbic_OFC_3 | -0.076467 | 0.24919 | 0.21767 | 0.032199 |
| 7Networks_RH_Limbic_TempPole_1 | 0.05501 | 0.37418 | -0.048601 | 0.59797 |
| 7Networks_RH_Limbic_TempPole_2 | 0.029765 | 0.62983 | 0.039198 | 0.68067 |
| 7Networks_RH_Limbic_TempPole_3 | 0.085408 | 0.17371 | -0.029891 | 0.75877 |
| 7Networks_RH_Cont_Par_1 | 0.071634 | 0.23087 | -0.11095 | 0.37827 |
| 7Networks_RH_Cont_Par_2 | -0.015592 | 0.79676 | 0.01979 | 0.89126 |
| 7Networks_RH_Cont_Par_3 | 0.0031196 | 0.95701 | 0.028848 | 0.8369 |
| 7Networks_RH_Cont_Temp_1 | -0.064613 | 0.31268 | 0.098763 | 0.37864 |
| 7Networks_RH_Cont_PFCv_1 | -0.088672 | 0.12959 | 0.11125 | 0.33196 |
| 7Networks_RH_Cont_PFCl_1 | -0.033655 | 0.59148 | 0.135 | 0.12496 |
| 7Networks_RH_Cont_PFCl_2 | 0.053021 | 0.39656 | 0.052858 | 0.6516 |
| 7Networks_RH_Cont_PFCl_3 | -0.043907 | 0.48244 | 0.088681 | 0.38786 |
| 7Networks_RH_Cont_PFCl_4 | -0.12275 | 0.050825 | 0.087198 | 0.39276 |
| 7Networks_RH_Cont_PFCl_5 | 0.091517 | 0.13735 | -0.12805 | 0.2722 |
| 7Networks_RH_Cont_PFCl_6 | 0.063746 | 0.29177 | -0.16194 | 0.18114 |
| 7Networks_RH_Cont_PFCl_7 | -0.039518 | 0.50464 | 0.060581 | 0.55761 |
| 7Networks_RH_Cont_pCun_1 | 0.028446 | 0.66013 | -0.029886 | 0.77348 |
| 7Networks_RH_Cont_PFCmp_1 | -0.0073472 | 0.90864 | 0.075432 | 0.43247 |
| 7Networks_RH_Cont_PFCmp_2 | 0.027413 | 0.66269 | 0.0010647 | 0.99194 |
| 7Networks_RH_Cont_PFCmp_3 | -0.14484 | 0.026066 | 0.16848 | 0.051071 |
| 7Networks_RH_Cont_PFCmp_4 | 0.085439 | 0.17698 | 0.081332 | 0.41424 |
| 7Networks_RH_Default_Par_1 | -0.057328 | 0.34771 | 0.15152 | 0.18387 |
| 7Networks_RH_Default_Par_2 | 0.012934 | 0.82638 | 0.059717 | 0.68175 |
| 7Networks_RH_Default_Par_3 | 0.075107 | 0.20211 | 0.011218 | 0.94287 |
| 7Networks_RH_Default_Temp_1 | 0.081089 | 0.19369 | -0.11096 | 0.22525 |
| 7Networks_RH_Default_Temp_2 | 0.038902 | 0.52451 | 0.036132 | 0.71947 |
| 7Networks_RH_Default_Temp_3 | -0.020647 | 0.72812 | 0.10914 | 0.27062 |
| 7Networks_RH_Default_Temp_4 | 0.048782 | 0.42426 | -0.024114 | 0.82545 |
| 7Networks_RH_Default_Temp_5 | 0.0087204 | 0.89048 | -0.0001928 | 0.99805 |
| 7Networks_RH_Default_PFCv_1 | -0.091311 | 0.12447 | 0.21188 | 0.046406 |
| 7Networks_RH_Default_PFCm_1 | -0.0021662 | 0.97395 | 0.074603 | 0.39298 |
| 7Networks_RH_Default_PFCm_2 | -0.042504 | 0.52273 | 0.077284 | 0.41688 |
| 7Networks_RH_Default_PFCm_3 | -0.094449 | 0.12485 | 0.21254 | 0.16935 |
| 7Networks_RH_Default_PFCm_4 | -0.010466 | 0.87596 | 0.11639 | 0.19579 |
| 7Networks_RH_Default_PFCm_5 | -0.057903 | 0.34928 | 0.27718 | 0.008664 |
| 7Networks_RH_Default_PFCm_6 | -0.035815 | 0.55059 | 0.13978 | 0.2108 |
| 7Networks_RH_Default_PFCm_7 | -0.04455 | 0.45858 | 0.27319 | 0.025216 |
| 7Networks_RH_Default_PCC_1 | -0.020008 | 0.77135 | 0.0085089 | 0.91442 |
| 7Networks_RH_Default_PCC_2 | -0.11392 | 0.073493 | 0.076246 | 0.46894 |
| 7Networks_RH_Default_PCC_3 | -0.11545 | 0.066358 | 0.11143 | 0.33935 |

**Supplementary Table 14. Genetic and environmental correlation between Extraversion and surface area in HCP.** Genetic correlation between Extraversion and local surface area calculated using solar 8.4.0. Regions are named according to the Schaefer 200 atlas, based on 7-networks. Here we report environmental correlation (ρ_e_) and genetic correlation (ρ_g_) and the associated p-values.

| ROIS | ρ_e_ | p | ρ_g_ | p |
| --- | --- | --- | --- | --- |
| 7Networks_LH_Vis_1 | -0.069857 | 0.32596 | -0.055292 | 0.57636 |
| 7Networks_LH_Vis_2 | 0.06263 | 0.371 | -0.084981 | 0.43827 |
| 7Networks_LH_Vis_3 | -0.09104 | 0.15332 | 0.069251 | 0.57585 |
| 7Networks_LH_Vis_4 | -0.044392 | 0.5071 | -0.005032 | 0.96778 |
| 7Networks_LH_Vis_5 | 0.068566 | 0.27542 | -0.32723 | 0.01656 |
| 7Networks_LH_Vis_6 | -0.049526 | 0.49842 | 0.077775 | 0.4303 |
| 7Networks_LH_Vis_7 | -0.082042 | 0.24644 | -0.098443 | 0.30347 |
| 7Networks_LH_Vis_8 | 0.044844 | 0.48344 | -0.13776 | 0.27819 |
| 7Networks_LH_Vis_9 | 0.055499 | 0.38767 | -0.23594 | 0.036426 |
| 7Networks_LH_Vis_10 | -0.10793 | 0.16665 | 0.0018348 | 0.98049 |
| 7Networks_LH_Vis_11 | 0.064511 | 0.32262 | -0.24558 | 0.033938 |
| 7Networks_LH_Vis_12 | -0.025753 | 0.70912 | -0.13385 | 0.17447 |
| 7Networks_LH_Vis_13 | -0.064463 | 0.36988 | -0.10446 | 0.26678 |
| 7Networks_LH_Vis_14 | -0.048898 | 0.46615 | -0.04207 | 0.7114 |
| 7Networks_LH_SomMot_1 | -0.076778 | 0.24452 | -0.038622 | 0.72393 |
| 7Networks_LH_SomMot_2 | 0.0008146 | 0.98997 | -0.092779 | 0.3697 |
| 7Networks_LH_SomMot_3 | 0.018054 | 0.77707 | -0.28398 | 0.013342 |
| 7Networks_LH_SomMot_4 | 0.085146 | 0.16685 | -0.23141 | 0.075315 |
| 7Networks_LH_SomMot_5 | -0.035157 | 0.59359 | -0.14335 | 0.2291 |
| 7Networks_LH_SomMot_6 | -0.02985 | 0.64161 | 0.042341 | 0.6906 |
| 7Networks_LH_SomMot_7 | -0.024083 | 0.69076 | 0.022864 | 0.87396 |
| 7Networks_LH_SomMot_8 | -0.14819 | 0.019914 | 0.0056536 | 0.95762 |
| 7Networks_LH_SomMot_9 | 0.01725 | 0.77043 | 0.051745 | 0.74026 |
| 7Networks_LH_SomMot_10 | -0.010778 | 0.86019 | -0.014308 | 0.90597 |
| 7Networks_LH_SomMot_11 | -0.021496 | 0.74373 | -0.047575 | 0.70234 |
| 7Networks_LH_SomMot_12 | 0.035364 | 0.57395 | -0.11001 | 0.29967 |
| 7Networks_LH_SomMot_13 | 0.063942 | 0.33051 | 0.010001 | 0.93519 |
| 7Networks_LH_SomMot_14 | -0.041288 | 0.49909 | 0.12516 | 0.3461 |
| 7Networks_LH_SomMot_15 | -0.02531 | 0.68916 | -0.1102 | 0.34444 |
| 7Networks_LH_SomMot_16 | 0.013903 | 0.82439 | 0.0078213 | 0.94695 |
| 7Networks_LH_DorsAttn_Post_1 | -0.017881 | 0.77757 | 0.11987 | 0.34285 |
| 7Networks_LH_DorsAttn_Post_2 | 0.0007401 | 0.9907 | 0.071837 | 0.55312 |
| 7Networks_LH_DorsAttn_Post_3 | -0.034962 | 0.60915 | 0.0015223 | 0.98942 |
| 7Networks_LH_DorsAttn_Post_4 | 0.073607 | 0.25171 | -0.13035 | 0.31118 |
| 7Networks_LH_DorsAttn_Post_5 | 0.050632 | 0.39198 | -0.15701 | 0.324 |
| 7Networks_LH_DorsAttn_Post_6 | -0.014326 | 0.80385 | -0.15366 | 0.34172 |
| 7Networks_LH_DorsAttn_Post_7 | 0.085837 | 0.16261 | -0.35538 | 0.0046277 |
| 7Networks_LH_DorsAttn_Post_8 | 0.013204 | 0.8295 | -0.26377 | 0.043139 |
| 7Networks_LH_DorsAttn_Post_9 | -0.054496 | 0.39414 | 0.078058 | 0.49817 |
| 7Networks_LH_DorsAttn_Post_10 | -0.027494 | 0.66098 | 0.081968 | 0.51277 |
| 7Networks_LH_DorsAttn_FEF_1 | -0.011609 | 0.84392 | -0.1215 | 0.3731 |
| 7Networks_LH_DorsAttn_FEF_2 | -0.037084 | 0.5302 | -0.077946 | 0.59304 |
| 7Networks_LH_DorsAttn_PrCv_1 | -0.010397 | 0.86897 | -0.0010411 | 0.99372 |
| 7Networks_LH_SalVentAttn_ParOper_1 | -0.0048539 | 0.94004 | -0.016334 | 0.89624 |
| 7Networks_LH_SalVentAttn_ParOper_2 | -0.013706 | 0.82942 | -0.1527 | 0.17671 |
| 7Networks_LH_SalVentAttn_ParOper_3 | 0.069263 | 0.2712 | -0.28728 | 0.031832 |
| 7Networks_LH_SalVentAttn_FrOper_1 | -0.15545 | 0.027579 | 0.060235 | 0.52938 |
| 7Networks_LH_SalVentAttn_FrOper_2 | -0.066034 | 0.31567 | -0.059574 | 0.53105 |
| 7Networks_LH_SalVentAttn_FrOper_3 | 0.043113 | 0.51216 | -0.13577 | 0.18533 |
| 7Networks_LH_SalVentAttn_FrOper_4 | -0.071051 | 0.27831 | 0.0042003 | 0.97283 |
| 7Networks_LH_SalVentAttn_PFCl_1 | 0.1176 | 0.069099 | -0.25363 | 0.089451 |
| 7Networks_LH_SalVentAttn_Med_1 | 0.03792 | 0.57508 | -0.16407 | 0.10924 |
| 7Networks_LH_SalVentAttn_Med_2 | -0.039596 | 0.54699 | 0.018368 | 0.86962 |
| 7Networks_LH_SalVentAttn_Med_3 | -0.018042 | 0.78243 | -0.11494 | 0.39545 |
| 7Networks_LH_Limbic_OFC_1 | 0.10494 | 0.1138 | -0.21587 | 0.021362 |
| 7Networks_LH_Limbic_OFC_2 | -0.14544 | 0.03428 | 0.10447 | 0.28729 |
| 7Networks_LH_Limbic_TempPole_1 | -0.042746 | 0.53571 | -0.084881 | 0.41292 |
| 7Networks_LH_Limbic_TempPole_2 | -0.014598 | 0.81813 | -0.10046 | 0.3637 |
| 7Networks_LH_Limbic_TempPole_3 | 0.062929 | 0.33105 | -0.23766 | 0.038449 |
| 7Networks_LH_Limbic_TempPole_4 | -0.046309 | 0.50661 | -0.063595 | 0.50572 |
| 7Networks_LH_Cont_Par_1 | 0.053729 | 0.3901 | -0.028812 | 0.85614 |
| 7Networks_LH_Cont_Par_2 | -0.0062264 | 0.91793 | -0.10946 | 0.49338 |
| 7Networks_LH_Cont_Par_3 | 0.023033 | 0.69576 | -0.26238 | 0.11955 |
| 7Networks_LH_Cont_Temp_1 | 0.046658 | 0.45244 | -0.070999 | 0.58337 |
| 7Networks_LH_Cont_PFCl_1 | 0.13564 | 0.039244 | -0.2042 | 0.068791 |
| 7Networks_LH_Cont_PFCl_2 | 0.092026 | 0.16288 | -0.20975 | 0.06714 |
| 7Networks_LH_Cont_PFCl_3 | 0.12193 | 0.068026 | -0.38889 | 0.0018155 |
| 7Networks_LH_Cont_PFCl_4 | -0.043186 | 0.50906 | -0.021075 | 0.86127 |
| 7Networks_LH_Cont_PFCl_5 | -0.14928 | 0.019045 | 0.2326 | 0.06263 |
| 7Networks_LH_Cont_PFCl_6 | 0.066062 | 0.27093 | -0.063098 | 0.68299 |
| 7Networks_LH_Cont_pCun_1 | -0.055676 | 0.41382 | -0.032829 | 0.7658 |
| 7Networks_LH_Cont_Cing_1 | -0.060405 | 0.37419 | 0.055293 | 0.61104 |
| 7Networks_LH_Cont_Cing_2 | -0.081803 | 0.19706 | 0.11908 | 0.36255 |
| 7Networks_LH_Default_Temp_1 | -0.025705 | 0.72052 | -0.11327 | 0.22686 |
| 7Networks_LH_Default_Temp_2 | 0.0024858 | 0.96847 | -0.15921 | 0.21486 |
| 7Networks_LH_Default_Temp_3 | -0.057902 | 0.35185 | -0.010143 | 0.93113 |
| 7Networks_LH_Default_Temp_4 | -0.032886 | 0.62032 | -0.0044913 | 0.96723 |
| 7Networks_LH_Default_Temp_5 | -0.0094834 | 0.87853 | -0.032164 | 0.80301 |
| 7Networks_LH_Default_Temp_6 | -0.0076243 | 0.90682 | 0.048609 | 0.70976 |
| 7Networks_LH_Default_Temp_7 | 0.0088722 | 0.88836 | -0.065347 | 0.65434 |
| 7Networks_LH_Default_Temp_8 | 0.046756 | 0.42777 | 0.08537 | 0.65547 |
| 7Networks_LH_Default_Temp_9 | -0.03457 | 0.57982 | 0.048989 | 0.70277 |
| 7Networks_LH_Default_PFC_1 | 0.053594 | 0.38251 | -0.18134 | 0.096197 |
| 7Networks_LH_Default_PFC_2 | 0.011244 | 0.8666 | -0.043052 | 0.64524 |
| 7Networks_LH_Default_PFC_3 | -0.04128 | 0.51915 | -0.17887 | 0.1152 |
| 7Networks_LH_Default_PFC_4 | 0.053885 | 0.43032 | -0.23373 | 0.029489 |
| 7Networks_LH_Default_PFC_5 | -0.062688 | 0.3251 | 0.026551 | 0.82514 |
| 7Networks_LH_Default_PFC_6 | 0.026905 | 0.69755 | -0.11478 | 0.242 |
| 7Networks_LH_Default_PFC_7 | 0.038663 | 0.56658 | -0.21795 | 0.071068 |
| 7Networks_LH_Default_PFC_8 | 0.070259 | 0.27969 | -0.23649 | 0.046337 |
| 7Networks_LH_Default_PFC_9 | 0.12626 | 0.067301 | -0.44947 | 0.0001624 |
| 7Networks_LH_Default_PFC_10 | -0.021217 | 0.74826 | -0.03496 | 0.75828 |
| 7Networks_LH_Default_PFC_11 | 0.017333 | 0.78044 | -0.0061739 | 0.9614 |
| 7Networks_LH_Default_PFC_12 | 0.013832 | 0.8283 | -0.29848 | 0.027895 |
| 7Networks_LH_Default_PFC_13 | -0.034521 | 0.60221 | -0.21233 | 0.13829 |
| 7Networks_LH_Default_PCC_1 | -0.1366 | 0.059954 | -0.0044891 | 0.95867 |
| 7Networks_LH_Default_PCC_2 | -0.032865 | 0.62355 | -0.036737 | 0.74269 |
| 7Networks_LH_Default_PCC_3 | -0.11484 | 0.071416 | 0.18155 | 0.13969 |
| 7Networks_LH_Default_PCC_4 | 0.083227 | 0.20342 | -0.15889 | 0.18773 |
| 7Networks_LH_Default_PHC_1 | -0.085015 | 0.19799 | 0.051284 | 0.59201 |
| 7Networks_RH_Vis_1 | -0.0090572 | 0.89264 | -0.09589 | 0.34655 |
| 7Networks_RH_Vis_2 | -0.0010422 | 0.98754 | -0.085869 | 0.4207 |
| 7Networks_RH_Vis_3 | 0.0008169 | 0.9901 | -0.0009071 | 0.99352 |
| 7Networks_RH_Vis_4 | 0.08874 | 0.17687 | -0.12606 | 0.22647 |
| 7Networks_RH_Vis_5 | 0.10872 | 0.075571 | -0.23694 | 0.11948 |
| 7Networks_RH_Vis_6 | -0.0063639 | 0.93232 | -0.080122 | 0.40454 |
| 7Networks_RH_Vis_7 | 0.055073 | 0.43398 | -0.075124 | 0.42912 |
| 7Networks_RH_Vis_8 | 0.063091 | 0.3392 | -0.23733 | 0.052102 |
| 7Networks_RH_Vis_9 | 0.020724 | 0.78721 | -0.10093 | 0.21988 |
| 7Networks_RH_Vis_10 | -0.050419 | 0.51946 | -0.020325 | 0.8004 |
| 7Networks_RH_Vis_11 | 0.06432 | 0.29713 | -0.26007 | 0.075173 |
| 7Networks_RH_Vis_12 | -0.0034789 | 0.95879 | -0.19743 | 0.053529 |
| 7Networks_RH_Vis_13 | -0.027655 | 0.6847 | -0.04703 | 0.65987 |
| 7Networks_RH_Vis_14 | -0.044214 | 0.49929 | -0.10757 | 0.37146 |
| 7Networks_RH_Vis_15 | -0.029018 | 0.63997 | -0.031736 | 0.81537 |
| 7Networks_RH_SomMot_1 | 0.02648 | 0.70317 | -0.12018 | 0.21238 |
| 7Networks_RH_SomMot_2 | -0.040322 | 0.56262 | -0.023045 | 0.81763 |
| 7Networks_RH_SomMot_3 | -0.039919 | 0.55976 | -0.02528 | 0.81881 |
| 7Networks_RH_SomMot_4 | 0.046051 | 0.47917 | -0.12583 | 0.25101 |
| 7Networks_RH_SomMot_5 | -0.031908 | 0.60409 | -0.074198 | 0.59647 |
| 7Networks_RH_SomMot_6 | 0.03119 | 0.65124 | -0.10541 | 0.35471 |
| 7Networks_RH_SomMot_7 | -0.051419 | 0.42204 | 0.090707 | 0.4382 |
| 7Networks_RH_SomMot_8 | 0.01113 | 0.85946 | -0.02914 | 0.82159 |
| 7Networks_RH_SomMot_9 | 0.10168 | 0.088741 | -0.29094 | 0.073851 |
| 7Networks_RH_SomMot_10 | -0.051179 | 0.3997 | 0.10456 | 0.42854 |
| 7Networks_RH_SomMot_11 | -0.079086 | 0.24353 | 0.091747 | 0.36898 |
| 7Networks_RH_SomMot_12 | 0.0067991 | 0.91282 | -0.021116 | 0.86151 |
| 7Networks_RH_SomMot_13 | -0.12111 | 0.061427 | 0.020454 | 0.88083 |
| 7Networks_RH_SomMot_14 | 0.018738 | 0.77188 | -0.021129 | 0.84951 |
| 7Networks_RH_SomMot_15 | 0.013518 | 0.82058 | -0.19047 | 0.24022 |
| 7Networks_RH_SomMot_16 | -0.093175 | 0.1368 | -0.025386 | 0.85512 |
| 7Networks_RH_SomMot_17 | -0.051202 | 0.39253 | -0.035372 | 0.84616 |
| 7Networks_RH_SomMot_18 | 0.0045581 | 0.9445 | -0.0066326 | 0.95168 |
| 7Networks_RH_SomMot_19 | -0.031888 | 0.62013 | 0.11098 | 0.31905 |
| 7Networks_RH_DorsAttn_Post_1 | -0.012253 | 0.85284 | 0.01053 | 0.92181 |
| 7Networks_RH_DorsAttn_Post_2 | 0.033748 | 0.59284 | 0.10842 | 0.41418 |
| 7Networks_RH_DorsAttn_Post_3 | 0.060908 | 0.32946 | -0.26228 | 0.023825 |
| 7Networks_RH_DorsAttn_Post_4 | 0.04499 | 0.44882 | -0.22893 | 0.12571 |
| 7Networks_RH_DorsAttn_Post_5 | 0.043718 | 0.47627 | -0.21245 | 0.12957 |
| 7Networks_RH_DorsAttn_Post_6 | -0.014696 | 0.81045 | -0.12251 | 0.41561 |
| 7Networks_RH_DorsAttn_Post_7 | 0.061069 | 0.33201 | -0.091905 | 0.56402 |
| 7Networks_RH_DorsAttn_Post_8 | -0.065514 | 0.2817 | 0.053112 | 0.73169 |
| 7Networks_RH_DorsAttn_Post_9 | -0.05487 | 0.39804 | -0.043748 | 0.71156 |
| 7Networks_RH_DorsAttn_Post_10 | -0.04124 | 0.5169 | -0.061069 | 0.62433 |
| 7Networks_RH_DorsAttn_FEF_1 | -0.0017255 | 0.9771 | -0.15985 | 0.33785 |
| 7Networks_RH_DorsAttn_FEF_2 | -0.039873 | 0.56145 | -0.11754 | 0.25536 |
| 7Networks_RH_DorsAttn_PrCv_1 | 0.069436 | 0.2618 | -0.15686 | 0.23001 |
| 7Networks_RH_SalVentAttn_TempOccPar_1 | -0.087508 | 0.16428 | 0.13158 | 0.31911 |
| 7Networks_RH_SalVentAttn_TempOccPar_2 | -0.066531 | 0.26438 | -0.051521 | 0.74503 |
| 7Networks_RH_SalVentAttn_TempOccPar_3 | -0.015066 | 0.81542 | -0.14408 | 0.22929 |
| 7Networks_RH_SalVentAttn_PrC_1 | -0.0084897 | 0.88978 | -0.0025178 | 0.98555 |
| 7Networks_RH_SalVentAttn_FrOper_1 | -0.09135 | 0.18042 | 0.065912 | 0.50056 |
| 7Networks_RH_SalVentAttn_FrOper_2 | -0.035142 | 0.60022 | -0.087122 | 0.38486 |
| 7Networks_RH_SalVentAttn_FrOper_3 | -0.078756 | 0.21317 | 0.023856 | 0.83429 |
| 7Networks_RH_SalVentAttn_FrOper_4 | -0.092243 | 0.16224 | 0.02245 | 0.83889 |
| 7Networks_RH_SalVentAttn_Med_1 | 0.13653 | 0.048741 | -0.25505 | 0.0099399 |
| 7Networks_RH_SalVentAttn_Med_2 | -0.017176 | 0.79506 | 0.10676 | 0.30106 |
| 7Networks_RH_SalVentAttn_Med_3 | -0.077997 | 0.2409 | -0.025791 | 0.81101 |
| 7Networks_RH_Limbic_OFC_1 | -0.013789 | 0.83873 | 0.04368 | 0.66669 |
| 7Networks_RH_Limbic_OFC_2 | 0.0066139 | 0.9177 | -0.057564 | 0.57811 |
| 7Networks_RH_Limbic_OFC_3 | 0.11219 | 0.10474 | -0.35027 | 0.0027319 |
| 7Networks_RH_Limbic_TempPole_1 | -0.022448 | 0.72915 | -0.19846 | 0.057436 |
| 7Networks_RH_Limbic_TempPole_2 | -0.058646 | 0.3616 | -0.084979 | 0.43926 |
| 7Networks_RH_Limbic_TempPole_3 | -0.096668 | 0.13601 | 0.11253 | 0.31484 |
| 7Networks_RH_Cont_Par_1 | -0.052905 | 0.38925 | -0.077826 | 0.58802 |
| 7Networks_RH_Cont_Par_2 | 0.094088 | 0.12982 | -0.27698 | 0.09667 |
| 7Networks_RH_Cont_Par_3 | 0.02238 | 0.70448 | -0.094327 | 0.55689 |
| 7Networks_RH_Cont_Temp_1 | -0.039225 | 0.55213 | -0.005503 | 0.96631 |
| 7Networks_RH_Cont_PFCv_1 | -0.0004562 | 0.99382 | 0.19048 | 0.14399 |
| 7Networks_RH_Cont_PFCl_1 | 0.06055 | 0.35242 | -0.16411 | 0.101 |
| 7Networks_RH_Cont_PFCl_2 | 0.077949 | 0.23229 | -0.31498 | 0.018201 |
| 7Networks_RH_Cont_PFCl_3 | 0.025399 | 0.69416 | -0.098645 | 0.40747 |
| 7Networks_RH_Cont_PFCl_4 | 0.17096 | 0.0081334 | -0.1758 | 0.13948 |
| 7Networks_RH_Cont_PFCl_5 | -0.017293 | 0.78512 | -0.17221 | 0.20318 |
| 7Networks_RH_Cont_PFCl_6 | 0.048917 | 0.42558 | -0.10142 | 0.47245 |
| 7Networks_RH_Cont_PFCl_7 | 0.052543 | 0.38908 | -0.031367 | 0.79047 |
| 7Networks_RH_Cont_pCun_1 | 0.0062329 | 0.92625 | -0.019425 | 0.87146 |
| 7Networks_RH_Cont_PFCmp_1 | -0.0021564 | 0.97361 | -0.098033 | 0.37439 |
| 7Networks_RH_Cont_PFCmp_2 | 0.051629 | 0.42806 | -0.14421 | 0.23238 |
| 7Networks_RH_Cont_PFCmp_3 | 0.080189 | 0.23938 | -0.21414 | 0.033501 |
| 7Networks_RH_Cont_PFCmp_4 | -0.12363 | 0.063213 | -0.016624 | 0.88425 |
| 7Networks_RH_Default_Par_1 | 0.015325 | 0.8089 | 0.0032329 | 0.98049 |
| 7Networks_RH_Default_Par_2 | -0.0078756 | 0.89634 | -0.07164 | 0.67004 |
| 7Networks_RH_Default_Par_3 | -0.018922 | 0.75412 | -0.027298 | 0.87948 |
| 7Networks_RH_Default_Temp_1 | -0.15259 | 0.017356 | -0.0027416 | 0.97895 |
| 7Networks_RH_Default_Temp_2 | 0.010553 | 0.87041 | -0.22569 | 0.048293 |
| 7Networks_RH_Default_Temp_3 | 0.0095789 | 0.87584 | -0.088825 | 0.43265 |
| 7Networks_RH_Default_Temp_4 | 0.017187 | 0.7854 | -0.099807 | 0.42694 |
| 7Networks_RH_Default_Temp_5 | -0.05424 | 0.41005 | 0.065554 | 0.55837 |
| 7Networks_RH_Default_PFCv_1 | 0.018189 | 0.7662 | -0.14404 | 0.24444 |
| 7Networks_RH_Default_PFCm_1 | 0.077812 | 0.27945 | -0.18106 | 0.069221 |
| 7Networks_RH_Default_PFCm_2 | 0.039252 | 0.57298 | -0.11694 | 0.29276 |
| 7Networks_RH_Default_PFCm_3 | 0.039935 | 0.52961 | -0.060825 | 0.73767 |
| 7Networks_RH_Default_PFCm_4 | 0.018019 | 0.79668 | -0.25123 | 0.016384 |
| 7Networks_RH_Default_PFCm_5 | -0.038041 | 0.54762 | -0.29632 | 0.015366 |
| 7Networks_RH_Default_PFCm_6 | 0.038207 | 0.53776 | -0.21663 | 0.089802 |
| 7Networks_RH_Default_PFCm_7 | 0.025745 | 0.67941 | -0.2438 | 0.080139 |
| 7Networks_RH_Default_PCC_1 | -0.0038321 | 0.95803 | 0.0043315 | 0.96213 |
| 7Networks_RH_Default_PCC_2 | 0.021832 | 0.74209 | -0.12838 | 0.29141 |
| 7Networks_RH_Default_PCC_3 | -0.0065024 | 0.92027 | -0.046126 | 0.73657 |

**Supplementary Table 15. Genetic and environmental correlation between Neuroticism and surface area in HCP.** Genetic correlation between Neuroticism and local surface area calculated using solar 8.4.0. Regions are named according to the Schaefer 200 atlas, based on 7-networks. Here we report environmental correlation (ρ_e_) and genetic correlation (ρ_g_) and the associated p-values.

| ROIS | ρ_e_ | p | ρ_g_ | p |
| --- | --- | --- | --- | --- |
| 7Networks_LH_Vis_1 | -0.068007 | 0.34042 | 0.029448 | 0.69411 |
| 7Networks_LH_Vis_2 | 0.0066816 | 0.92484 | -0.0053475 | 0.94698 |
| 7Networks_LH_Vis_3 | 0.016102 | 0.80347 | 0.083454 | 0.36972 |
| 7Networks_LH_Vis_4 | -0.17496 | 0.0092098 | 0.22772 | 0.015307 |
| 7Networks_LH_Vis_5 | 0.0026652 | 0.96733 | 0.087098 | 0.39278 |
| 7Networks_LH_Vis_6 | -0.073012 | 0.31484 | -0.013772 | 0.85017 |
| 7Networks_LH_Vis_7 | 0.010809 | 0.878 | -0.034599 | 0.62556 |
| 7Networks_LH_Vis_8 | 0.086478 | 0.18713 | 0.035793 | 0.70227 |
| 7Networks_LH_Vis_9 | -0.034676 | 0.59568 | 0.057689 | 0.48778 |
| 7Networks_LH_Vis_10 | 0.015371 | 0.8376 | -0.045368 | 0.42607 |
| 7Networks_LH_Vis_11 | -0.054648 | 0.4115 | 0.098965 | 0.23771 |
| 7Networks_LH_Vis_12 | 0.1344 | 0.051629 | -0.091552 | 0.20821 |
| 7Networks_LH_Vis_13 | 0.070589 | 0.32136 | -0.072898 | 0.29531 |
| 7Networks_LH_Vis_14 | 0.047989 | 0.48213 | -0.071846 | 0.39428 |
| 7Networks_LH_SomMot_1 | 0.049176 | 0.46306 | -0.029461 | 0.71735 |
| 7Networks_LH_SomMot_2 | 0.042723 | 0.51304 | -0.05971 | 0.43632 |
| 7Networks_LH_SomMot_3 | -0.016193 | 0.80244 | 0.022897 | 0.79096 |
| 7Networks_LH_SomMot_4 | -0.011805 | 0.85161 | 0.022589 | 0.81584 |
| 7Networks_LH_SomMot_5 | -0.0042463 | 0.94933 | 0.043267 | 0.62444 |
| 7Networks_LH_SomMot_6 | 0.0066004 | 0.91873 | 0.046303 | 0.55773 |
| 7Networks_LH_SomMot_7 | 0.029598 | 0.63263 | 0.022985 | 0.82883 |
| 7Networks_LH_SomMot_8 | 0.0314 | 0.62501 | -0.0018212 | 0.98167 |
| 7Networks_LH_SomMot_9 | -0.062258 | 0.30833 | 0.15047 | 0.19943 |
| 7Networks_LH_SomMot_10 | -0.025632 | 0.68153 | -0.028252 | 0.75566 |
| 7Networks_LH_SomMot_11 | 0.053526 | 0.42288 | 0.09126 | 0.32043 |
| 7Networks_LH_SomMot_12 | -0.049916 | 0.43386 | 0.075903 | 0.33672 |
| 7Networks_LH_SomMot_13 | 0.075649 | 0.25332 | 0.099404 | 0.267 |
| 7Networks_LH_SomMot_14 | -0.014417 | 0.81649 | 0.045136 | 0.652 |
| 7Networks_LH_SomMot_15 | -0.066565 | 0.30247 | 0.038974 | 0.65296 |
| 7Networks_LH_SomMot_16 | -0.019692 | 0.75733 | 0.084106 | 0.33638 |
| 7Networks_LH_DorsAttn_Post_1 | 0.0098909 | 0.87865 | 0.14843 | 0.11709 |
| 7Networks_LH_DorsAttn_Post_2 | 0.16224 | 0.012589 | -0.016426 | 0.8578 |
| 7Networks_LH_DorsAttn_Post_3 | 0.060024 | 0.38701 | 0.0039345 | 0.96329 |
| 7Networks_LH_DorsAttn_Post_4 | -0.045331 | 0.49133 | 0.0063049 | 0.94655 |
| 7Networks_LH_DorsAttn_Post_5 | 0.030519 | 0.61734 | 0.043285 | 0.71072 |
| 7Networks_LH_DorsAttn_Post_6 | 0.11712 | 0.049935 | -0.14805 | 0.21857 |
| 7Networks_LH_DorsAttn_Post_7 | 0.12796 | 0.045229 | -0.22529 | 0.014342 |
| 7Networks_LH_DorsAttn_Post_8 | 0.063548 | 0.3109 | -0.060276 | 0.53753 |
| 7Networks_LH_DorsAttn_Post_9 | 0.007845 | 0.90435 | 0.056455 | 0.51214 |
| 7Networks_LH_DorsAttn_Post_10 | -0.0092061 | 0.88576 | 0.12458 | 0.18291 |
| 7Networks_LH_DorsAttn_FEF_1 | 0.029642 | 0.62476 | -0.02677 | 0.79394 |
| 7Networks_LH_DorsAttn_FEF_2 | -0.034236 | 0.57237 | 0.095313 | 0.38335 |
| 7Networks_LH_DorsAttn_PrCv_1 | 0.015542 | 0.81007 | 0.059391 | 0.55349 |
| 7Networks_LH_SalVentAttn_ParOper_1 | 0.048738 | 0.45959 | 0.017298 | 0.85249 |
| 7Networks_LH_SalVentAttn_ParOper_2 | 0.0031256 | 0.96127 | 0.044207 | 0.60075 |
| 7Networks_LH_SalVentAttn_ParOper_3 | 0.073775 | 0.25099 | -0.028616 | 0.77562 |
| 7Networks_LH_SalVentAttn_FrOper_1 | -0.027864 | 0.68969 | 0.13208 | 0.063104 |
| 7Networks_LH_SalVentAttn_FrOper_2 | 0.10967 | 0.094759 | -0.047348 | 0.51976 |
| 7Networks_LH_SalVentAttn_FrOper_3 | 0.01294 | 0.84428 | -0.014995 | 0.8451 |
| 7Networks_LH_SalVentAttn_FrOper_4 | 0.040221 | 0.54753 | -0.073773 | 0.41816 |
| 7Networks_LH_SalVentAttn_PFCl_1 | 0.018861 | 0.7786 | 0.060733 | 0.58161 |
| 7Networks_LH_SalVentAttn_Med_1 | 0.14558 | 0.032317 | -0.055851 | 0.45529 |
| 7Networks_LH_SalVentAttn_Med_2 | 0.0086782 | 0.8966 | -0.04317 | 0.60589 |
| 7Networks_LH_SalVentAttn_Med_3 | 0.021918 | 0.74328 | -0.041034 | 0.68189 |
| 7Networks_LH_Limbic_OFC_1 | 0.10695 | 0.10996 | -0.042555 | 0.55359 |
| 7Networks_LH_Limbic_OFC_2 | 0.094751 | 0.16954 | -0.030863 | 0.67643 |
| 7Networks_LH_Limbic_TempPole_1 | -0.028563 | 0.6765 | 0.12558 | 0.10078 |
| 7Networks_LH_Limbic_TempPole_2 | -0.054979 | 0.3935 | 0.11515 | 0.1653 |
| 7Networks_LH_Limbic_TempPole_3 | 0.084772 | 0.18901 | 0.0027584 | 0.97465 |
| 7Networks_LH_Limbic_TempPole_4 | -0.032196 | 0.64248 | 0.09903 | 0.16247 |
| 7Networks_LH_Cont_Par_1 | -0.0032301 | 0.95996 | -0.019337 | 0.869 |
| 7Networks_LH_Cont_Par_2 | -0.031767 | 0.60892 | 0.051573 | 0.66639 |
| 7Networks_LH_Cont_Par_3 | 0.048085 | 0.43317 | -0.11791 | 0.33812 |
| 7Networks_LH_Cont_Temp_1 | 0.071285 | 0.26425 | 0.0497 | 0.61397 |
| 7Networks_LH_Cont_PFCl_1 | 0.043948 | 0.51407 | -0.075792 | 0.36538 |
| 7Networks_LH_Cont_PFCl_2 | 0.043742 | 0.51662 | -0.044998 | 0.59867 |
| 7Networks_LH_Cont_PFCl_3 | 0.018194 | 0.78875 | -0.039825 | 0.67402 |
| 7Networks_LH_Cont_PFCl_4 | 0.049934 | 0.45111 | 0.026807 | 0.76431 |
| 7Networks_LH_Cont_PFCl_5 | 0.15261 | 0.017027 | -0.13916 | 0.14681 |
| 7Networks_LH_Cont_PFCl_6 | 0.038903 | 0.53012 | 0.13405 | 0.24569 |
| 7Networks_LH_Cont_pCun_1 | -0.072223 | 0.29603 | 0.098433 | 0.22469 |
| 7Networks_LH_Cont_Cing_1 | 0.16042 | 0.018783 | -0.10148 | 0.20558 |
| 7Networks_LH_Cont_Cing_2 | 0.058905 | 0.36022 | -0.0045949 | 0.96259 |
| 7Networks_LH_Default_Temp_1 | 0.095682 | 0.18045 | 0.027457 | 0.69335 |
| 7Networks_LH_Default_Temp_2 | 0.013842 | 0.82908 | 0.13308 | 0.16536 |
| 7Networks_LH_Default_Temp_3 | 0.026116 | 0.68812 | -0.10408 | 0.24013 |
| 7Networks_LH_Default_Temp_4 | 0.03664 | 0.59007 | -0.035119 | 0.66903 |
| 7Networks_LH_Default_Temp_5 | -0.02404 | 0.7064 | 0.13413 | 0.163 |
| 7Networks_LH_Default_Temp_6 | 0.080458 | 0.22827 | 0.063839 | 0.51437 |
| 7Networks_LH_Default_Temp_7 | -0.0304 | 0.64081 | 0.1473 | 0.16533 |
| 7Networks_LH_Default_Temp_8 | -0.040661 | 0.50741 | 0.134 | 0.34729 |
| 7Networks_LH_Default_Temp_9 | 0.080692 | 0.20416 | -0.062488 | 0.51992 |
| 7Networks_LH_Default_PFC_1 | 0.048059 | 0.43864 | -0.041559 | 0.60659 |
| 7Networks_LH_Default_PFC_2 | 0.0025262 | 0.96981 | -0.028494 | 0.68515 |
| 7Networks_LH_Default_PFC_3 | -0.040131 | 0.54066 | 0.035786 | 0.67128 |
| 7Networks_LH_Default_PFC_4 | 0.055111 | 0.41422 | -0.041856 | 0.60504 |
| 7Networks_LH_Default_PFC_5 | 0.10167 | 0.11405 | -0.14129 | 0.11689 |
| 7Networks_LH_Default_PFC_6 | 0.060641 | 0.3806 | -0.019982 | 0.78218 |
| 7Networks_LH_Default_PFC_7 | 0.10045 | 0.1444 | -0.074672 | 0.40106 |
| 7Networks_LH_Default_PFC_8 | 0.13497 | 0.042446 | -0.047906 | 0.57546 |
| 7Networks_LH_Default_PFC_9 | 0.0072077 | 0.91532 | 0.039641 | 0.67218 |
| 7Networks_LH_Default_PFC_10 | -0.06919 | 0.29544 | 0.062071 | 0.45733 |
| 7Networks_LH_Default_PFC_11 | 0.069642 | 0.27311 | -0.011654 | 0.9024 |
| 7Networks_LH_Default_PFC_12 | 0.013126 | 0.83881 | -0.045453 | 0.65799 |
| 7Networks_LH_Default_PFC_13 | 0.011048 | 0.87092 | 0.028849 | 0.78514 |
| 7Networks_LH_Default_PCC_1 | 0.058833 | 0.41411 | -0.088013 | 0.1722 |
| 7Networks_LH_Default_PCC_2 | 0.039639 | 0.56057 | -0.0041223 | 0.96077 |
| 7Networks_LH_Default_PCC_3 | 0.14655 | 0.023179 | -0.14556 | 0.11527 |
| 7Networks_LH_Default_PCC_4 | -0.0019279 | 0.97696 | 0.049436 | 0.58326 |
| 7Networks_LH_Default_PHC_1 | -0.0087008 | 0.89509 | 0.01137 | 0.87473 |
| 7Networks_RH_Vis_1 | 0.017079 | 0.80047 | -0.04894 | 0.51798 |
| 7Networks_RH_Vis_2 | -0.013538 | 0.84073 | 0.11212 | 0.15501 |
| 7Networks_RH_Vis_3 | 0.10699 | 0.10887 | -0.010452 | 0.90505 |
| 7Networks_RH_Vis_4 | -0.08046 | 0.22471 | 0.17222 | 0.029979 |
| 7Networks_RH_Vis_5 | 0.045442 | 0.47601 | 0.1455 | 0.19109 |
| 7Networks_RH_Vis_6 | -0.010686 | 0.88477 | -0.0035346 | 0.95992 |
| 7Networks_RH_Vis_7 | -0.089125 | 0.20311 | 0.0009493 | 0.98924 |
| 7Networks_RH_Vis_8 | 0.031107 | 0.65166 | 0.038879 | 0.67616 |
| 7Networks_RH_Vis_9 | 0.060809 | 0.41931 | -0.04703 | 0.4396 |
| 7Networks_RH_Vis_10 | 0.0035838 | 0.96202 | -0.094545 | 0.11483 |
| 7Networks_RH_Vis_11 | 0.047501 | 0.46241 | 0.020103 | 0.85272 |
| 7Networks_RH_Vis_12 | -0.012149 | 0.85691 | 0.021351 | 0.77971 |
| 7Networks_RH_Vis_13 | 0.057383 | 0.40491 | -0.028162 | 0.72141 |
| 7Networks_RH_Vis_14 | 0.030948 | 0.6444 | -0.044582 | 0.61743 |
| 7Networks_RH_Vis_15 | 0.050314 | 0.4286 | 0.0004017 | 0.99661 |
| 7Networks_RH_SomMot_1 | 0.020398 | 0.768 | -0.12004 | 0.0879 |
| 7Networks_RH_SomMot_2 | -0.01268 | 0.8551 | -0.018725 | 0.80051 |
| 7Networks_RH_SomMot_3 | 0.030292 | 0.66035 | 0.0021707 | 0.97889 |
| 7Networks_RH_SomMot_4 | 0.086116 | 0.18907 | -0.10128 | 0.20934 |
| 7Networks_RH_SomMot_5 | 0.085564 | 0.17596 | -0.12314 | 0.24325 |
| 7Networks_RH_SomMot_6 | -0.065261 | 0.34916 | 0.080293 | 0.33491 |
| 7Networks_RH_SomMot_7 | 0.078803 | 0.2263 | -0.076355 | 0.37417 |
| 7Networks_RH_SomMot_8 | 0.00347 | 0.95701 | 0.059669 | 0.53454 |
| 7Networks_RH_SomMot_9 | 0.076984 | 0.21682 | -0.10956 | 0.36627 |
| 7Networks_RH_SomMot_10 | 0.12832 | 0.039391 | -0.025935 | 0.79231 |
| 7Networks_RH_SomMot_11 | -0.013643 | 0.84123 | 0.030977 | 0.68546 |
| 7Networks_RH_SomMot_12 | 0.0064861 | 0.91863 | 0.068166 | 0.45202 |
| 7Networks_RH_SomMot_13 | -0.085365 | 0.19568 | 0.15402 | 0.11572 |
| 7Networks_RH_SomMot_14 | 0.050392 | 0.44191 | 0.10659 | 0.20021 |
| 7Networks_RH_SomMot_15 | -0.055944 | 0.36223 | 0.21942 | 0.07195 |
| 7Networks_RH_SomMot_16 | -0.012345 | 0.84797 | 0.0049661 | 0.96155 |
| 7Networks_RH_SomMot_17 | -0.033286 | 0.59455 | 0.14299 | 0.28804 |
| 7Networks_RH_SomMot_18 | -0.074603 | 0.25607 | 0.10418 | 0.19728 |
| 7Networks_RH_SomMot_19 | 0.018226 | 0.77622 | 0.072234 | 0.38751 |
| 7Networks_RH_DorsAttn_Post_1 | -0.045211 | 0.4959 | 0.057267 | 0.47408 |
| 7Networks_RH_DorsAttn_Post_2 | -0.02301 | 0.72175 | 0.12618 | 0.2008 |
| 7Networks_RH_DorsAttn_Post_3 | 0.080868 | 0.20385 | -0.079285 | 0.36217 |
| 7Networks_RH_DorsAttn_Post_4 | 0.035117 | 0.56632 | -0.033372 | 0.76726 |
| 7Networks_RH_DorsAttn_Post_5 | 0.013895 | 0.8256 | -0.001846 | 0.98573 |
| 7Networks_RH_DorsAttn_Post_6 | 0.059758 | 0.3445 | -0.14899 | 0.18343 |
| 7Networks_RH_DorsAttn_Post_7 | -0.031564 | 0.62571 | 0.036058 | 0.76171 |
| 7Networks_RH_DorsAttn_Post_8 | 0.03828 | 0.54588 | -0.048781 | 0.67234 |
| 7Networks_RH_DorsAttn_Post_9 | 0.0021054 | 0.97457 | -0.0094916 | 0.91386 |
| 7Networks_RH_DorsAttn_Post_10 | 0.0014683 | 0.98195 | -0.0099315 | 0.91445 |
| 7Networks_RH_DorsAttn_FEF_1 | 0.079162 | 0.20956 | -0.12121 | 0.32557 |
| 7Networks_RH_DorsAttn_FEF_2 | 0.016121 | 0.81496 | 0.025683 | 0.737 |
| 7Networks_RH_DorsAttn_PrCv_1 | 0.0009034 | 0.98849 | -0.01195 | 0.9023 |
| 7Networks_RH_SalVentAttn_TempOccPar_1 | -0.081272 | 0.2064 | 0.13702 | 0.16425 |
| 7Networks_RH_SalVentAttn_TempOccPar_2 | 0.068953 | 0.26021 | -0.11203 | 0.34809 |
| 7Networks_RH_SalVentAttn_TempOccPar_3 | 0.085953 | 0.19188 | -0.11376 | 0.198 |
| 7Networks_RH_SalVentAttn_PrC_1 | 0.11656 | 0.061621 | -0.048999 | 0.63967 |
| 7Networks_RH_SalVentAttn_FrOper_1 | 0.11168 | 0.10177 | -0.002085 | 0.97702 |
| 7Networks_RH_SalVentAttn_FrOper_2 | 0.022845 | 0.73248 | 0.13909 | 0.061821 |
| 7Networks_RH_SalVentAttn_FrOper_3 | 0.13268 | 0.039311 | -0.090385 | 0.29176 |
| 7Networks_RH_SalVentAttn_FrOper_4 | 0.087709 | 0.18509 | 0.055962 | 0.50167 |
| 7Networks_RH_SalVentAttn_Med_1 | 0.015248 | 0.82659 | 0.086828 | 0.23975 |
| 7Networks_RH_SalVentAttn_Med_2 | 0.029646 | 0.65818 | 0.046992 | 0.54443 |
| 7Networks_RH_SalVentAttn_Med_3 | 0.10795 | 0.10521 | -0.10479 | 0.19415 |
| 7Networks_RH_Limbic_OFC_1 | 0.038044 | 0.57509 | 0.011916 | 0.87501 |
| 7Networks_RH_Limbic_OFC_2 | 0.11262 | 0.079996 | -0.1011 | 0.20309 |
| 7Networks_RH_Limbic_OFC_3 | 0.042355 | 0.53823 | 0.065519 | 0.4556 |
| 7Networks_RH_Limbic_TempPole_1 | -0.081619 | 0.20897 | 0.096619 | 0.21866 |
| 7Networks_RH_Limbic_TempPole_2 | -0.029244 | 0.65099 | 0.031499 | 0.70019 |
| 7Networks_RH_Limbic_TempPole_3 | 0.013965 | 0.83319 | 0.089576 | 0.28226 |
| 7Networks_RH_Cont_Par_1 | 0.084728 | 0.18141 | 0.052031 | 0.62808 |
| 7Networks_RH_Cont_Par_2 | 0.0578 | 0.36977 | 0.025362 | 0.83899 |
| 7Networks_RH_Cont_Par_3 | -0.003623 | 0.95265 | 0.10939 | 0.36095 |
| 7Networks_RH_Cont_Temp_1 | -0.096918 | 0.15315 | 0.2503 | 0.0089107 |
| 7Networks_RH_Cont_PFCv_1 | 0.0042992 | 0.94441 | 0.016241 | 0.86868 |
| 7Networks_RH_Cont_PFCl_1 | 0.032386 | 0.62269 | -0.055774 | 0.46138 |
| 7Networks_RH_Cont_PFCl_2 | 0.15474 | 0.019795 | -0.096092 | 0.33492 |
| 7Networks_RH_Cont_PFCl_3 | 0.053398 | 0.41721 | 0.026933 | 0.76123 |
| 7Networks_RH_Cont_PFCl_4 | 0.050323 | 0.45038 | -0.018167 | 0.83454 |
| 7Networks_RH_Cont_PFCl_5 | 0.14369 | 0.027382 | -0.081631 | 0.41532 |
| 7Networks_RH_Cont_PFCl_6 | 0.089408 | 0.15839 | -0.0042561 | 0.9675 |
| 7Networks_RH_Cont_PFCl_7 | -0.022655 | 0.7148 | 0.0066978 | 0.93934 |
| 7Networks_RH_Cont_pCun_1 | 0.036178 | 0.59828 | -0.072584 | 0.4116 |
| 7Networks_RH_Cont_PFCmp_1 | 0.10097 | 0.13419 | -0.06476 | 0.44105 |
| 7Networks_RH_Cont_PFCmp_2 | 0.086079 | 0.19602 | -0.054674 | 0.54473 |
| 7Networks_RH_Cont_PFCmp_3 | 0.056229 | 0.40926 | 0.058532 | 0.4305 |
| 7Networks_RH_Cont_PFCmp_4 | 0.058874 | 0.37771 | 0.028386 | 0.74014 |
| 7Networks_RH_Default_Par_1 | 0.066239 | 0.30002 | -0.011591 | 0.90546 |
| 7Networks_RH_Default_Par_2 | -0.047016 | 0.45552 | 0.077472 | 0.53418 |
| 7Networks_RH_Default_Par_3 | -0.019326 | 0.75842 | 0.16881 | 0.20261 |
| 7Networks_RH_Default_Temp_1 | 0.13187 | 0.040662 | -0.049892 | 0.52591 |
| 7Networks_RH_Default_Temp_2 | 0.037192 | 0.56147 | 0.032262 | 0.70739 |
| 7Networks_RH_Default_Temp_3 | 0.10744 | 0.085095 | -0.11888 | 0.16009 |
| 7Networks_RH_Default_Temp_4 | 0.13335 | 0.036885 | -0.085202 | 0.3631 |
| 7Networks_RH_Default_Temp_5 | 0.038821 | 0.55844 | -0.023684 | 0.7775 |
| 7Networks_RH_Default_PFCv_1 | 0.03138 | 0.61708 | -0.042085 | 0.64799 |
| 7Networks_RH_Default_PFCm_1 | 0.095385 | 0.18298 | -0.050518 | 0.50949 |
| 7Networks_RH_Default_PFCm_2 | 0.083691 | 0.23389 | 0.0086305 | 0.91555 |
| 7Networks_RH_Default_PFCm_3 | -0.023209 | 0.72355 | 0.1459 | 0.27552 |
| 7Networks_RH_Default_PFCm_4 | 0.10024 | 0.15657 | -0.066918 | 0.3801 |
| 7Networks_RH_Default_PFCm_5 | -0.091333 | 0.15616 | 0.15343 | 0.092462 |
| 7Networks_RH_Default_PFCm_6 | -0.023323 | 0.71201 | 0.094011 | 0.32774 |
| 7Networks_RH_Default_PFCm_7 | 0.01083 | 0.86525 | 0.065998 | 0.52771 |
| 7Networks_RH_Default_PCC_1 | 0.078633 | 0.27349 | -0.068992 | 0.30637 |
| 7Networks_RH_Default_PCC_2 | 0.0191 | 0.77788 | -0.055595 | 0.5432 |
| 7Networks_RH_Default_PCC_3 | 0.033364 | 0.61639 | -0.053764 | 0.60017 |

**Supplementary Table 16. Genetic and environmental correlation between Openness and surface area in HCP.** Genetic correlation between Openness and local surface area calculated using solar 8.4.0. Regions are named according to the Schaefer 200 atlas, based on 7-networks. Here we report environmental correlation (ρ_e_) and genetic correlation (ρ_g_) and the associated p-values.

|  | Average cortical thickness | | Total surface area | |
| --- | --- | --- | --- | --- |
|  | **GSP** | **eNKI** | **GSP** | **eNKI** |
| Agreeableness | 0.67 | -0.22 | 0.35 | 0.47 |
| Conscientiousness | -0.81 | -0.55 | -0.67 (>2.5) | -1.12 (0.56) |
| Extraversion | -0.15 | 1.03 | -0.86 | 0.22 |
| Neuroticism | -0.33 (>3) | -0.16 (>1) | 0.11 (>3) | 0.01 (>1.5) |
| Openness | 1.17 | 0.39 | 0.34 (>3) | 1.55 (0.33) |

**Supplementary Table 17. Replication of association between personality traits and whole brain summaries of surface area and cortical thickness.** Replication analysis in the GSP and eNKI sample, t-values are reported. Bayes Factors (BF) were computed if there was a phenotypic correlation at p<0.05 in the HCP sample. If a BF_01_ is between 0 and 1/3 there is a moderate/strong evidence for H1 (replication), between 1/3 and 1 anecdotal evidence for H1, between 1 and 3 anecdotal evidence for H0 (no replication) and >3 moderate to strong evidence of H0. We underlined replications with a correct sign.

| Personality trait | A | C | E | N | O |
| --- | --- | --- | --- | --- | --- |
| *Agreeableness* | 1 |  |  |  |  |
| *Conscientiousness* | 0.04 (0.13) |  |  |  |  |
| *Extraversion* | 0.38 (0.12) ** | 0.33(0.09) ** |  |  |  |
| *Neuroticism* | -0.14 (0.14) | -0.27(0.10) * | -0.31(0.10) * |  |  |
| *Openness* | 0.17 (0.11) | -0.24 (0.09) * | 0.05(0.09) | -0.01(0.10) |  |

**Supplementary Table 18. Genetic correlation of personality traits.** Genetic correlation across personality traits using solar 8.4.0. ** indicates FDRq<0.05 corrected at p<0.05, controlled for number of analysis (10), whereas * indicates p<0.05.

| Personality trait | A | C | E | N | O |
| --- | --- | --- | --- | --- | --- |
| *Agreeableness* | 1 |  |  |  |  |
| *Conscientiousness* | 0.32 (0.06) ** |  |  |  |  |
| *Extraversion* | 0.26 (0.06) ** | 0.24(0.06) ** |  |  |  |
| *Neuroticism* | -0.41(0.06) ** | -0.51(0.05) ** | -0.39(0.06) ** |  |  |
| *Openness* | 0.06 (0.06) | 0.01 (0.07) | 0.14(0.07) * | 0.02(0.07) |  |

**Supplementary Table 19. Environmental correlation of personality traits.** Environmental correlation across personality traits using solar 8.4.0. ** indicates FDRq<0.05 corrected at p<0.05, controlled for number of analysis (10), whereas * indicates p<0.05.
